# Entangled adaptive landscapes facilitate the evolution of gene regulation by exaptation

**DOI:** 10.1101/2024.11.10.620926

**Authors:** Cauã Antunes Westmann, Leander Goldbach, Andreas Wagner

## Abstract

Exaptation, the co-option of existing traits for new functions, is a central process in Darwinian evolution. However, the molecular changes leading to exaptations remain unclear. We investigated the potential of bacterial transcription factor binding sites (TFBSs) to evolve exaptively for the three global *E. coli* transcription factors (TFs) CRP, Fis, and IHF. Using a massively parallel reporter assay, we mapped three combinatorially complete adaptive landscapes, encompassing all intermediate sequences between three pairs of strong TFBSs for each TF. Our results revealed that these landscapes are smooth and navigable, with a monotonic relationship between mutations and their impact on gene regulation. Starting from a strong TFBS for one of our TFs, Darwinian evolution can create a strong TFBS for another TF through a small number of individually adaptive mutations. Notably, most intermediate genotypes are prone to transcriptional crosstalk – gene regulation mediated by both TFs. Because our landscapes are smooth, Darwinian evolution can also easily create TFBSs that show such crosstalk whenever it is adaptive. We also present evidence of exaptive evolution and crosstalk from an analysis of bacterial genomes. Our study presents the first in vivo evidence that new TFBSs can evolve exaptively through multiple small and adaptive mutational steps. It also highlights the importance of regulatory crosstalk for the diversification of gene regulation.

## Introduction

Exaptation^1^—the evolutionary repurposing of an existing trait for a new function—is a major source of biological innovation. Classic examples span multiple levels of biological organization, from organismal traits such as bird feathers, which likely originated as thermoregulatory structures ^2,3^, to molecular traits ^4,5^ such as vertebrate eye crystallins, which evolved from metabolic enzymes^6,7^. Despite the recognized importance of exaptation, the molecular mechanisms that enable it, and the evolutionary accessibility of genotypes intermediate between ancestral and exapted genotypes, remain poorly understood.

Transcriptional regulation is an important system for studying exaptation because changes in such regulation can alter how genes are connected within a gene regulatory network, and thus enable the emergence of new regulatory functions and adaptations^8–14^. In transcriptional regulation, a transcription factor protein (TF) binds a DNA sequence called a transcription factor binding site (TFBS), which leads to activation or repression of a nearby gene’s transcription^15–17^. TFBSs can evolve de novo from non-regulatory DNA, or through exaptation, in which a pre-existing TFBS originally recognized by one TF becomes repurposed to interact with another TF. Well-documented cases of exaptation include the co-option of a heat shock element as a binding site for the transcription factor Pax6 during the evolution of vertebrate αA-crystallin expression in the eye lens ^6^, as well as the widespread recruitment of transposable element–derived regulatory sequences in mammalian gene regulation ^18–20^.

To understand the origin of such exaptations in molecular detail requires an understanding of the adaptive landscapes on which TFBSs evolve ^21–23^. An adaptive landscape is an analogue to a physical landscape, in which individual locations correspond to a genotype, such as the DNA sequence of a TFBS ^24–27^. The elevation at each location corresponds to a quantitative phenotype, such as the ability of a TFBS to regulate gene expression, which depends on how strongly the TF binds the TFBS ^24–27^. The evolutionary process by which a strong ancestral TFBS for some TF (TF1) becomes exapted as a strong TFBS for another TF (TF2) usually requires an evolutionary path that involves multiple nucleotide changes. We call an adaptive landscape with evolutionary paths that correspond to transitions between TFBSs (or other traits) an *exaptation landscape*. One fundamental question about such a landscape is whether it harbors evolutionary paths towards an exapted trait (TFBS) that are accessible to natural selection. These are paths in which each mutational step is favored by natural selection, for example because it increases the strength of gene regulation by a new TF.

Although most studies of adaptive landscapes assume that one genotype has only one phenotype^24,28–30^, biological reality is more complex. Viral genomes, for example, commonly contain overlapping genes, “entangling” the evolutionary trajectories of distinct proteins through shared nucleotide sequences^31^. Similarly, TFBSs often overlap in both prokaryotic ^32–35^ and eukaryotic genomes^36,37^. Such overlaps can give rise to *transcriptional crosstalk*, in which a single TFBS can mediate gene regulation by more than one TF ^38–41^. A single mutation can then also alter more than one phenotype. In consequence, a single regulatory sequence may be subject to competing selective pressures, which can create regulatory conflicts that must be resolved during adaptive evolution^42^. We refer to an entire adaptive landscape in which one genotype has two or more phenotypes as an *entangled* landscape.

Transcriptional crosstalk is commonly viewed as a byproduct of a regulator’s imperfect specificity that imposes physical constraints on reliable gene regulation^40^. Theoretical estimates suggest that transcriptional crosstalk may affect approximately 1–10% of prokaryotic genes and 10–50% of eukaryotic genes^40^. Crosstalk can also impose direct fitness costs. Pertinent evidence comes from experiments in which transcription factors from bacteriophages were expressed in *E. coli* and *Salmonella enterica*. When five such TFs were introduced into these bacteria, and their crosstalk was quantified by the number of genomic sites they could bind, TFs with more crosstalk caused stronger growth defects ^43^.

Yet crosstalk need not always be deleterious. Theory predicts that it can be adaptive in changing environments, in large regulatory networks, and when selection against weak misexpression is limited ^41^. One experimental example comes from *Pseudomonas fluorescens*, where loss of the master flagellar regulator FleQ was rescued by the unrelated regulator NtrC. After mutations increased NtrC activity and expression, its weak binding to flagellar promoters became strong enough to restore flagellar gene expression and motility. Subsequent mutations further refined this new regulatory interaction^44,45^. In sum, crosstalk can hinder or help adaptive evolution, and the circumstances in which it does either are poorly understood.

While eukaryotic regulatory DNA regions typically rely on the combinatorial action of multiple TFBSs embedded within large, modular enhancers ^34,46,47^, prokaryotic regulatory regions are shorter and less complex, often comprising only a few TFBSs that function independently of each other^17,34,47^. That individual TFBSs in prokaryotes can act as discrete functional units ^48–50^, makes them well-suited for studying their evolutionary dynamics, and their potential for exaptive evolution.

Here, we experimentally map the exaptation landscape of TFBSs for three major prokaryotic TFs in *Escherichia coli* ^15,51,52^. These are CRP^53^ (cAMP receptor protein or catabolite activator protein), Fis^54^ (Factor for Inversion Stimulation), and IHF^15,51,52^ (Integration Host Factor). We chose these TFs, because they have independent evolutionary origins^55^, perform different biological functions, and are important global regulators that control the expression of hundreds of genes across the genome ^15,51,56^. CRP regulates carbon source utilization in the absence of glucose^57^. Fis primarily acts in fast-growing cells and can serve as either a repressor or activator depending on sequence context^58,59^. In contrast, IHF mainly regulates genes during the transition to stationary phase^60–62^. In addition to their role in transcriptional regulation, Fis and IHF are nucleoid-associated proteins that also mediate chromosome condensation^62–65^.

Despite their distinct evolutionary origins and physiological functions, these TFs share a key feature: promiscuous DNA binding. They recognize a broad array of genomic sequences and are frequently found acting in combination within complex promoter architectures, with their binding sites arranged in tandem or overlapping ^34,35,66–68^. This co-occurrence leads us to ask whether a binding site of one of these TFs could originate from a binding site of another by a process of exaptation that consists of one or more individually adaptive mutational steps.

More specifically, we ask for all three pairs of these TFs (CRP-Fis, CRP-IHF, Fis-IHF) whether it is possible for a strong “wild-type” TFBS (WT^TF^) recognized by one TF to evolve into a strong TFBS for another TF through evolutionarily accessible mutational paths, i.e., paths on which each step increases regulation by the new TF. Using a combinatorially complete library^69^ comprising all possible mutant intermediates between wild-type binding site pairs, we measured the strength of gene regulation conferred by each intermediate through a green fluorescent protein (GFP) reporter system.

We show that all three TF exaptation landscapes are smooth. Each landscape has two global peaks that correspond to high-affinity DNA binding sites mediating strong regulation by each of the two TFs. All intermediate TFBSs between the wild-type sequences for each pair of TFs are subject to crosstalk. That is, they can mediate regulation by both TFs to some extent. The entangled landscape of each TF pair allows TFBS exaptation through multiple steps that are all individually adaptive.

## Results

### Mapping *in vivo* exaptation landscapes

To quantify TFBS regulation strength, we re-engineered a plasmid from a previous study^70^ (**Figure 1a, Supplementary Figures S1–S2, Supplementary Methods 2–3**). This plasmid allows us to control expression of any one TF through the addition of an inducer (anhydrotetracycline, Atc), and it allows us to measure the TF’s ability to recognize and regulate a given TFBS by monitoring the expression of a downstream GFP reporter (**Figures 1b–c**). We refer to this measurable output as *regulation strength*. While TFBSs can naturally occur in various positions relative to promoters^34^—both upstream and downstream—our expression system standardizes regulation by placing TFBSs between a constitutive promoter and a reporter gene encoding GFP. In this arrangement, TF-TFBS interactions always repress the expression of GFP ^71^ via steric hindrance^16^.

**Figure 1.**
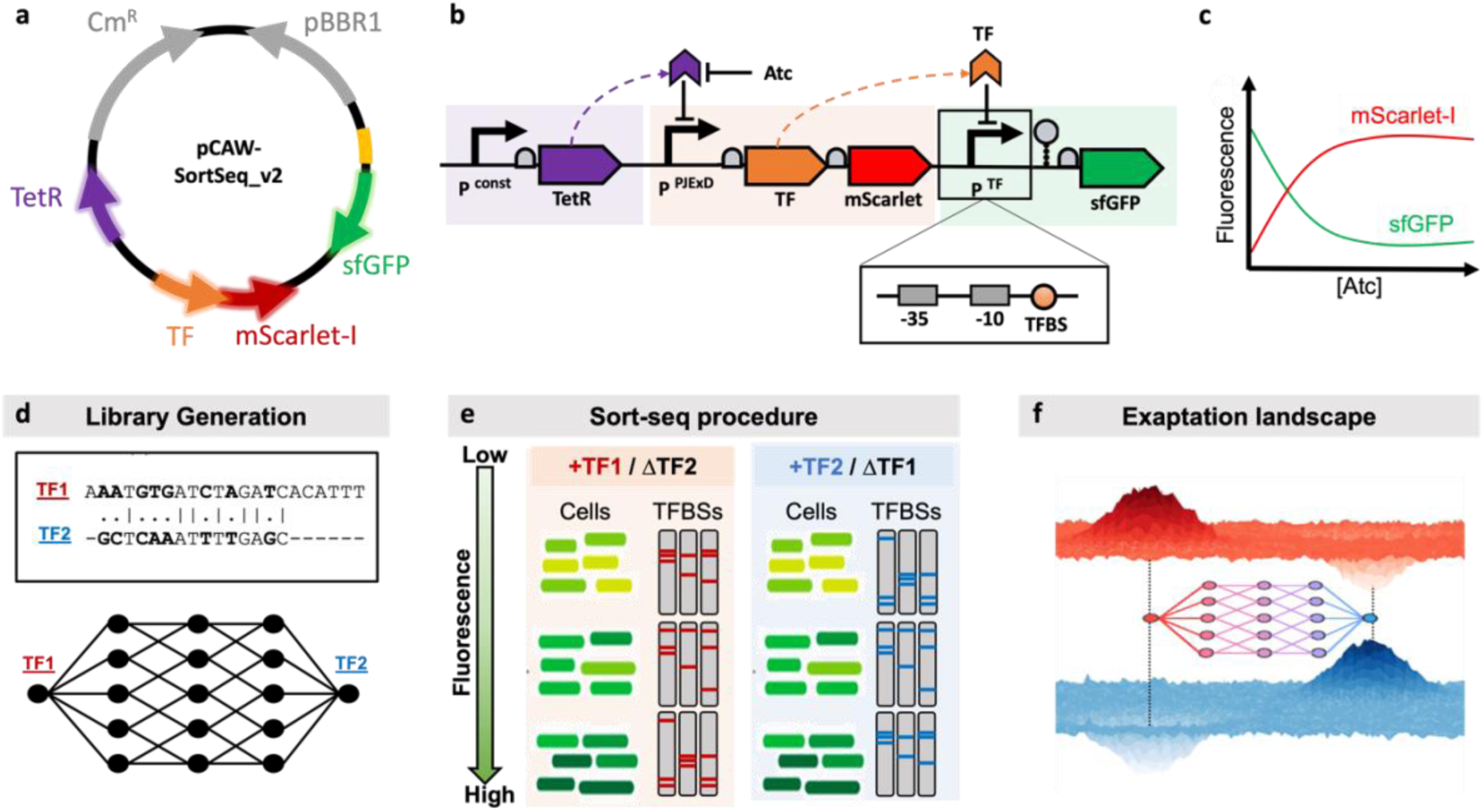
Experimental workflow. **a.** The plasmid system. We redesigned an existing plasmid^70^ (**Supplementary Methods 2-3**) encoding a chloramphenicol resistance gene (CmR, grey, right-oriented arrow) and a broad-host low-copy replication origin (pBBR1, grey, left-oriented arrow) to contain the following regulatory modules. **b.** The plasmid contains three connected modules. The first (purple box) constitutively expresses the *tetR* gene (purple arrow). The TetR regulator controls the second module (red box) through the *plTetO1*^75^ promoter, which consists of a bicistronic operon with the TF of interest (orange arrow) followed by the *mScarlet-I*^76^ reporter gene (red arrow). The third module (green box) allows the measurement of TFBS regulatory effects. TF-TFBS interactions repress the expression of the *sfGFP* ^71^ reporter gene (green arrow) via steric hindrance^16^. The addition of anhydrotetracycline (Atc) alleviates TetR repression. A RiboJ transcriptional insulator ^77,78^ (grey hairpin) prevents unintended 5’-UTR effects from TFBSs. Ribosome binding sites (RBSs) are shown as grey semi-circles. **c. Schematic illustration of reporter dynamics.** Increasing Atc sequesters TetR, allowing the TF and *mScarlet-I* expression (red), which in turn represses *sfGFP* fluorescence (green). We performed all sort-seq measurements at a saturating Atc concentration (100 ng/mL) to ensure maximum TF expression, because the phenotypic output depends on both TFBS sequence and TF concentration. **d. Library generation.** Alignment of two strong-binding (“wild-type”) TFBSs for individual TFs, indicating positions (dots) at which the two aligned sites differ from each other. We created a binding site library by varying the nucleotides at all nucleotide positions that differ between the two sites. The library comprises *2^n^* sequences, where *n* is the number of positions at which the sites differ (See **Supplementary Methods 4, Supplementary Table S3** and **Supplementary Figure S3**). **(e) Sort-seq procedure.** For each TFBS library, we measured regulatory activity under three genetic backgrounds: a TF knockout strain lacking the TF of interest (ΔTF) and strains expressing either TF1 or TF2 (+TF1 or +TF2). We grew cells carrying the reporter plasmid under saturating Atc concentrations, and sorted by sfGFP fluorescence into discrete expression bins using fluorescence-activated cell sorting (FACS). We sorted cells independently for +TF1 and +TF2 conditions, yielding TF-specific measurements of regulatory activity. We identified TFBS sequences present in each bin and quantified their incidence by high-throughput sequencing, enabling reconstruction of regulation strength for every TFBS with respect to each TF. **f. Schematic representation of an exaptation landscape.** For each library, each TFBS is characterized by two regulation strengths—one for each TF—defining a dual adaptive landscape. The top and bottom surfaces represent schematic regulation strength landscapes for TF1 (top, red) and TF2 (bottom, blue). Individual locations (x and y axis) correspond to TFBS genotypes and the elevation at each location (z-axis) corresponds to the regulatory strength of each genotype. The central network represents this dual landscape, where nodes correspond to TFBSs connected by single-nucleotide mutations. Node colors indicate regulatory phenotypes, transitioning from red (stronger TF1 regulation) to blue (stronger TF2 regulation).

We started our investigation of three exaptation landscapes for *E. coli* TF pairs by identifying a strong TFBS for each TF that we refer to as the wild-type (WT_CRP_, WT_Fis_, WT_IHF_), and by aligning these TFBSs to identify the minimal number of nucleotide changes required to convert them into each other (**Figure 1d, top panel**, **Supplementary Figure S3**, **Supplementary Tables S2-S3**). Based on these alignments, we synthesized DNA libraries covering all sequences along these mutational paths (**Figure 1d, bottom panel**). We then measured the regulation strength of each sequence for both TFs using a “sort-seq” reporter assay^26,72–74^, in which cells carrying different TFBS variants are sorted by GFP fluorescence, the underlying sequences are identified by high-throughput sequencing, and the regulation strength is quantified (**Figure 1e**, **Supplementary Methods 7.2**). With this assay, we characterized each sequence with two regulation strength values. The resulting landscapes for CRP-Fis, CRP-IHF, and Fis-IHF pairs consisted of 256, 128, and 128 distinct TFBSs, respectively (**Figure 1f**, **Supplementary Figure S3**, **Supplementary Table S3**). Importantly, each TFBS library occupies a distinct sequence space defined by the specific pairwise alignment of two wild-type TFBSs. Consequently, the CRP–Fis, CRP–IHF, and Fis–IHF libraries contain non-overlapping sets of sequences and are not interchangeable. No single TFBS is shared across all three libraries.

### TFBS libraries for pairs of global regulators vary widely in their regulation strengths

To isolate the regulatory effects of individual TFs in our libraries, we performed all measurements in *Escherichia coli* strains carrying single deletions of the corresponding chromosomal TF gene (*Δcrp*, *Δfis*, or *Δihf*; **Supplementary Table S4**). This design ensured that TF activity arose exclusively from the plasmid-encoded TF and was not confounded by endogenous expression. We did not use double-deletion strains (e.g., *Δcrp Δfis* for CRP–Fis experiments), because simultaneous removal of two global regulators leads to severe pleiotropic growth defects that preclude reliable quantitative measurements under matched conditions^32,33^. Although the single-deletion strains grew modestly more slowly than the wild type, all strains reached comparable cell densities in late exponential to early stationary phase, the growth window in which we performed all measurements (**Supplementary Figure S4**). We validated our experimental system by quantifying the regulation strength of each wild-type TFBS (WT_CRP_, WT_Fis_, WT_IHF_; **Supplementary Table S2**) by its cognate TFs. We confirmed strong regulation and negligible crosstalk between the wild-type TFBSs (**Supplementary Figure S5**). We then characterized each TF-pair library using two complementary plasmid–strain combinations. For example, we assayed regulation strength in the CRP–Fis library by measuring CRP-dependent regulation in a Δfis background (pCRP–Δfis) and Fis-dependent regulation in a Δcrp background (pFis–Δcrp; **Supplementary Figure S13a**). We applied the same experimental design to the CRP–IHF and Fis–IHF libraries (**Supplementary Figure S13b-c; Methods**). We quantified regulation strengths for all TFBS variants using sort-seq (**Methods**; **Supplementary Methods 5**; **Supplementary Figures S6–S8**) by averaging sequencing read counts across fluorescence bins. For each TF, we normalized regulation strengths to the cognate wild-type TFBS, which represents the maximal regulatory output we observed for that TF (**Supplementary Methods 7.2**).

We next confirmed that our regulation strength measurements are reproducible, with Pearson correlation coefficients among technical replicates ranging from 0.913 to 0.988 (**Supplementary Figures S10–S12**). To further validate the sort-seq measurements, we selected sequences spanning a wide range of regulation strengths and measured their GFP output individually by a plate reader. These more direct measurements agreed well with the sort-seq estimates (**Figure 2d–f**), with negative correlations as expected for a repression-based reporter, in which stronger regulation corresponds to lower GFP fluorescence (**Figure 1c**). Because baseline expression and wild-type repression are similar across transcription factors and strain backgrounds, WT-normalized regulation strengths is directly comparable between TFs. Also, we performed all measurements at a fixed, near-maximal inducer concentration, such that differences in regulation strength among TFBS variants primarily reflect TFBS sequence rather than transcription factor abundance (**Supplementary Figures S5 and S9; Supplementary Methods 7.3**). Regulation strength varied broadly across all three libraries (**Figures 2a–c**).

**Figure 2.**
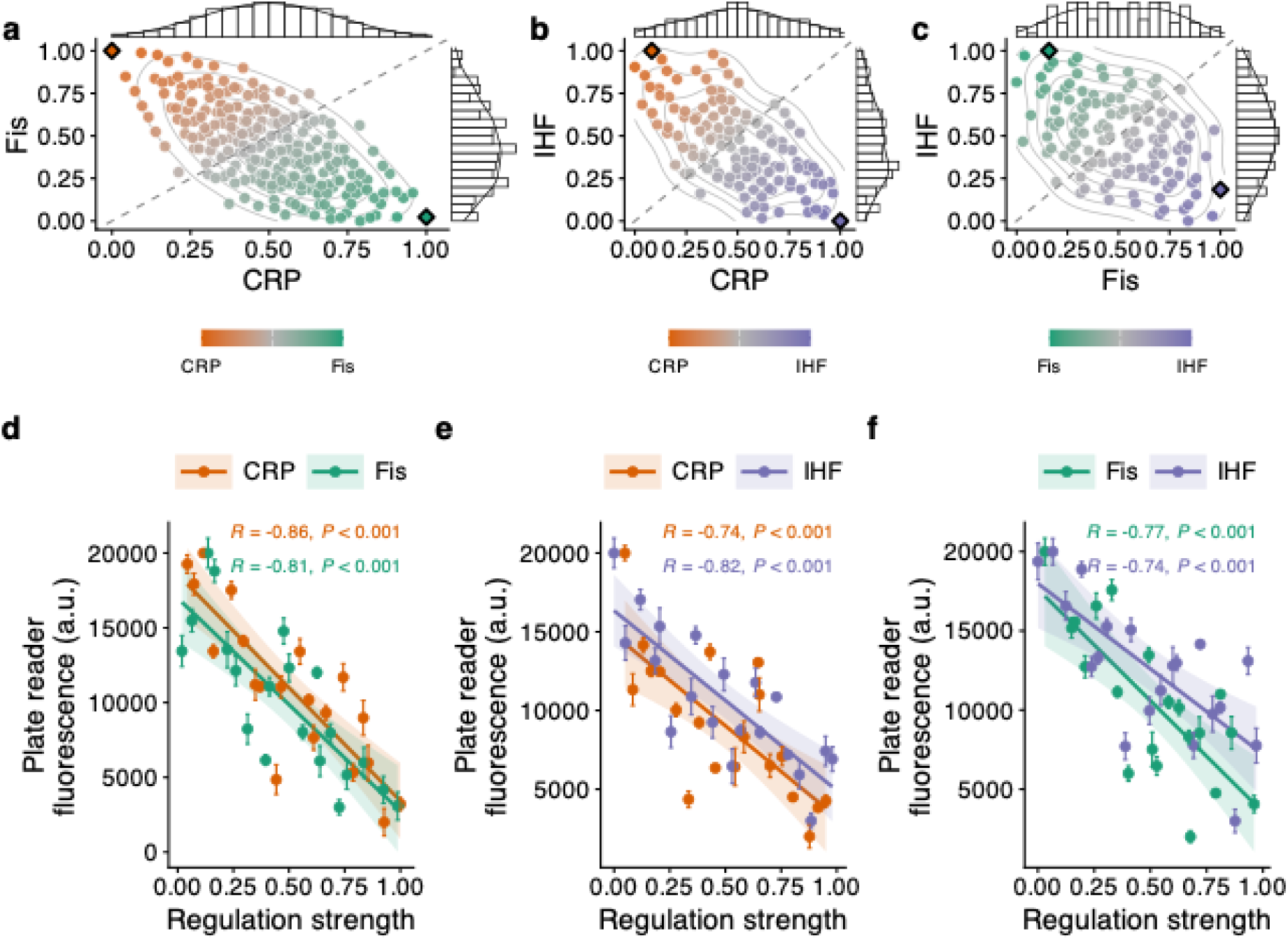
Characterization of regulation strengths for individual TFs. **a-c.** Distribution of regulation strengths for three TFBS libraries. Each point represents a single TFBS sequence. The horizontal and vertical axes indicate regulation strength for the transcription factors labeled on the axes. Point color reflects the relative strength of regulation by each TF (CRP, orange; Fis, green; IHF, purple), with intermediate colors indicating similarly strong regulation by both TFs. Diamond markers denote the cognate wild-type TFBSs, which define regulation-strength maxima for each TF. Marginal histograms (top and right) show the distributions of regulation strength for each TF. The dashed diagonal line indicates equal regulation strength for both TFs, and gray contour lines show kernel density estimates of regulation strength. **a. CRP–Fis library (N = 256). b. CRP–IHF library (N = 128). c. Fis–IHF library (N = 128). d–f. Experimental validation of sort-seq–derived regulation strengths using plate reader fluorescence measurements.** For each TFBS library, we selected 20 sequences spanning a broad range of regulation strengths (**Supplementary Tables S6–S8**). The horizontal axis shows regulation strength inferred from sort-seq, and the vertical axis shows GFP reporter fluorescence measured by plate reader (arbitrary units, a.u.). Points represent the mean of three independent biological replicates measured after eight hours of growth. Error bars indicate one standard deviation. Solid lines show linear regressions, with shaded regions indicating 95% confidence intervals. Pearson correlation coefficients R (*N* = 20) and corresponding two-sided *P*-values are shown. *P*-values are based on the t-statistic test of the null hypothesis that plate reader fluorescence values and regulation strength are not associated. Negative correlations indicate that higher regulation strength corresponds to lower reporter fluorescence, consistent with the repression-based transcriptional reporter system. **d. CRP–Fis library.** CRP (orange, *R* = −0.86, *P* = 0.004) and Fis (green, *R* = −0.81, *P* = 0.009). **e. CRP–IHF library**. CRP (orange, *R* = −0.75, *P* = 0.007) and IHF (purple, *R* = −0.82, *P* = 0.015). **f. Fis–IHF library**. Fis (green, *R* = −0.76, *P* = 0.013) and IHF (purple, *R* = −0.74, *P* = 0.022).

### Exaptation landscapes are smooth and reveal abundant crosstalk

Next we mapped the three exaptation landscapes as networks of genotypes (TFBSs), in which two genotypes are connected if they differ by a single nucleotide (**Figure 3**). The topography of this landscape reveals whether exaptation could occur through natural selection alone.

**Figure 3.**
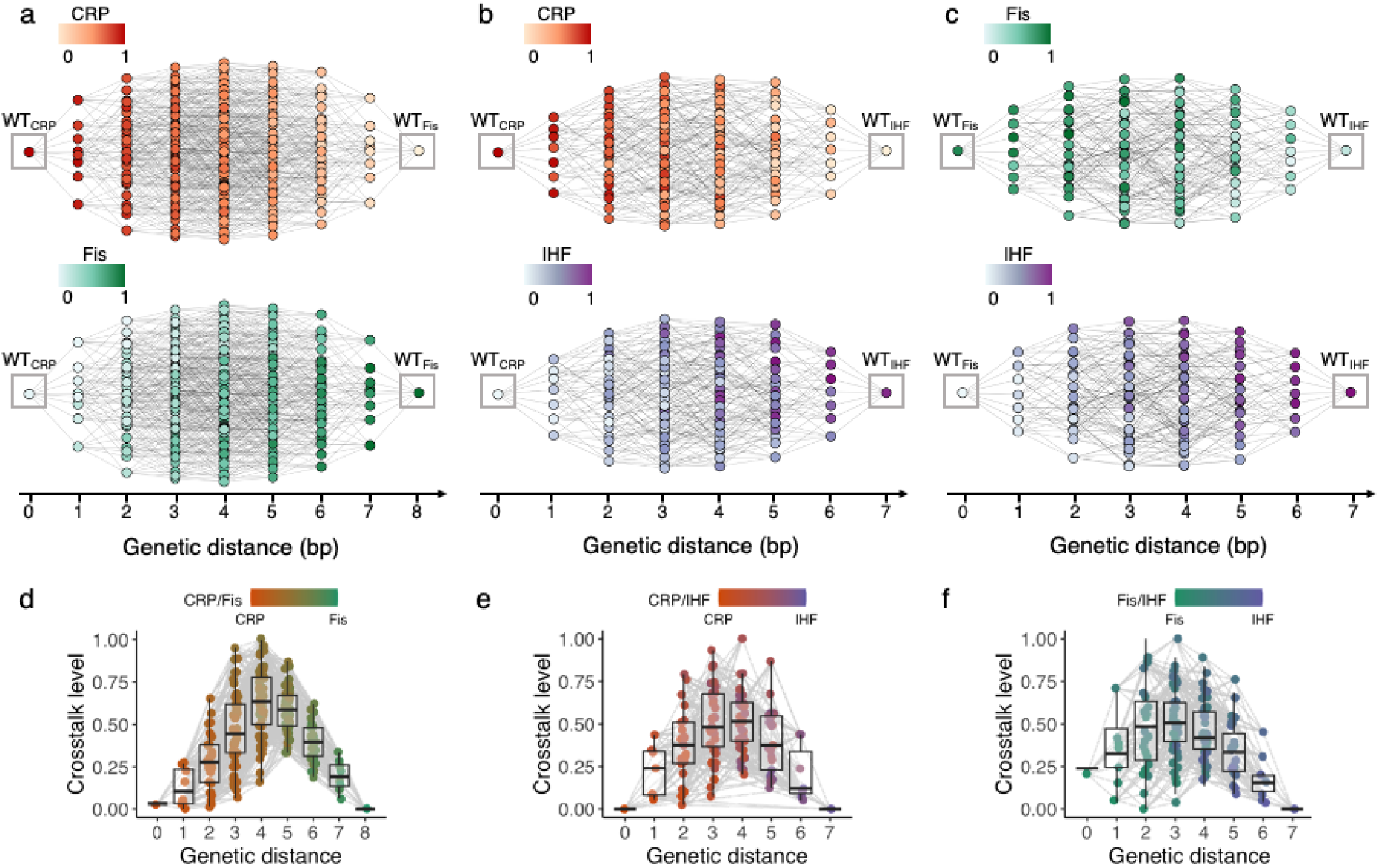
Exaptation landscapes are smooth, with many intermediates subject to transcriptional crosstalk. **a-c.** Exaptation landscapes as networks. Nodes (circles) represent TFBS variants connected by single-nucleotide mutations, with the x-axis showing genetic distance from one of the wild-type sequences (grey squares, x = 0), which are peaks in their respective landscapes. Colors indicate regulation strengths normalized by the wild-type value, with light and dark colors denoting weak and strong regulation strengths, respectively. **a. CRP-Fis landscape.** Regulation strengths for CRP (red heatmap, top) and Fis (green heatmap, bottom). N = 256 TFBSs**. b. CRP-IHF landscape**. Regulation strengths for CRP (red heatmap, top) and IHF (purple heatmap, bottom). N = 128 TFBSs. **c. Fis-IHF landscape**. Regulation strengths for Fis (green heatmap, top) and IHF (purple heatmap, bottom). N = 128 TFBSs. **d-f. Crosstalk levels.** For each landscape, crosstalk (y-axis) is plotted against genetic distance from the wild-type TFBS (x-axis). Heatmap colors reflect the regulation strength ratio for each TFBS. That is, sequences binding predominantly to CRP, Fis or IHF are displayed in red, green, and purple, respectively. Intermediate colors indicate binding at intermediate strength to two TFs. Boxplots display the distribution of crosstalk values (**Methods**). Each box covers the range between the first and third quartiles (interquartile range, IQR). The horizontal line within the box represents the median value, and whiskers span 1.5 times the IQR. Crosstalk values are defined as the geometric mean of the two wild-type regulation strengths, yielding a scale that ranges from 0 (no crosstalk) to 1 (equal maximal regulation by both TFs). **d. Crosstalk in the CRP-Fis landscape. e. Crosstalk in the CRP-IHF landscape. f. Crosstalk in the Fis-IHF landscape.**

The first pertinent line of evidence comes from a gradual change in regulation strength as the distance of a TFBS from the wild-type increases (**Figures 3a-c**). While wild-type TFBSs are primarily regulated by their cognate TF, all intermediate TFBSs show cross-talk, that is, they are also regulated by the non-cognate TF. We quantified crosstalk as the geometric mean of regulation strengths for TF1 and TF2 (**Figures 3d-f**, **Methods**). Its ubiquity implies that single mutations in a TFBS partially preserve regulation and can therefore be favored when selection acts to promote regulation by the second TF (**Figures 3d-f**). Nucleotide frequency analyses identify specific positions that contribute disproportionately to changes in regulation strength (**Supplementary Fig. S14**).

Beyond the existence of selectively viable intermediates, exaptation also depends on the global structure of the landscape. We therefore examined the distribution of regulatory peaks, defined as TFBSs with higher regulation strength than all single-mutant neighbors. Each landscape contains only two peaks, corresponding to the wild-type TFBSs for TF1 and TF2 (**Figures 3a– c**). The absence of additional local optima indicates globally smooth landscapes. Evolving populations cannot become trapped on suboptimal peaks^24,79^.

While the gradual changes in regulation strength and the absence of suboptimal peaks indicate that these landscapes are smooth, smoothness alone does not guarantee that evolution can proceed monotonically toward alternative regulatory optima. To assess how landscape structure constrains the set of possible evolutionary trajectories, we quantified the fraction of accessible shortest mutational paths (among all possible paths) connecting the TF1 and TF2 wild-type TFBSs. A path is shortest if it comprises no more mutational steps than the number of nucleotide differences between the wild-type TFBSs of TF1 and TF2. It is accessible if regulation strength increases monotonically at every mutational step along the path.

Accessibility is high in all three landscapes: regardless of orientation and direction of evolution, at least 65% of shortest paths are accessible (**Supplementary Figure S15**). The inaccessible paths reflect instances of reciprocal sign epistasis^80^, a form of non-additive interaction among mutations in which individual mutations reduce regulation strength, whereas their combination increases it. Such epistasis is rare in our data. It affects only 4%, 11%, and 0.3% of double mutants in the CRP–Fis, the CRP–IHF, and the Fis-IHF landscape (**Supplementary Tables S9–S11; Supplementary Fig. S15**). Thus, these landscapes are not only smooth, they would also be highly “navigable” by adaptive Darwinian evolution.

### *In silico* analysis suggests exaptation between two closely related bacterial species

To study the exaptive evolution of TFBSs, one would ideally reconstruct their evolutionary history with phylogenetic methods, as has been done for eukaryotic TFBSs ^81–84^. However, this is challenging for bacterial TFBSs, because their short length, rapid turnover, and frequent reshaping by recombination and horizontal gene transfer make their histories difficult to infer reliably ^85–88^. We therefore used a comparative genomic approach and analyzed regulatory sequences from the closely related enteric bacteria *Escherichia coli K-12 MG1655* and *Salmonella enterica serovar Typhimurium LT2*, which diverged approximately 100 million years ago ^89^ (**Supplementary Methods 7.7**). Because *E. coli* global regulators also exist in *S. Typhimurium*, even though the specific composition of their regulons has diverged^90,91^, we hypothesized that some of their TFBSs might have been repurposed during evolution. We thus looked for genomic signatures of such exaptation.

To this end, we identified 105 orthologous gene pairs with conserved operon structure and extracted their upstream intergenic regions. We then aligned these regulatory regions and searched them for candidate binding sites for eight global regulators^51^ (CRP, Fis, IHF, ArcA, FNR, Cra, Lrp, and Fur). To identify candidate binding sites for these regulators, we used position-weight matrices (PWMs) – computational tools to identify TFBS motifs for specific TFs^92,93^ (**Supplementary Methods 7.8**). These PWMs are derived from experimentally validated sequences deposited in the database RegulonDB^55^ (370 sequences for CRP, 267 for Fis, and 119 for IHF). We classified a site as a candidate exaptation event when a binding site for one regulator in one species aligned with a binding site for a different regulator in the orthologous region of the other species. By contrast, we classified sites present in only one species as lineage-specific site gains or losses. Across the 105 aligned (intergenic regions, we detected an average of 3.2 ± 0.3 differences in predicted TFBS identity per regulatory region, and 34% of these matched our criterion for candidate exaptation events (**Supplementary Figure S16; Supplementary Methods 7.7**). Thus, among TFBS differences between these two genomes, about one third are consistent with exaptive turnover. We note that this criterion cannot unambiguously distinguish exaptation from de novo evolution: a TFBS that arose independently in one species could, by chance, align with a different TFBS in the other. However, theoretical work predicts that exapted sites should retain greater sequence similarity to ancestral binding motifs than sites evolved de novo, because they originate from pre-existing functional sequences ^94^. Our approach captures this signature: by requiring positional overlap between a TFBS for one TF in *E. coli* and a TFBS for a different TF in *S. typhimurium*, we select for cases where the underlying sequence retains sufficient compatibility with both motifs consistent with the enhanced cross-motif similarity expected under exaptation (**Supplementary Methods 7.7)**.

### Rapid exaptation when Darwinian evolution favors strong binding by one TF

Motivated by the smoothness of exaptation landscapes (**Figure 3a-c**) and by the potential for TFBS exaptations in bacterial genomes (**Supplementary Figure S16**), we also examined how populations would evolve on our experimentally mapped landscapes. To this end, we simulated population evolution in the strong selection weak mutation (SSWM) regime ^95–98^ (**Supplementary Methods 8**), which is relevant for the small mutational targets of our TFBSs (7-8 base pairs), and for organisms like *E. coli* with large population sizes (>10⁸ individuals^99^). In this regime beneficial mutations typically fix before new mutations arise, resulting in mostly monomorphic populations that adapt via successive fixation events of individual mutations ^95-98^. In other words, adaptive evolution can be modeled as an adaptive walk, in which each mutational step is taken with a fixation probability that has been derived by Kimura ^100^. For each TF1-TF2 landscape, we simulated 10⁵ adaptive walks starting from the 10 strongest TFBSs for TF1, with fitness defined as the regulation strength mediated by TF2 (**Supplementary Methods 8**). We also simulated walks in the opposite direction, starting from TF2’s strongest binding sites. We incorporated known *E. coli* mutation biases^101,102^ into our simulation model.

In large populations (10^8^ individuals), all simulated walks reached the wild-type TFBS of the target TF, corresponding to maximal regulation strength (regulation strength = 1; **CRP–Fis: Fig. 4a–b; CRP–IHF: Supplementary Fig. S17a–b; Fis–IHF: Supplementary Fig. S18a– b**). Crosstalk typically peaked at intermediate mutational distances and declined as walks approached the target optimum. Adaptive walks were also short, requiring only slightly more steps than the minimum imposed by the sequence differences between the two wild-type TFBSs. In the CRP–Fis landscape, we tested the hypothesis that adaptive path lengths are identical between directions (CRP→Fis vs. Fis→CRP), and found that walks from CRP to Fis were slightly shorter than in the reverse direction (mean ± s.d.: 6.6 ± 0.1 vs. 6.9 ± 0.9 steps; Welch’s t-test, P < 0.001; **Figure 4c–d**, **Supplementary Figures S17e–f** and **S18e–f**). More generally, small deviations in average walk length from the minimal distance reflect the modest incidence of reciprocal sign epistasis in our landscapes ^24,103,104^ (**Supplementary Tables S9–S11**). Taken together, these observations underscore the smoothness of our landscapes. When selection favors stronger binding by an alternative TF, exaptation can proceed rapidly across these landscapes.

**Figure 4.**
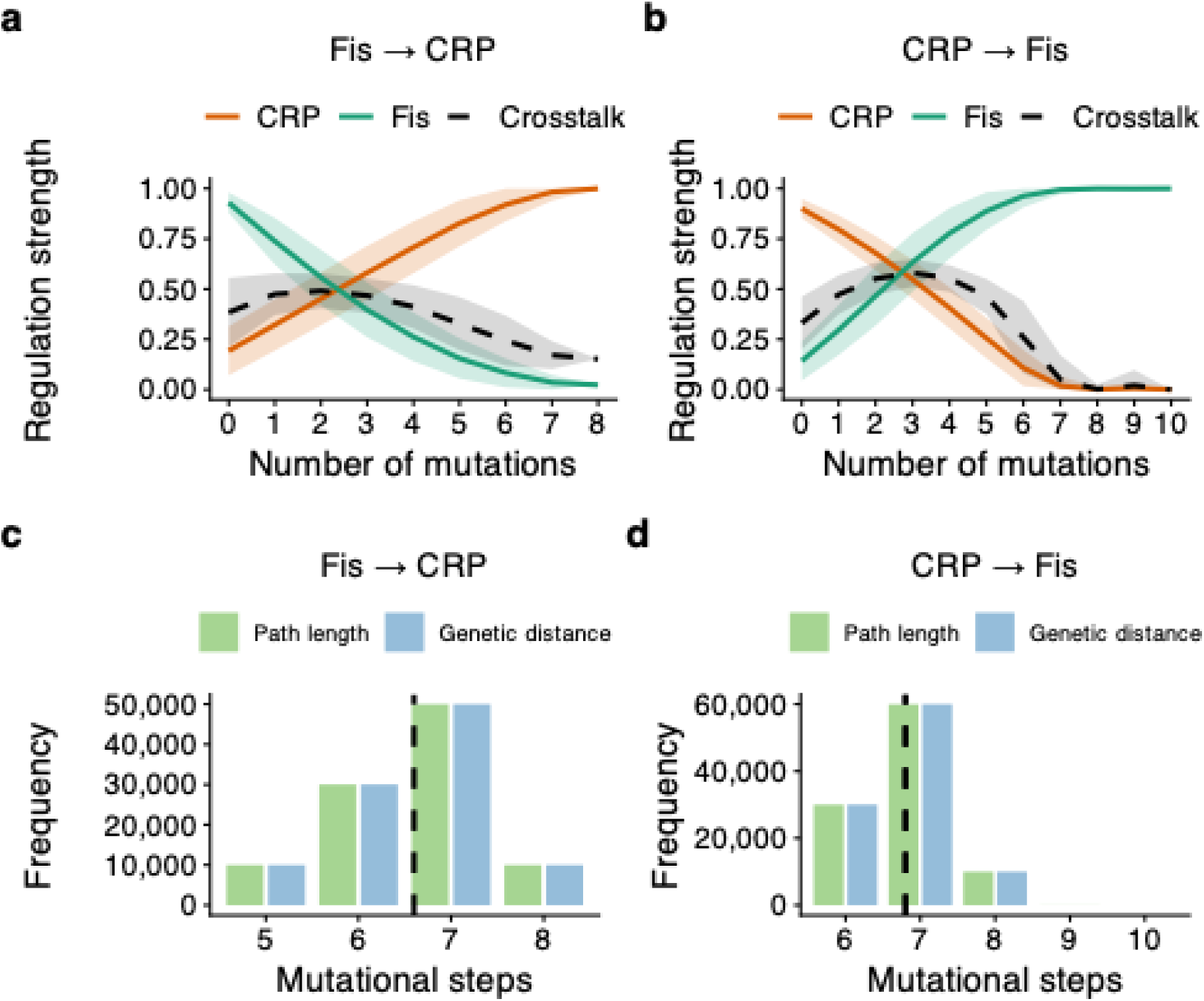
Rapid exaptation on the CRP-Fis landscape. All data are based on a population of 10⁸ individuals and 10⁵ simulated adaptive walks (10⁴ walks for each of the 10 starting genotypes with the highest regulation strength for the opposite TF). **a-b. Adaptive walks readily attain peaks.** Each plot shows how regulation strength (vertical axis) for CRP (orange) and Fis (green) changes as a function of path length traversed (horizontal axis) from the initial sequence to the reached peak during adaptive walks. Crosstalk along the walks is indicated by the black dashed line and is defined as the geometric mean of CRP and Fis regulation strengths. Curves represent averages over all 10^5^ adaptive walks, and shading indicates one standard deviation of regulation strength. **a**. In the CRP landscape, adaptive walks start from the 10 strongest Fis TFBSs and favor strong CRP binding. **b**. In the Fis landscape, adaptive walks start from the 10 strongest CRP TFBSs and favor strong Fis binding. **c-d. Evolutionary paths to a peak are not much longer than the shortest genetic distances.** Grouped bar charts show the frequency distributions of path lengths (green) and genetic distances (blue), displayed side by side for each number of mutational steps (horizontal axes). The black dashed vertical line represents the mean, which is not significantly different between distributions. We used a t-test of the null hypothesis that the means of these distance distributions are statistically indistinguishable. **c. Accessible path lengths and genetic distances to the CRP peak.** Mean ± s.d.: 6.6 ± 0.1. Welch Two Sample t-test: t = 31.973, *P* = 0.7, N = 10⁵ for both distributions. **d. Accessible path lengths and genetic distances to the Fis peak.** Mean ± s.d.: 6.9 ± 0.9. Welch Two Sample t-test: t = 35.113, *P* = 0.82, N = 10⁵ for both distributions.

While accessibility determines which evolutionary paths are possible (**Supplementary Figure S15**), it does not determine which of these paths are realized. To examine whether contingency may shape evolutionary trajectories within the set of accessible paths, we next analyzed the diversity and structure of adaptive walks. Multiple paths to the same peak exist, highlighting a potential for contingency, the dependence of evolutionary processes on chance events ^105–107^ (**Supplementary Figures S19-S20**). Conversely, we also observed trajectory biases, with adaptive walks preferentially passing through TFBSs with stronger rather than weaker regulation (**Supplementary Figure S22**).

In small populations (10²), where drift is strong (**Supplementary Methods 8)**, final fitness was only modestly below the global maximum (by ∼1%, p < 2.2 × 10⁻¹⁶, One-Sample t-test), and walks were marginally longer (10%, p < 2.2 × 10⁻¹⁶, Welch t-test, **Supplementary Figures S22a-l**), indicating that stochastic effects slightly impede but do not prevent exaptation. Finally, we studied the dynamics of populations outside the SSWM regime. Sufficiently large populations or populations at high mutation rates can experience clonal interference, in which multiple beneficial mutations compete for fixation^108–110^ (**Supplementary Methods 8)**. Deviations from the SSWM regime do not change the ability of populations to evolve exaptations (**Supplementary Figures S23-S25**). Overall, our smooth, single-peaked landscapes facilitate rapid exaptation by selection, regardless of drift or clonal interference.

### Transcriptional crosstalk is widespread within the *E. coli* genome

Motivated by these findings, and by the fact that CRP, Fis, and IHF often act together at *E. coli* promoters^34,66–68^, we asked how much transcriptional crosstalk these three regulators might show across the *E. coli* genome. To do so, we used the same PWM computational tools as before ^92,93^. Because PWMs summarize the DNA sequence preferences of each TF, they can provide a score estimating how strongly a TF may bind a given site ^92,93^ (see **Supplementary Methods 7.8**). We applied these models to 756 experimentally validated TFBSs from the *E. coli* genome ^55^. Because these predicted binding strengths correlate with regulatory activity in our TFBS libraries (mean Pearson *R*=0.56±0.1, **Supplementary Methods 7.8**, **Supplementary Figure S26**), they provide a useful approximation of each TF’s potential to recognize genomic sites beyond its annotated targets.

We considered a site to have crosstalk potential when it is predicted to be recognized by two different TFs. This analysis revealed extensive potential crosstalk in the *E. coli* genome (**Figure 5a; Supplementary Figure S29**, **Supplementary Methods 7.8**). For example, it predicts that 44% of the 756 genomic TFBSs are recognized strongly by both CRP and Fis, whereas 8% and 9% are recognized strongly by both CRP and IHF, and by both Fis and IHF, respectively (**Figure 5b–d**). Thus, while crosstalk potential varies among TF pairs, it is especially pronounced between CRP and Fis.

**Figure 5.**
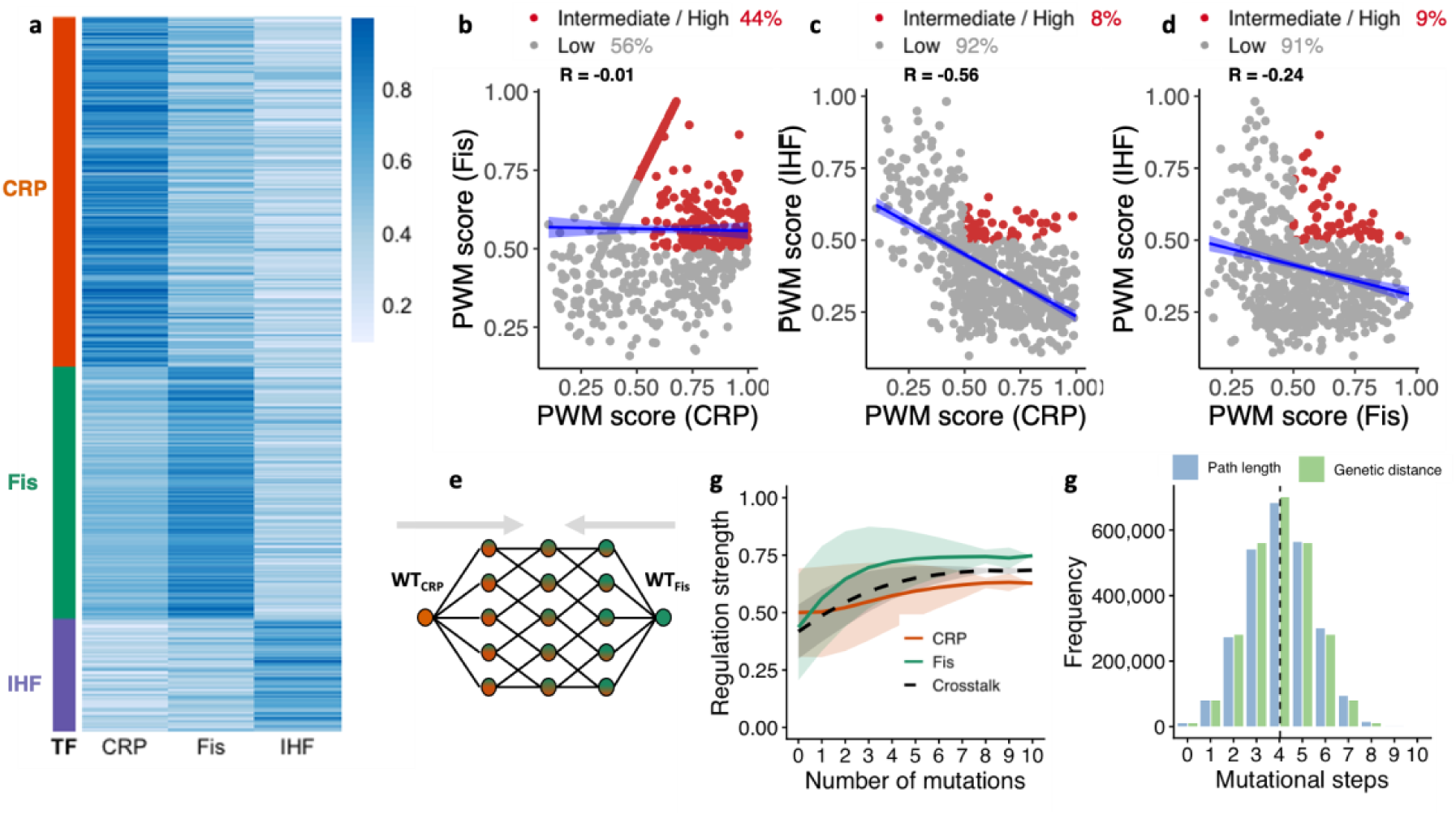
Transcriptional crosstalk and its evolutionary dynamics. a. PWM scores for genomic TFBSs from *E. coli* suggest ample crosstalk among CRP, Fis, and IHF. The heatmap shows 756 experimentally validated TFBS sequences (rows) from RegulonDB ^55^, ordered by their annotated cognate TF, i.e., 370 TFBSs for CRP (orange), 267 for Fis (green), and 119 for IHF (purple). Columns show the (0-1) normalized PWM score of each sequence against each of the three TF PWMs. If TFBSs were strictly specific for their cognate TF, binding sites in each of the three groups would score high only in its cognate column; instead, many CRP and Fis sites score highly on each other’s PWM, indicating widespread predicted crosstalk between these two TFs. IHF sites show lower predicted crosstalk with CRP and Fis, consistent with their more distinct binding motif (see **Supplementary Figures S27-S28**). Hierarchical clustering of this dataset is shown in **Supplementary Figure S29**. **b-d. Predicted binding strength associations.** Scatter plots show the relationships between predicted binding strengths for TF pairs. Red circles indicate TFBSs with high crosstalk (average score > 0.5), while gray circles denote low crosstalk. Percentages of TFBSs in each category and Pearson correlation coefficients (R) are shown. The blue line is a linear regression line (blue shading: 95% confidence interval). **b. CRP and Fis.** No significant association is observed (R = −0.01, *P* = 0.71). **c. CRP and IHF.** A negative correlation is detected (R = −0.56, p < 2 × 10⁻⁶). **d. Fis and IHF.** Another negative correlation is present (R = −0.24, p < 2 × 10⁻⁶). **e-g. Rapid evolution of crosstalk in the CRP-Fis landscape. e. Schematic of evolution towards TFBSs with high crosstalk.** A schematic representation of the CRP–Fis landscape as a network illustrates how adaptive walks in genotype space preferentially (grey arrows) traverse genotypes with increasing binding to both transcription factors, converging on a peak of elevated crosstalk (crosstalk peak). **f. All adaptive walks rapidly attain the crosstalk peak.** Simulated adaptive walks (10⁴ walks starting from each of 256 genotypes) frequently reach a crosstalk adaptive peak TFBSs that bind CRP (orange) and Fis (green) with intermediate strengths. Regulation strength is plotted against path length, with shading representing one standard deviation. **g. Evolutionary paths to the crosstalk peak have the same length as minimal genetic distances.** The green and blue vertical lines show the average length of traversed paths during 10^4^ × 256 walks between pairs of non-peaks and peaks, and the respective mutational distances between such pairs, respectively. The means of the distributions are statistically indistinguishable (3.31±1, mean ± s.d., for both distance metrics, Welch Two Sample t-test t = −5.5782, df = 2029535, *P*-value = 0.7, N_1_= 256 x 10^4^, N_2_= 256 x 10^4^).

### Crosstalk can readily evolve when selection favors dual regulation

Finally, we also asked whether crosstalk itself can evolve adaptively when selection favors regulation by both TFs. To find out, we simulated adaptive walks on the same experimentally mapped landscapes, but defined fitness as the sum of regulation strengths for the two TFs (**Supplementary Methods 8**). We found that high-crosstalk genotypes are readily accessible in all three landscapes, regardless of drift or clonal interference (**Figure 5e–g**; **Supplementary Figures S30-S32, Supplementary Methods 8**).

The CRP–Fis and Fis–IHF landscapes each harbor a single optimal TFBS with substantial crosstalk, whereas the CRP–IHF landscape contain two alternative optima, one of which is reached more frequently than the other. Specifically, 87% of walks reached the first peak with equal regulation strength (0.68) for both CRP and IHF, and 13% reached the second peak, with regulation strengths of 0.87 and 0.32 for CRP and IHF, respectively. In sum, dual regulation can readily evolve when selection favors it.

## Discussion

We experimentally mapped exaptation landscapes for three pairs of global *Escherichia coli* regulators (CRP–Fis, CRP–IHF, and Fis–IHF) and found that all three landscapes are smooth. Each landscape has two peaks that correspond to the wild-type TFBS of each regulator, and most mutational paths between these peaks can be traversed through individually adaptive substitutions. Reciprocal sign epistasis, which can create ruggedness in adaptive landscapes^80^, is rare, and does not present substantial barriers to exaptation. These observations indicate that changes in regulatory specificity of our three transcription factors can occur without requiring passage through strongly deleterious intermediates.

This finding has important implications for regulatory evolution. It suggests that new regulatory interactions can arise not only de novo, but also through gradual repurposing of existing binding sites. Intermediate TFBSs retain measurable regulation by both transcription factors, and adaptive-walk simulations show that populations can readily move between alternative regulatory optima. Such exaptation will be most likely when the ancestral role of a regulator becomes dispensable or maladaptive, for example after ecological change, or a loss of its inducing signal. Our observations are also consistent with evolution experiments demonstrating that weak promiscuous binding can serve as a starting point for regulatory rewiring ^44,45,111–113^. In addition, it has been shown that deletion of a global *E.coli* regulator can be compensated by promoter mutations that enable regulation by alternative transcription factors ^114^. Together, these observations support a view in which exaptation in gene regulation often proceeds gradually through functional intermediates rather than through dramatic changes in regulation. It may often rely on such intermediates rather than on entirely new sequences.

Our results also help clarify the role of transcriptional crosstalk in adaptive evolution. Crosstalk is often considered deleterious because non-cognate interactions can disrupt gene regulation^43^. However, it can also facilitate evolutionary innovation by enabling transitions between regulatory states ^44,45,111–113^. Our landscapes reconcile these perspectives. They show that crosstalk is neither inherently harmful nor inherently beneficial, but instead an evolvable property of regulatory systems. Selection can favor low-crosstalk optima, but it can also exploit crosstalk-competent intermediates when they facilitate adaptive change. Consistent with this view, we observe a potential for widespread crosstalk among CRP, Fis, and IHF binding sites across the *E. coli* genome (Figure 5). How frequently natural selection favors or suppresses crosstalk in different ecological contexts remains an open question ^44,45,111–113^.

The smoothness of the TFBS landscapes we mapped contrasts with observations in other molecular systems ^25,115,116^. For example, in proteins ^115^ and catalytic RNAs ^25^, transitions between distinct phenotypes are often constrained by rugged landscapes, narrow regions of high fitness, or intermediates with reduced performance ^115,117^. In catalytic RNA, for example, dual-function intermediates exist but typically exhibit low activity for both functions, and occupy fitness-depleted regions of sequence space ^25^. In contrast, the TFBS landscapes we studied contain extensive sets of intermediates that retain substantial regulatory activity for both transcription factors. These differences likely reflect the molecular nature of regulatory DNA, which may impose fewer structural constraints than proteins or RNA on the ability of a single sequence to have multiple activities.

Regulatory rewiring is widespread across bacterial lineages ^88,118–121^. Consistent with this observation, our comparative analysis of *E. coli* and *S. Typhimurium* identified changes in conserved regulatory regions that suggest widespread TFBS exaptation (**Supplementary Figure S16**). Because TFBS evolution in bacteria is difficult to reconstruct—owing to rapid TFBS turnover, recombination in intergenic regions, and horizontal transfer of regulatory and mobile DNA ^85–88^ —this evidence is necessarily indirect. Nonetheless, the smooth landscapes, and the short accessible mutational routes we observed may help to render exaptation important for regulatory divergence among genomes.

Our results also broaden the scope of regulatory exaptation beyond closely related transcription factors. In homologous transcriptional repressors of bacteriophages, shifts in DNA-binding specificity can be strongly asymmetric – one repressor can evolve binding to the regulatory DNA of the other, whereas the reverse transition is more difficult ^122^. In contrast, the TFs we study are not even homologous, have distinct biological functions and dissimilar binding sites, yet their binding sites are still connected in a smooth exaptation landscape that can be traversed through successive adaptive mutations. This suggests that accessible regulatory exaptation is not restricted to closely related regulators with similar binding preferences.

Several limitations qualify our conclusions. First, we quantified regulation strength rather than organismal fitness directly, although previous work suggests that regulatory strength is associated with fitness for at least some genes ^48,49,123–125^. Second, our assays used a simplified promoter architecture and fixed, near-saturating TF concentrations, whereas native promoters often integrate multiple regulators ^34,68^ and operate across varying TF abundances ^126,127^. Third, our libraries sample sequence space between TFBSs separated by 7–8 substitutions (**Supplementary Figure S3**) and therefore do not capture all possible evolutionary trajectories. Finally, we examined only three TF pairs, all involving global regulators, and it remains to be seen whether similarly smooth landscapes characterize other classes of transcription factors.

In sum, and within these limitations, we have shown that exaptive evolution in gene regulation can be easily accomplished, because the three exaptation landscapes we study are smooth. Examples of regulatory exaptation^7,18,128–130^ and the existence of crosstalk in the wild^111,112^ suggest that this may be a general principle of exaptation in gene regulation. Further corroboration will require studying more transcription factors, measuring fitness and crosstalk directly, exploring a larger sequence space, and doing so in the chromosomal context in which TFs operate. Such work will help us to understand the entangled landscapes in which regulatory evolution unfolds.

## Methods

The Supplementary methods contain extended details of experimental procedures and data analysis.

### Media and reagents

To prepare SOB medium, we mixed 25.5g from the solid medium stock (VWR J906) with 960 ml of purified water and subjected the resulting suspension to autoclaving. To prepare SOC medium, we dissolved 20 ml of 1 M D-glucose (Sigma G8270) and 20 ml of 1 M magnesium sulfate (Sigma 230391) in 960 ml of the pre-prepared SOB solution. For LB medium, we combined 25g of solid medium stock (Sigma-Aldrich L3522) with 1 liter of purified water and then autoclaved it. To prepare M9 minimal medium, we diluted M9 minimal salt sourced from Sigma (M6030) in distilled water as per the manufacturer’s guidelines, autoclaved the solution, and added 0.4% glucose (Sigma G8270), 0.2% casamino acid (Merck Millipore, 2240), 2 mM magnesium sulfate (Sigma 230391), and 0.1 mM calcium chloride (Sigma C7902). Where required, we supplemented growth media with chloramphenicol (50 µg/mL), anhydrotetracycline (100 ng/mL, Cayman-chemicals #10009542), and/or glucose (0.4% w/v final concentration). We prepared anhydrotetracycline by diluting the dried chemical in absolute ethanol, from a stock concentration (1000X) of 100 µg/mL to a working concentration of 100 ng/mL.

### Strains and plasmids

Bacterial strains and plasmids used in this work are listed in **Supplementary Tables S1-S2**. We obtained electrocompetent *E. coli* cells (strain SIG10-MAX^®^) from Sigma Aldrich (CMC0004). The genotype of this strain (**Supplementary Table S4**) is similar to DH5α (Sigma Aldrich commercial information, see **Supplementary Table S4**). The strain is resistant to the antibiotic streptomycin. Due to its high transformation efficiency, we used strain SIG10-MAX^®^ for molecular cloning and library generation.

We amplified plasmid libraries in DH5α, extracted their DNA, and transformed them into the *E. coli* K-12 BW25113 wild-type strain and its derivative mutants to study gene regulation. We obtained the *E. coli* wild type (BW25113) strain and its mutants, which harbor deletions of CRP, IHF, or Fis global regulators from the KEIO collection ^131^, and used them for sort-seq experiments. The design, genetic parts, and assembly of the plasmid vectors we used in this study are available in the Supplementary material. All primers, TFBS sequences/libraries, strains and plasmids are listed in **Supplementary Tables S1-S5**.

### Sort-seq procedure

We performed our sort-seq experiments as previously described^70^. We designed TF binding site libraries in the Snapgene® software (snapgene.com). The libraries were synthesized by IDT (Coralville, USA) as a single-stranded DNA Ultramer® of 140 bp (4 nmol). We resuspended each library in nuclease-free distilled water and serially diluted it to a concentration of 50ng/uL. We PCR-amplified library oligonucleotides, ligated them into the PCR-amplified pCAW-Sort-Seq-V2 plasmid backbone (a derivative of pCAW-Sort-Seq ^70^, a low-copy number, multi-host plasmid with the pBBR1 replication origin) by Gibson assembly, and transformed cells with the resulting library after column-based purification with a commercial electroporation kit (NEB #T1020L). We grew populations of *E. coli* cells hosting each library in LB medium supplemented with 50 µg/mL chloramphenicol to saturation (overnight growth, 16 hours, 200 rpm, 37°C). After overnight growth, we diluted the overnight culture in LB medium supplemented with 50 µg/mL chloramphenicol and anhydrotetracycline (100 ng/mL) in a 1:100 ratio (v/v) and grew the culture for 8 h until it had reached stationary phase. Before sorting, we diluted 50µL of the culture in 1mL of cold Dulbecco’s PBS (Sigma-Aldrich #D8537) in a 15 mL FACS tube. We performed FACS-sorting on a FACS Aria III flow cytometer (BD Biosciences, San Jose, CA). We sorted and binned cells according to their GFP-mediated fluorescence (FITC-H channel, 488nm laser, emission filter 502LP, 530/30). To this end, we recorded the fluorescence of 10^6^ cells and created 4 evenly spaced gates (“bins”) on a binary logarithmic (log_2_) scale spanning the observed range of fluorescence values (**see Methods, Supplementary Figures S6-S8**). The number of cells sorted into each bin corresponded to the fraction of the 10^6^ cells recorded for that bin. Thus, the total number of sorted cells was 10^6^. We performed each sorting in triplicate from the same overnight culture. We isolated plasmids from cells sorted into each bin (Qiagen, Germany) after overnight growth, and used a polymerase chain reaction (PCR) to amplify the TFBS region from each plasmid for Illumina sequencing. The distribution of library variants among the different bins allowed us to map each individual genotype to a measure of how strongly it regulates gene expression. The Supplementary material has additional details on library construction and sort-seq procedures.

### Procedure for dissecting the effects of individual TFs

To reduce the effects of expression variation among cells (also known as extrinsic noise)^132^ we followed common practice^133–135^ and normalized our *gfp* fluorescence values by *mscarlet* fluorescence values after flow-cytometry assays (**see Supplementary Methods 5**). To quantify within each TF-pair library the regulation strength of one transcription factor independently from that of the other, we combined inducible TF-expression plasmids with strain backgrounds that lack one regulator (**Supplementary Figure S5**). This design allowed us to quantify the contribution of one TF at a time while holding the library context constant. Below, we describe the plasmid–strain combinations we used for each TF pair.

### Sorting the CRP-Fis library

#### Characterizing the effect of Fis binding on gene regulation

To study the influence of Fis binding on our CRP-Fis library sequences independent of that of CRP binding, we proceeded as follows: We cloned the CRP-Fis library onto the plasmid pCAW-Sort-Seq-V2-Fis, which expresses the Fis transcription factor upon Atc induction. Next, we introduced this plasmid into the *E. coli* Δ*crp* strain, in which the *crp* gene has been deleted (**Supplementary Table S4**, **Supplementary Figure S13**). We then cultured the transformed bacterial cells overnight in LB medium supplemented with Atc, which induces the expression of plasmid-encoded Fis within the bacterial population. By doing so, we guaranteed complete absence of CRP and that Fis was expressed. We then subjected the CRP-Fis library to the sort-seq procedure under these conditions, allowing us to specifically examine the effects of Fis binding on our library sequences.

#### Characterizing the Effect of CRP Binding

To study the impact of CRP binding on gene regulation in isolation from that of Fis binding, we cloned the same CRP-Fis library on plasmid pCAW-Sort-Seq-V2-CRP, expressing CRP in the Δ*fis* strain. We induced CRP expression with Atc, ensuring that of the two transcription factors, only CRP is present. After overnight growth, we sorted cells to measure the strength of CRP binding on the CRP-Fis library sequences.

#### Sorting the CRP-IHF Library

We adopted the same procedure we just described for the CRP-Fis library, except that we cloned the CRP-IHF library into the plasmid pCAW-Sort-Seq-V2-IHF, which expresses the IHF transcription factor upon Atc induction, and transformed this library into a Δ*crp* strain. Briefly, to study the effect of IHF binding independent of that of CRP binding, we performed sort-seq after library growth in LB supplemented with Atc. To study the effect of CRP binding independent of that of IHF binding, we cloned the CRP-IHF library into the plasmid pCAW-Sort-Seq-V2-CRP, which expresses the CRP transcription factor upon Atc induction, and transformed this library into a Δ*ihf* strain. We performed the sort-seq procedure after library growth in LB supplemented only with Atc.

#### Sorting the Fis-IHF Library

To study the effect of Fis binding in isolation from that of IHF, we cloned the Fis-IHF library into the plasmid pCAW-Sort-Seq-V2-Fis, and transformed the resulting construct into the Δ*ihf* strain. Inducing the expression of Fis with Atc in this strain ensured the expression of Fis in the absence of IHF. To study the effect of IHF binding in isolation from that of Fis-binding, we cloned the same (Fis-IHF) library into the plasmid pCAW-Sort-Seq-V2-IHF, and transformed the resulting plasmid library into the Δ*fis* strain. By inducing the expression of IHF with Atc in this strain, we ensured the expression of IHF in the absence of Fis.

#### Replication of experiments

Data from high-throughput methods to measure gene expression are intrinsically noisy. The reasons include biological cell-to-cell expression variation ^132^, as well as technical ^136–139^ variation (e.g. pipetting errors, equipment biases, limit of detection thresholds etc.). To account for such variability, we performed sort-seq in three replicates, i.e., we performed three independent sorting procedures for the same overnight culture.

#### Regulation strengths

Due to gene expression and measurement noise, individual TFBS variants in a sort-seq experiment usually appear in more than a single bin, and their read count (frequency) varies among bins^140–143^. Following established practice^70,142,144^, we used a weighted average of these frequencies for each variant to represent the mean expression level driven by the variant. To facilitate interpretation, we converted this expression level into a regulation strength relative to the wild type binding site (regulation strength of one) for that specific TF. Values below one indicate weaker TF binding (higher GFP expression), whereas values above one indicate stronger regulation strength (lower GFP expression).

#### Calculating crosstalk levels

Crosstalk levels were calculated as the geometric mean of the regulatory strengths of both transcription factors for each sequence. Each TF’s regulatory strength was normalized to a maximum value of 1, corresponding to its highest observed regulation, which corresponded to the WT sequence for each cognate TF. Consequently, the crosstalk levels were bounded between 0 and 1.

#### Validating regulation strengths with plate reader measurements

To further validate our regulation strengths thus inferred, we chose 20 DNA binding sites that cover a wide range of measured regulation strengths. We cloned these sequences into the appropriate vector pCAW-Sort-Seq-V2-TF (TF: Fis or IHF) and transformed them into the appropriate mutant strain to ensure that only Fis or IHF are expressed. We picked individual colonies and grew them overnight (16 hours, 37°C, 220 rpm) in liquid LB supplemented with 50 μg/mL of chloramphenicol and anhydrotetracycline. We diluted the cultures to 1:10 (v/v) in cold Dulbecco’s PBS (Sigma-Aldrich #D8537) to a final volume of 1 mL. We transferred 200 μl of the diluted cultures into individual wells in 96-well plates and measured GFP fluorescence (emission: 485nm/excitation: 510nm, bandpass: 20nm, gain: 50) as well as the optical density at 600nm (OD_600_) as an indicator of cell density. We then normalized fluorescence by the measured OD_600_ value to account for differences in cell density among cultures and compared the obtained ratios to the previously inferred regulation strengths for the 20 selected variants. We performed all such measurements in biological and technical triplicates (three colonies per sample and three wells per colony, respectively).

## Data availability

The DNA sequencing data generated in this study have been deposited in the NCBI database (BioProject) under the accession code: PRJNA1162486 [https://www.ncbi.nlm.nih.gov/bioproject/?term=PRJNA1162486]. The flow cytometry and plate reader data generated in this study have been deposited in the Zenodo public repository and are accessible publicly via the following DOI: 10.5281/zenodo.14008581 [https://zenodo.org/records/14008581].

## Code availability

The computer code generated in this study has been deposited in the Zenodo public repository and is publicly accessible via the following DOI: 10.5281/zenodo.14008581 [https://zenodo.org/records/14008581]

## Acknowledgements

We gratefully acknowledge financial support from the Swiss National Science Foundation grant 310030_208174. We also extend our thanks to the UZH University Priority Research Program in Evolutionary Biology and the UZH flow cytometry facility for their technical support. We are especially thankful to Andrei Papkou for engaging in theoretical discussions.

## Author contributions

C.A.W. and A.W. conceived the study and designed experiments. C.A.W. carried out experiments. C.A.W. wrote computer code to carry out bioinformatic work and data analysis. A.W. provided analytical guidance in data analysis. C.A.W. generated figures. L.G. wrote computer code to carry out simulations. C.A.W. and A.W. wrote the paper, which was edited by all authors.

