## Supporting Information for "Entangled adaptive landscapes facilitate the evolution of gene regulation by exaptation"

**Author list:** Cauã Antunes Westmann[1,2], Leander Goldbach[1,2], Andreas Wagner[1,2,3*]

**Affiliations:**
[1] – Department of Evolutionary Biology and Environmental Studies, University of Zurich, Winterthurerstrasse 190, Zurich CH-8057, Switzerland
[2] – Swiss Institute of Bioinformatics, Quartier Sorge-Batiment Genopode, 1015 Lausanne, Switzerland
[3] – The Santa Fe Institute, Santa Fe, NM 87501, USA

### Supporting Information - Index

**Supplementary Methods**

1. **General cloning procedures**
    - **1.1.** *Overnight incubation of cultures in liquid and solid medium*
    - **1.2.** *PCR*
    - **1.3.** *Verifying PCR products through gel electrophoresis*
    - **1.4.** *DNA purification with commercial kits*
    - **1.5.** *Gibson assembly*
    - **1.6.** *Preparation of electro-competent cells*
    - **1.7.** *Electroporation*
2. **The design of vector pCAW-Sort-Seq-V2**
3. **Construction of the plasmid pCAW-Sort-Seq-V2 and its variants**
4. **Library design, synthesis, and cloning**
5. **Analyzing and sorting cells**
6. **DNA extraction and sequencing**
7. **Data analysis**
    - **7.1.** *Filtering and preparing sequencing reads*
    - **7.2.** *Calculating regulation strengths*
    - *7.3. Assessing comparability across TFs and strain backgrounds*
    - **7.4.** *Combining data from triplicates*
    - **7.5.** *Frequency matrix and sequence logo*
    - **7.6.** *Creation of genotype networks and network metrics*
    - *7.7. TFBS analysis between conserved intergenic regions from E. coli and Salmonella*
    - **7.8.** *PWM analysis of genomic sequences*
    - **7.9.** *Clustering of E. coli PWMs*
8. **Simulated adaptive walks**
    - *8.1. General framework for adaptive walks*
    - *8.2. Kimura walks*
    - ***8.3.*** *Population sizes and selection strength*
    - *8.4. Starting genotypes and number of walks*
    - *8.5. Greedy walks*
    - **8.6.** *Adaptive walks when selection favors dual regulation*

**Supplementary tables**

**S1. Primers for engineering the plasmid pCAW-Sort-Seq-V2 and cloning libraries**
**S2. TFBS reference sequences used in this study.**
**S3. Libraries used in this study**
**S4. Table of strains**
**S5. Table of plasmids used in this study**
**S6. Table of sequences for CRP-Fis library regulation strength validation**
**S7. Table of sequences for CRP-IHF library regulation strength validation**
**S8. Table of sequences for Fis-IHF library regulation strength validation**
**S9. CRP-Fis landscape metrics**
**S10. CRP-IHF landscape metrics**
**S11. Fis-IHF landscape metrics**

**Supplementary Figures**

**S1. Construction of pCAW-Sort-Seq-V2**
**S2. Components of plasmid pCAW-Sort-Seq-V2**
**S3. Library design**

**S4. Growth profile for strains used in this study**
**S5. Combinatorial characterization of wild-type TFBSs in all mutant *E. coli* strains of this study**
**S6. Distribution of fluorescence levels for controls and CRP-Fis library under CRP or Fis expression**
**S7. Distribution of fluorescence levels for controls and CRP-IHF library under CRP or IHF expression**
**S8. Distribution of fluorescence levels for controls and Fis-IHF library under Fis or IHF expression**
**S9. Calibration of Atc-dependent mScarlet-I expression**
**S10. Pairwise associations of sequencing read counts for replicates of the CRP-Fis library**
**S11. Pairwise associations of sequencing read counts for replicates of the CRP-IHF library**
**S12. Pairwise associations of sequencing read counts for replicates of the Fis-IHF library**
**S13. Specific plasmid-strain combinations ensure the presence of only one of two TFs during our measurements**
**S14. Effects of TF1 and TF2 regulation strength ratios on TFBS nucleotide composition**
**S15. Most paths to peaks are accessible across landscapes**
**S16. TFBS shifts for global regulators in 50 conserved promoter regions between *Salmonella typhimurium* and *E. coli***
**S17. Rapid exaptation on the CRP-IHF landscape under the SSWM regime**
**S18. Rapid exaptation on the Fis-IHF landscape under the SSWM regime**
**S19. Frequencies of sequences and paths among adaptive walks**
**S20. Little path overlap, moderate sequence overlap, and strong path preferences during adaptive walks towards stronger binders for a specific TF**
**S21. Adaptive walks tend to pass through strong TFBSs more often**
**S22. Drift has negligible effects on adaptive walks**
**S23. Rapid exaptation on the CRP-Fis landscape outside of the SSWM regime**
**S24. Rapid exaptation on the CRP-IHF landscape outside of the SSWM regime**
**S25. Rapid exaptation on the Fis-IHF landscape outside of the SSWM regime**
**S26. PWM scores are associated with measured regulation strengths**
**S27. Clustering of *E. coli* PWMs for exploring potential crosstalk candidates**
**S28. Clustering of PWMs for *E. coli's* global regulators**
**S29. Hierarchical clustering of predicted crosstalk for experimentally validated TFBSs across CRP, Fis, and IHF.**
**S30. Rapid evolution of crosstalk in exaptation landscapes**
**S31. Drift has negligible effects on adaptive walks favoring crosstalk**
**S32. Rapid evolution of crosstalk in exaptation landscapes outside of the SSWM regime**

### Supporting Information

#### Supplementary Methods

##### 1. General procedures

All general procedures have been previously described in ref.[1]. We briefly describe them below for completeness.

*Overnight incubation of cultures in liquid and solid medium*

We cultivated bacteria in liquid LB medium (using either 15 mL or 50 mL Falcon tubes), enriched with chloramphenicol at a concentration of 50 µg/mL. We incubated these cultures for a period of 16 hours at a temperature of 37°C, with a shaking speed of 200rpm and 50 mm orbital motion, using an Infors HT Multitron Incubator Shaker. Similarly, for cultures in solid medium, we grew bacterial colonies on LB-agar plates (using sterile plastic petri dishes of 90mm × 15mm dimensions), also supplemented with chloramphenicol at a concentration of 50 µg/mL, and incubated for the same time and at the same temperature.

*PCR*

Except where specifically mentioned, we carried out the amplification of DNA fragments through PCR, employing Q5® high-fidelity polymerase (NEB #M0491L) to minimize mutation introduction into the amplicons. We followed the protocol recommended by NEB, aiming for a final reaction mixture of 50uL. We conducted each PCR twice, and combined the products from these duplicates after the completion of the reaction. We determined the primer melting temperatures (Tm) using the NEB Tm calculator (accessible at https://tmcalculator.neb.com/#!/main), with primers at a concentration of 500nM.

*Verifying PCR products through gel electrophoresis*

Unless indicated otherwise, we verified successful PCR amplification via gel electrophoresis to ensure the presence of singular-band amplicons and the absence of non-specific bands. This process involved separating PCR products in a 0.8% agarose Tris-EDTA (TAE) gel. We conducted electrophoresis for 45 minutes at 120V, or until the bands had progressed beyond the halfway point of the gel's total length.

*DNA purification with commercial kits*

Once we had confirmed a successful PCR through gel electrophoresis, we purified the PCR products utilizing the Monarch® DNA PCR/Gel Extraction Kit (NEB #T1020L), adhering to the manufacturer's protocol. In scenarios requiring further purification (such as the occurrence of non-specific bands post-PCR), we employed a gel purification step. This involved mixing 10 µL of 6x NEB DNA dye with each 50 µL PCR product, then loading all 60 µL onto a 1% agarose gel. We carried out electrophoresis for 45 minutes at 120V, or until the bands had moved more than halfway through the gel. We then excised the DNA band corresponding to the amplified sequence using a scalpel. For extracting DNA from the gel, we used the Monarch® DNA Gel Extraction Kit (NEB #T1020L).

*Gibson assembly*[2]

We assembled PCR-amplified fragments using the NEBuilder-HiFi® DNA Assembly Master Mix kit (NEB #E2621L). We determined the molarity required for the assembly based on the guidelines outlined by the Barrick Lab (details available at https://barricklab.org/twiki/bin/view/Lab/ProtocolsGibsonCloning). We incubated the assembly mix for one hour at 50°C in a dry bath incubator, followed by cooling on ice for subsequent steps.

*Preparation of electro-competent cells*

We utilized glycerol/mannitol step centrifugation to prepare electro-competent cells [3]. To this end, we first cultured selected *E. coli* strains (**Supplementary Table S4**) in 5 mL SOB medium at 37°C with a shaking speed of 250 rpm overnight. The next day, we transferred 3 mL of these cultures to 300 mL of SOB medium and incubated under the same conditions until the $OD_{600}$ reached a value between 0.4 and 0.6 (measured at an optical path length of 1 cm), which took approximately 2-4 hours. After cooling the culture on ice for 15 minutes, we centrifuged the cells at 4°C and 1,500 g for 15 minutes. We then resuspended the cells in 60 mL of ice-cold distilled $H_2O$ and divided them into three 50 mL tubes. Gradually, we added 10 mL of an ice-cold glycerol/mannitol solution (consisting of 20% glycerol (w/v) and 1.5% mannitol (w/v)) to each tube using a 10 mL pipette. We centrifuged the tubes at 1,500 g and 4°C for 15 minutes in an Eppendorf 5810/5810 R centrifuge with acceleration/deceleration set to zero. After discarding the supernatant, we resuspended cells in 3.0 mL of the same glycerol/mannitol solution. We transferred these suspensions, each now 100 µL in volume, to pre-cooled 1.5 mL tubes, and incubated them in a dry ice-ethanol bath for about 1 minute. Finally, we stored these suspensions at -80°C for future transformation experiments.

*Electroporation*

In all transformation experiments described in this study, we used 100 µL of electrocompetent cells for electroporation, employing 0.2 cm cuvettes (EP202, Cell Projects, UK) and a Micropulser electroporator (Bio-Rad) set to the EC3 setting (15k V/cm). Post-electroporation, we recovered cells in 1 mL of SOC medium, warmed in advance in 15 mL Falcon tubes. This recovery step lasted for 1.5 hours at 37°C with a shaking speed of 220 rpm. Unless specified otherwise, we spread 300 µL of each recovered culture on a LB agar plate that contained 50 µg/mL chloramphenicol. We then incubated this plate overnight for 16 hours at 37 °C. Subsequently, we confirmed the identity of the clones on the plate via Sanger sequencing.

#### 2. The design of vector pCAW-Sort-Seq-V2

The plasmid pCAW-Sort-Seq-V2 is a derivative of the plasmid pCAW-Sort-Seq[1]. Briefly, plasmid pCAW-Sort-Seq harbours a pBBR1 replication origin, which ensures a broad host range and maintains a low copy number—typically between 5 to 10 copies per cell[4] – a chloramphenicol resistance gene, a TetR repression system[5] and a TFBS measuring module, consisting of an interchangeable TFBS between a constitutive promoter and a superfolder GFP (*sfgfp*[6]) reporter gene. In this system, TF-TFBS interactions can decrease GFP production by physically obstructing the bacterial RNA polymerase, a phenomenon known as steric hindrance. The reporter gene *sfgfp* is insulated by a transcriptional insulator named RiboJ, a synthetic ribozyme that removes 5'UTR interferences from variable TFBS sequence in the mRNA by self-cleavage[7,8]. More details and features have been previously described in ref. [1]. Here, we have integrated a bicistronic expression cassette into our plasmid to enable the regulated expression of a TF of choice. This cassette is designed with the gene encoding the focal TF situated upstream of a *mscarlet-I*[9] reporter gene to monitor the expression of this bicistronic operon via fluorescence. Regulation is achieved through the *pLtetO-1*[5] promoter, a synthetic promoter tightly repressed by TetR. Expression is initiated by the addition of anhydrotetracycline (Cayman Chemicals, catalog #10009542), which relieves TetR-mediated repression, activating the expression of the entire cassette.

#### 3. Construction of the pCAW-Sort-Seq-V2 plasmid and its variants

Following the assembly of plasmid pCAW-Sort-Seq-V2 and its variants (pCAW-Sort-Seq-V2-CRP, pCAW-Sort-Seq-V2-Fis, and pCAW-Sort-Seq-V2-IHF), we cloned selected wild-type

TFBSs into these plasmids (**Supplementary Table S5**). Initially, we designed these TFBSs using the Snapgene® software (available at snapgene.com), and had them synthesized by IDT (Coralville, USA) as 140bp single-stranded DNA Ultramer® oligonucleotides (4nmol). We resuspended each TFBS in nuclease-free water and diluted it to a concentration of 50ng/uL. The Ultramers® served as templates in PCR for generating double-stranded DNA fragments and amplifying the TFBS sequences.

The PCR conditions were 98°C for 30 seconds; 25 cycles at 98°C for 10 seconds, 60°C for 15 seconds, and 72°C for 80 seconds; followed by a final elongation at 72°C for 5 minutes. Post-PCR, we used gel electrophoresis to verify the presence of a single 140bp band for each TFBS. We then purified these bands using the Monarch® DNA gel extraction kit (NEB #T1020L).

Subsequently, we digested both the appropriate pCAW-Sort-Seq-V2-TF plasmid and the amplified Ultramers® using HindIII-HF (NEB #R3104) and BamHI (NEB #R3136) enzymes. We performed ligation with a 10:1 insert-to-vector ratio, using 100ng of the vector backbone, 10 units of T4 DNA ligase (NEB #M0202L), and 2 µL of 10X ligation buffer in a 20 µL reaction. We purified the ligation product using the Monarch® DNA PCR/Gel Extraction Kit (NEB #T1020L) and eluted in 15 µL of dH2O. We then introduced this product into *E. coli* DH5α cells via electroporation. We verified correct cloning with Sanger sequencing of the TFBS (NightSeq® service, Microsynth, Switzerland), followed by full sequencing of the plasmid (Full PlasmidSeq® service, Microsynth, Switzerland). We cultured positive clones overnight in liquid media, stored them in 20% glycerol at -80°C, and extracted their plasmids using the QIAprep spin miniprep kit (Qiagen, Germany) for further cloning steps.

We used the steps just described to successfully construct all plasmids listed in **Supplementary Table S5**. We then extracted these plasmids from the *E. coli* DH5α cloning strain and transformed into the various mutant strains (Δ*crp*, Δ*fis*, Δ*ihf*, as detailed in **Supplementary Table S4**) for further experimentation.

#### 4. Library design, synthesis and cloning

For each TF-pair we studied, we aligned one strongly binding TFBS (**Supplementary Table S2** and **Supplementary Figure S3**) for each member of the pair in a pairwise manner using a local alignment method[10] with a high gap penalty to avoid indels. For each TFBS we performed this alignment both with the TFBS and its reverse complement to identify the optimal alignment, i.e., the alignment characterized by the fewest mismatches. For each pairwise comparison, we thus established a consensus sequence and denoted allowed nucleotide variation in this consensus using the IUPAC nomenclature for ambiguous nucleotides (**Supplementary Table S3** and **Supplementary Figure S3**). This approach allows for no more than two variable nucleotides per position. Thus, the theoretical library size for this method is $2^n$, where $n$ is the number of variable positions in each consensus sequence. In this way, we calculated the library sizes for the TF pairs CRP-Fis, CRP-IHF, and Fis-IHF as $2^8 = 256$, $2^7 = 128$, and $2^7 = 128$ sequences, respectively.

We designed each of the three resulting libraries with the Snapgene® software (snapgene.com), and had it synthesized as 140bp single-stranded DNA Ultramer® oligonucleotides (4nmol) by IDT (Coralville, USA). We resuspended the library in nuclease-free water and diluted it to 50ng/uL. The Ultramers® served as templates in a PCR for double-stranded DNA fragment formation and library amplification. We used the following PCR program: 98°C for 30 seconds; 25 cycles of 98°C for 10 seconds, 60°C for 15 seconds, and 72°C for 80 seconds; followed by a final extension at 72°C for 5 minutes. To minimize amplification bias, we used a maximum of 25 PCR cycles. Post-amplification, we confirmed the presence of single 140bp bands for each library via gel electrophoresis, and purified these bands using the Monarch® DNA gel extraction kit (NEB #T1020L).

For the construction of each plasmid-based library, as depicted in **Figure 4.2a**, we digested 1μg of the purified library with HindIII-HF (NEB #R3104) and BamHI (NEB #R3136) enzymes in a 100μL reaction, followed by overnight incubation at 37°C. We isolated the appropriate cloning plasmid using the QIAprep spin miniprep kit (Qiagen, Germany), and digested it with the same enzymes. Post-digestion, we added 3μL of the Quick CIP phosphatase to prevent self-ligation by dephosphorylating DNA ends. We purified samples using the Monarch® DNA gel extraction kit (NEB #T1020L).

We performed the ligation with a 10:1 molar ratio of insert-to-vector, using 100ng of vector, 10 units of T4 DNA ligase (NEB #M0202L), and 2 μL of 10X ligation buffer in a 20 μL reaction. We incubated the mixture at 20-22°C for approximately 16 hours, followed by 10-minute inactivation of the ligase at 65°C. We purified the ligation product, resulting in 10 μL of resuspended DNA in distilled $H_2O$. We transformed *E. coli* SIG10-MAX® cells by electroporation with this purified product.

Post-transformation, we plated 50 μL of each recovered culture on LB agar for colony forming unit counting (cfu) and transformation efficiency estimation. We diluted the remaining 950 μL in 9 mL of LB medium with chloramphenicol and cultured it overnight. We aliquoted the culture into 1 mL cryotubes with 20% glycerol and stored at -80°C. From the agar plates, we selected 30 colonies for colony PCR and Sanger sequencing to assess library diversity

(NightSeq® service, Microsynth, Switzerland). Our mean transformation efficiency was $10^6$ cells per transformation for the SIG10-MAX® strain.

After transforming plasmid libraries into the SIG10-MAX® strains for reasons of transformation efficiency and plasmid maintenance, we extracted the plasmids using a QIAprep spin miniprep kit (Qiagen, Germany), and transformed them into the appropriate host strain (see the strain combinations of **Figure 4.2a**). We cultured the transformed strains, each harboring a library, overnight, aliquoted them in 1 mL cryotubes with 20% glycerol, and stored them at -80°C for further experimentation.

#### 5. Analysing and sorting cells

In preparation for cell sorting, we cultivated cells harboring each library in liquid LB medium enriched with chloramphenicol. Specifically, we cultured 1 mL of transformed cell aliquots and a streak of cells with a control plasmid (a promoterless pCAW-Sort-Seq-V2 plasmid lacking sfGFP expression) in 50 ml Falcon tubes with 9 mL LB medium (containing 50 µg/mL chloramphenicol) overnight. Subsequently, we diluted these overnight cultures at a 1:100 ratio (v/v) in LB medium with chloramphenicol and divided them into two aliquots. We supplemented one aliquot with the inducer anhydrotetracycline (Atc) to express the plasmid-encoded TF. The other aliquot remained unchanged. To reduce cell-to-cell noise, we normalized GFP fluorescence from flow cytometry by mScarlet-I fluorescence, which is expressed from the same bicistronic operon as the TF and thus serves as a proxy for TF expression level. For plate reader-based validation assays (**Figure 2d-f**), we normalized GFP fluorescence by OD600 to account for differences in cell density, as described in the Methods. We incubated both cultures for 5 hours until late-exponential/early-stationary phase (200RPM,

37°C). Then we diluted 20 µL of the cultures in 1 mL of cold filtered Dulbecco's PBS (Sigma-Aldrich #D8537) in 15 mL FACS tubes.

We performed FACS-sorting on a FACS Aria III flow cytometer (BD Biosciences, San Jose, CA) using a 70 µm nozzle. We utilized a 488 nm laser for detecting forward scatter (FSC) and side scatter (SSC) with a 488nm/10nm band-pass filter. We set the flow rate to 1.0, adjusting sample dilution as necessary to achieve no more than ≈10,000 events/second. Considering the small size of bacterial cells, we reduced the particle detection threshold to the lowest feasible setting (200 arbitrary units on FSC and SSC channels), increasing it to a maximum of 500 units if background noise was excessive. We then adjusted FSC-H and SSC-H for cells with the negative control plasmid to center the bacterial population in the cytometer's visualization panel. We based the sorting and binning of cells on both mScarlet-I (PE-Texas-Red channel, excitation laser: YellowGreen 561nm, LP filter: 600 nm, BP filter: 610/20nm) and sfGFP fluorescence (FITC channel, excitation laser: 488nm , LP filter: 502nm, BP filter: 530nm/30nm), setting both the PE-Texas-Red and FITC channels voltages so that the median fluorescence of the negative control was between 0 and 100 (arbitrary units) on the FITC-H and PE-Texas-Red-H axes.

Initially, the sorting process involved establishing a gate for identifying cells that are red-fluorescence-positive , i.e., reporter and thus TF-expressing. To establish this gate, we began by measuring the autofluorescence of our negative control culture on the PE-Texas-Red-H axis. Thereafter, we examined the fluorescence of a positive control that had been induced with Atc and was expressing the mScarlet-I protein. We then configured a gate around the positive mScarlet-I -expressing population, ensuring that all cells sorted through this gate exhibited a comparable level of reporter expression to the positive control.

Next, we established the green-fluorescent sorting gates. For setting sorting gates on the FITC-H axis, we first recorded the autofluorescence of the negative control culture. This median autofluorescence defined the upper boundary of the lowest bin (B1) for the experimental population. To maintain statistical robustness and minimize sampling error in our downstream analyses, we wanted to ensure that each unique sequence in our library was represented by at least 100 cells after sorting. In addition, it was crucial to account for an anticipated diversity loss of up to 70% of the cell population during sorting, due to factors such as cell death and the dilution of low-frequency and lower-fitness genotypes during the post-sorting recovery phase [11]. To establish the number of cells required to be sorted initially, we thus determined a multiplicative factor based on the minimally needed number of 100 cells per sequence and the expected cell retention rate post-sorting of 30%, i.e., cells/sequence. We thus multiplied our initial library size by 330 for sorting purposes, i.e., we set the total of 84,480 cells for the CRP-Fis library and 42,240 cells for the Fis-IHF library for the sorting procedure. This calculation aims to ensure that despite a substantial reduction in cell numbers post-sorting, each unique sequence remains adequately represented in the surviving cell population.

Before sorting, to set our sorting gates, we assessed the fluorescence of the aforementioned number of cells expressing sfGFP for each library. We also analyzed Atc-induced and uninduced control samples that contained the wild-type sequences for each plasmid, as shown in **Supplementary Figures S6-S8**. This analysis allowed us to define two distinct thresholds based on the geometric mean of the fluorescence levels of these controls. We positioned the threshold set by the induced control above the autofluorescence level, delineating a lower expression limit that corresponds to the fluorescence distribution in cells where the focal TF is strongly binding to promoter variants. Conversely, the threshold established by the uninduced control marks a higher expression limit, indicative of the fluorescence distribution in cells where the focal TF is not binding to promoter variants (refer to **Supplementary Figures S6-S8** for details). We placed our four fluorescence bins between these thresholds, equidistantly on a binary logarithmic ($log_2$) scale.

We then determined the fraction of the recorded cells within each gate as the number of cells to be sorted into each bin. We sorted cells into 1.5 mL Eppendorf tubes, each containing 500 µL of LB medium, and kept at 4 °C to prevent growth during sorting and sample processing. We replicated the entire sorting procedure three times, each based on independent library transformations.

We added 1mL of LB without antibiotics to each tube, and used 20 µL for serial dilutions ($10^{-4}$ and $10^{-6}$), plated on LB-Cm agar plates to estimate post-sorting viability via colony forming unit (cfu) counting. We transferred the rest of the culture (980 µL) to 50 mL Falcon tubes for a 2-hour recovery period (37°C, 220 rpm), followed by overnight growth with added chloramphenicol for glycerol stock preparation, binning reassessment, and plasmid DNA extraction for PCR and sequencing. The cfu counts from the diluted samples allowed us to estimate that across bins 85% of cells remained viable (standard deviation: 17%). We assessed library genetic diversity through Sanger sequencing of colonies (NightSeq® service, Microsynth, Switzerland).

To reassess our binning procedure, we regrew the sorted cultures and analyzed their expression distributions via flow cytometry, replicating the original expression measurements. This reassessment aimed to verify if the post-sorting fluorescence distribution matched the pre-sorting distribution for each bin. During this reassessment, we compared the distribution of GFP fluorescence for each bin and their respective geometric means from the day of the sorting procedure to those observed in our samples post-sorting. We preferred the geometric mean over the arithmetic mean for this analysis because it is less influenced by outliers and better represents the central tendency of data that is log-normally distributed, which is typical for fluorescence measured during flow cytometry [12,13]. If the distributions and geometric mean for a bin did not match between the sorting day analysis and the post-sorting one, we re-sorted cells for that specific bin.

#### 6. DNA extraction and sequencing

We diluted 500 µL of individual glycerol stocks of each replicate subpopulation of sorted cells (i.e., cells from each "bin" of fluorescence intensity) in 5mL of LB supplemented with chloramphenicol in 15mL Falcon tubes, and grew the resulting cell culture overnight (16 hours, 37°C, 220 rpm). On the next day, we isolated plasmids from each culture using a QIAprep® spin miniprep kit (Qiagen, Germany). In order to allow the sequencing of multiple pooled samples (multiplexing), we barcoded our regulatory region through PCR with specific HPLC-purified primers (**Supplementary Table S6**) provided by Eurofins (Konstanz, Germany). We added barcodes to the 5'region of the amplicon through a PCR. We performed this PCR with the Q5 high-fidelity polymerase in triplicate for each sample. To calculate primer melting temperatures (Tm), we used the NEB Tm calculator (https://tmcalculator.neb.com/#!/main) for a primer concentration of 500nM. We performed the PCR with the following program: 98°C/30 s; 25 cycles of 98°C/10 s, 64°C/30 s and 72°C/30 s; and 1 cycle of 72°C/2 min.

After PCR amplification, we digested the reaction products with the restriction enzymes DpnI (NEB #R0176L) and Exonuclease I (NEB #M0293L) in order to remove traces of genomic DNA, plasmids, and single-stranded DNA that could interfere with sequencing. The Master Mix we used for a single digestion harbored 1 µL of 10x CutSmart® Buffer (NEB #B6004S),

1 µL of Exonuclease I (NEB #M0293L), 1 µL of DpnI restriction enzyme (NEB #R0176L), and 7 µL of distilled nuclease-free water. For each PCR product, we added 10 µL of the Master Mix. We incubated the reaction for 1 hour at 37°C, following 15 minutes at 80°C for deactivation of the enzymes.

After digestion, we purified samples using the Monarch® DNA PCR/Gel Extraction Kit (NEB #T1020L). We analyzed samples through gel electrophoresis to confirm that only a single band with a size ≈150bp was present for each of them. After confirmation, we pooled the samples from the different bins of each replicate equimolarly to a total mass of 2,600 ng and a volume of 100 µL (26ng/µL of DNA) in 1.5 mL Eppendorf tubes. We then sent the pooled samples for adapter ligation and sequencing at Eurofins (NGSelect Amplicons® on Illumina HiSeq), obtaining 15 million paired-end reads (2 × 150 bp) for all samples.

We quantified the total number of reads ($t$) required for sequencing based on several factors, including the number of bins ($b$), biological replicates ($r$), strains ($s$), genotypes ($g$), and the reads per genotype ($p$). Using these variables, we first calculated the total number of needed barcodes as:

$$n_{bc} = b \times r \times s$$

For any one of our TFs, the values of these parameters are $b = 4$ bins, $r = 3$ biological replicates, and $s = 2$ strains, yielding $n_{bc} = 24$. Subsequently, the total number of required reads can be determined through the equation:

$$t = n_{bc} \times g \times p$$

where we aimed for $p = 50$ reads per genotype. The number g of genotypes per library differs for each TF combination. Specifically, the CRP-Fis library harbors $g = 256$ genotypes, while the CRP-IHF and Fis-IHF libraries harbor $g = 128$ genotypes each. These quantities lead to a total of $t=n_{bc}\ \ g\ \ p$ = 24 x 50 x 256 = 307,200 required reads for the CRP-Fis library and, analogously, t = 153,600 reads each for the CRP-IHF and Fis-IHF libraries.

#### 7. Data analysis

##### 7.1. Filtering and preparing sequencing reads

We processed the sequencing data with a blend of custom Python and awk scripts, complemented by established bioinformatics utilities. Firstly, we trimmed sequences by computationally excising Illumina adapters, followed by merging paired-end reads. We then organized these paired reads into distinct files, each tagged with a sequence barcode identifying the sequence bin from which its reads originated.

For the removal of Illumina adapter sequences from the paired-end reads, we employed Cutadapt [14] with the following parameters:

```
cutadapt -j 8 -e 0.1 --no-indels --overlap=8 --discard-untrimmed \
  -a "^\$FWD...\$REV_RC;max_error_rate=0.2;min_overlap=6" \
  -A "^\$REV...\$FWD_RC;max_error_rate=0.2;min_overlap=6" --pair-filter=any \
  -o 'Sample_${sample}_trimmed_1.fastq.gz' \
  -p 'Sample_${sample}_trimmed_2.fastq.gz' \
  --max-ee = 2 -l = 114 \
  $reads
```

Here, “ADAPTER_FWD” and “ADAPTER_REV_RC” are placeholders for the actual adapter sequences used, which are 5’-AGATCGGAAGAGCACACGTCTGAACTCCAGTCA-3’ for read 1 of the pair, and 5’-AGATCGGAAGAGCGTCGTGTAGGGAAAGAGTGT-3’ for read 2 of the pair.

Given that our amplicons were brief (153 bp) and the paired-end reads fully overlapped, merging of the paired-end reads was necessary, for which we used the FLASH software [15]. After the refinement and merging of reads, we demultiplexed the merged reads into their respective bins and replicates with the following FLASH parameters:

```
flash  -t $ncores $reads -O -m 60 -M 140 -z -o 'Sample_${sample}_merged'
```

Next, we utilized the FastX toolkit (http://hannonlab.cshl.edu/fastx_toolkit/)[16] to discern and preserve only those sequences that exceeded a high quality threshold of Q = 33. Subsequently, we filtered our data to exclude sequences with mutations or indels outside the variable TFBS library region within the RiboJ sequence to avoid the unwanted effect of 5'UTR mutations that are known to change gene expression levels[17]. We performed this filtering step using a custom Python script. Thereafter, we used custom *awk* and *R* [18] scripts to convert the data into a table. Each row of this table contains data from one TFBS in the library, and each column contains the number of reads for this TFBS from one of the four bins into which we had sorted cells. We estimated the average regulation strength of each TFBS from this data.

#### 7.2. Calculating regulation strength levels

We calculated regulation strengths as previously described in ref. [1]. Briefly, in sort-seq experiments, sequences often appear in multiple fluorescence bins due to random mis-sorting[19,20] and the stochastic nature of gene expression[21]. Following methods established in previous research [22–24], we determined the reporter expression level driven from each TFBS variant by calculating a weighted average of bins in which the variant occurred. This involved multiplying the frequency of each sequence ($x_i$) in a given bin $i$ by a numerical value representing that bin ($w_i$ = 1,2,3,4), and then averaging these products over the total sequence count. Mathematically, this weighted average calculates as

$$e = \sum_{i=1}^{n} \frac{(x_i \times w_i)}{\sum_{i=1}^{n} x_i},$$

where $e$ is the expression level driven by the sequence.

This approach yields a continuous spectrum of expression levels $e$ in the range of 1 to 4. High expression values indicate robust GFP expression and thus weak binding of a TF to a TFBS variant. To facilitate interpretation, we inverted this scale so that higher values indicate stronger repression and lower GFP expression, i.e., we define the regulation strength $b$ as

$$b = e_{max} + 1 - e,$$

where $e_{max} = 4$ is the maximal expression level. We then normalized $b$ by the regulation strength of the corresponding wild-type TFBS, which in all cases was the maximal observed regulation strength for that TF. The resulting normalized regulation strength scores $b_{norm}$ vary from 0 to 1, where low scores denote TFBS variants with weak TF binding (weak reporter repression), and high scores indicate variants with strong TF binding (strong reporter repression). A score of $b_{norm} = 1$ signifies the regulation strength (repression) of the wild-type TFBS.

#### 7.3. Assessing comparability across TFs and strain backgrounds

To assess whether WT-normalized regulation strengths can be compared across TFs and strain backgrounds, we quantified both baseline GFP expression and repression by the cognate wild-type TFBSs (Supplementary Figure S5). Baseline GFP expression in the absence of induction was similar across TFBS variants and strain backgrounds, with a coefficient of variation of

2.9% across all 12 experimental conditions. Although a two-way ANOVA detected a small effect of strain background on baseline expression ($F(2,24) = 8.15$, $P = 0.002$), we observed no significant interaction between TFBS sequence and strain background ($F(6,24) = 1.78$, $P = 0.15$), indicating that differences among TFBS variants are consistent across strains. In addition, the cognate wild-type TFBSs ($WT_{CRP}$, $WT_{Fis}$, and $WT_{IHF}$) produced similar absolute repression levels in the three strain backgrounds, indicating comparable dynamic ranges for all three TFs in our assay. Together, these observations support comparison of WT-normalized regulation strengths among TFs.

In all experiments, we used a fixed Atc concentration of 100 ng/mL, corresponding to approximately 98% of maximal induction in the Atc-dependent mScarlet-I calibration curve (Supplementary Figure S9). We therefore measured regulatory output at a near-maximal and approximately constant TF concentration. This design minimizes variation in TF abundance across conditions and allows differences in reporter output to be interpreted primarily as consequences of cis-regulatory sequence variation.

##### 7.4. Combining data from triplicates

As a quality-filtering step, we eliminated TFBS variants that did not appear in all three replicates or that were represented by less than 100 reads in total (summing their read counts over the four bins). Given that we have 4 bins, we wanted a minimum of 25 paired-end reads per bin to prevent misinterpretation of regulation strength levels and also guide our threshold's choice. All sequences from our libraries passed this first threshold.

Subsequently, we calculated the regulation strengths $b$ of the remaining TFBS variants (**Supplementary Methods 7.2 and Methods**). We then determined the coefficient of variation of regulation strengths across replicates for each TFBS variant, which estimates the consistency of measured regulation strengths among replicates. Notably, most TFBS variants exhibited a coefficient of variation (CV) below 0.5, which was the threshold we set for filtering sequences. We excluded sequences with a CV above 0.5 from the analysis. This threshold is commonly used in transcriptional studies employing fluorescent reporters to account for transcriptional noise and measurement variability [21]. Finally, we averaged regulation strengths for each TFBS variant across all replicates and normalized the resulting averages by the maximal observed regulation strength among all TFBSs of a given TF.

#### 7.5. Frequency matrix and sequence logo

We generated frequency matrices of nucleotides for TFBSs from each fluorescence bin by counting the frequency of each nucleotide at each variable position of the TFBS library. From this data we generated heatmaps and DNA sequence logos representing the frequency matrices graphically. A sequence logo consists of a stack of the letters A, C, G, and T, at each position of a DNA sequence, where the relative size of each letter indicates its frequency in the sequence. The total height of the stack corresponds to the information content of that position, in bits [25,26]. A sequence logo is a graphical representation of informational properties of a TFBS motif. When mutated, nucleotides with high information content are more likely to lead to a loss of binding (repression) than nucleotides with low information content[27,28].

#### 7.6. Creation of genotype networks and network metrics

We used in-house Python scripts and the Python package *igraph*[29] to generate directed genotype networks. These are graphs in which TFBS variants are nodes (vertices, genotypes), and variants that differ in a single nucleotide are connected by an edge. Each edge is directed, i.e., it corresponds to a binding-score-increasing mutation, and points from the variant with lower regulation strength to the neighbor with higher regulation strength. Each vertex of this network is associated with the corresponding DNA sequence and the associated regulation strength. We used in-house python and R scripts for all network analyses described below.

*Epistasis:* Epistasis, non-additive interactions between mutations, can impose severe constraints on molecular evolution, because the mutations that are beneficial in one genetic background may be deleterious in another[30]. Epistasis can be classified as magnitude, simple sign, or reciprocal sign epistasis, depending on the sign (i.e., positive or negative) of the fitness effect of individual mutations and their combinations[30]. In magnitude epistasis, the effect of a mutation on regulation strength varies depending on the genetic background but the sign of this effect (increasing or decreasing regulation strength) does not. Simple sign epistasis occurs if one single mutant has a lower regulation strength than both the wild type and the double mutant, while the other single mutant has a regulation strength that is intermediate to the wild type and double mutant. Reciprocal sign epistasis occurs when both mutations independently decrease regulation strength, but their combination increases regulation strength. The presence of reciprocal sign epistasis is a necessary condition for the existence of multiple peaks in an adaptive landscape [31,32].

To determine the incidence of epistasis in our landscapes, we employed a method that involves identifying all "squares" in a genotype network with the *motifs* function from the *igraph* library in R. Each square consists of a "wild-type" sequence, a double-nucleotide mutant, and the corresponding two single mutants. We assessed epistasis for each square along a single axis by selecting the highest-regulation strength sequence as the double mutant. Our analysis placed each square into one of three categories: no sign epistasis, simple sign epistasis, and reciprocal sign epistasis. The no sign epistasis category included both magnitude epistasis and additivity (no epistasis) without differentiating between them, because neither affects peak accessibility[33]. We determined the proportion of all squares that fell into each category (see **Supplementary Tables S9-S11**).

*Peaks:* A peak is a genotype (TF binding site variant) whose neighbors all convey lower regulation strength than itself. Two peaks are connected if they are neighbors and convey the same regulation strength. We refer to the genotype with the highest regulation strength as the summit or global peak[33,34].

*Accessible Paths:* A mutational path through the genotype network is accessible if and only if the regulation strength increases for each mutational step along the path[33,34]. We enumerated accessible paths of all lengths (mutational steps) exhaustively.

*Sequence and path overlap between different starting points.* We obtained sets of unique visited sequences across all adaptive walks starting from each of our starting genotypes. The sets of visited sequences may overlap. To determine the overlap between two sets (from two different starting sequences) $B_1$ and $B_2$, we used the Jaccard index $J$[35,36], which is equal to the size of the intersection between two sets of variants divided by the size of their union

$$J = \frac{B1 \cap B2}{B1 \cup B2}$$

We also obtained the sets of unique traversed paths (meaning vectors containing genotypes in a specific order) across all adaptive walks starting from each of our starting genotypes and performed the same aforementioned analysis.

#### 7.7. TFBS analysis between conserved intergenic regions from *E. coli* and *Salmonella*

To compare changes in the promoter regions of *E. coli K-12 MG1655* and *Salmonella typhimurium LT2*, we performed the following bioinformatic analysis. First, we obtained annotated whole genome sequences for these species from NCBI (GenBank reference codes: *E. coli K-12 MG1655*: GCA_000005845.2; *Salmonella typhimurium LT2:* GCA_000006945.2). We then identified orthologous proteins in these genomes using the OrthoVenn3 tool[37]. Subsequently, we identified *E. coli* genes that are operon “heads”, i.e., they are the first genes of an operon, using experimental data from RegulonDB[38]. We only analyzed pairs of orthologs further whose *E. coli* member fulfils this criterion. In addition, we only considered orthologs in which the 150 base pairs upstream of each orthologous gene are intergenic, i.e., they do not overlap with any coding region on either strand. We then aligned these intergenic regions globally for each pair of orthologs with the Needleman-Wunsch algorithm[39]. After that, we identified putative TFBSs in each pair of intergenic regions and compared them. To this end, we used PWMs from RegulonDB [40], and a custom script that scans each strand of an intergenic region and calculates PWM scores of putative TFBSs in it. PWM thresholds for this analysis are based on previously established values in the RegulonDB database[40]. We considered high-scoring putative TFBSs as evidence for exaptive evolution when a TFBS for one TF within an intergenic region in one species overlapped with a TFBS for a different TF in the other species. We represented the resulting data in the form of a matrix, in which the orthologous intergenic regions correspond to rows, and the number of binding sites found for each transcription factor correspond to columns. For each of the two organisms and all intergenic regions we studied, this matrix includes the number of binding sites for the TFs CRP, Fis, and IHF, as well as for several other TFs. We then subtracted the matrix of *Salmonella* from that of *E. coli* (and vice versa) to identify the number of gained (positive entries) and lost (negative entries) binding sites in each organism. This subtraction resulted in another matrix, whose entries represent changes in binding site numbers for various intergenic regions and transcription factors, including CRP, Fis, and IHF. The results of this procedure for 50 promoters sampled from the 991 orthologous intergenic regions, are shown in the clustered heatmap of **Supplementary Figure S16**.

We classified a pair of aligned intergenic regions as having experienced a putative exaptation event when: (i) a TFBS for TF1 exists in the *E. coli* intergenic region but not in the aligned *Salmonella* region, (ii) a TFBS for a different TF2 exists in the same *Salmonella* intergenic region but not in the *E. coli* region, and (iii) the two TFBSs (TF1 in *E. coli*, TF2 in *Salmonella*) overlapped by at least one base pair in the global alignment. This criterion identifies cases where one TFBS appears to have replaced another through mutation, consistent with an exaptive event. This overlap requirement is grounded in the theoretical prediction that exapted sites retain greater sequence similarity to the ancestral binding motif than sites that arose de novo, because exaptation proceeds from a pre-existing functional sequence[41]. By requiring that a TFBS for TF1 in one species overlaps positionally with a TFBS for TF2 in the other, we select for cases where the underlying sequence is compatible enough with both motifs to score above threshold for one TF in one lineage and for a different TF in the other — precisely the enhanced cross-motif similarity expected under exaptation. A site arising entirely de novo from non-functional sequence would be unlikely to occupy the same aligned position as a TFBS for a different TF in the sister species.

We excluded cases where TFBSs for both TF1 and TF2 are present in both organisms, because these likely reflect ancestral crosstalk rather than exaptation. As a worked example, consider the 150 bp intergenic region upstream of the acs gene (acetyl-CoA synthetase), which is conserved between E. coli K-12 and *Salmonella typhimurium* LT2. Here we detected a CRP TFBS at positions 42–63 (on the forward strand) in the *E. coli* sequence, with a PWM score above threshold. In the corresponding (aligned) *Salmonella* sequence, this site scored below the CRP threshold, but a Fis TFBS exists that overlaps with positions 48–61 — a region that differs by three nucleotide substitutions between the two species. This pattern (CRP site in *E. coli*, overlapping Fis site in *Salmonella*, with no corresponding Fis or CRP site, respectively, in the other organism) is consistent with an exaptive transition from CRP regulation in *E. coli* to Fis regulation in *Salmonella*, or vice versa, depending on the direction of evolutionary change. We counted such cases as candidate exaptation events. Overall, 34% of TFBS differences between orthologous intergenic regions were consistent with this criterion for exaptation. We emphasize that these are putative events: definitively establishing evolutionary direction would require phylogenetic reconstruction and outgroup comparisons that are beyond the scope of the present study.

#### 7.8. PWM analysis of genomic sequences

In this section, we describe how we generated heatmaps representing TFBS sequences in the *E. coli* K-12 genome, based on data curated from the RegulonDB[38] database. Our RegulonDB dataset encompassed 756 experimentally validated TFBSs, 370 for CRP, 267 for Fis, and 119 for IHF. We downloaded PWMs for each of our three TFs CRP, Fis, and IHF from RegulonDB[38]. Based on the PWM of each TF, we calculated one PWM score for each TFBS, which led to three PWM scores per sequence, one for each TF.

We calculated PWM scores using the *matchPWM* function from the R package *Biostrings*[42], applying a minimum score threshold of 30%. The score of a TFBS is a quantitative prediction of its binding strength to its cognate TF [26,43] (**Supplementary Figure S26**). The matchPWM function returns log-odds scores representing the sum of position-specific log-odds weights for each sequence relative to a background nucleotide frequency. To facilitate comparison across TFs with PWMs of different lengths and information content, we normalized each score by dividing it by the maximum achievable score for that PWM, defined as the sum of the highest log-odds weight at each position. This yields score values in the range [0, 1], where 1 corresponds to the theoretically optimal binding sequence for that TF.

We scanned each TFBS sequence on both strands, and included flanking bases in the scan. For sequences longer than the PWM, the score reported is the maximal log-odds score among all positions of a window that we slid across the sequence in 1 bp increments. This ensures that the highest-affinity sub-sequence is captured regardless of its position within the annotated TFBS.

Because the PWMs and the scanned TFBSs were both derived from RegulonDB[38], we asked whether this shared source could artificially inflate evidence for crosstalk. To assess this possibility, we examined each RegulonDB annotated TFBS class separately and asked how often sites annotated for one TF also received high scores for another TF. This pattern was not restricted to a single TF pair. For example, high cross-TF scores (> 0.5) were common in both CRP-annotated and Fis-annotated sites (45% and 61%, respectively), and analogous analyses for the other TF pairs likewise supported substantial overlap in predicted recognition. In addition, hierarchical clustering of the full PWM-score matrix (**Supplementary Figure S29**) did not separate sequences into distinct groups based on their original TF annotation. Together, these observations argue against the idea that the observed cross-TF signal is simply an artefact of using PWMs and TFBSs derived from the same database.

We visualized the data using a heatmap, in which each row represents one of the 756 TFBS sequences. We used the R package *Pheatmap*[44] to create this heatmap. Each column of the heatmap shows the normalized PWM scores for the TFBSs of one of the three TFs. To reveal patterns of PWM score similarity among TFBSs, we applied hierarchical clustering with complete linkage[44] to the Euclidian distance matrix of the PWM scores. This method groups sequences based on their similarities in PWM scores (**Figure 5a, Supplementary Figure S30**).

##### 7.9. Clustering of *E. coli* PWMs

We clustered Position Weight Matrices (PWMs) of TFBSs from the RegulonDB database [38], which is a comprehensive database cataloging regulatory elements in *E. coli*. The primary outcome of this analysis is a circular dendrogram representing the clustering of PWMs (**Supplementary Figure S27**). For this analysis, we sourced a total of 109 PWMs from RegulonDB[38]. Additionally, we identified and annotated 33 different TF families, represented as colored or numbered circles on the dendrogram. (**Supplementary Figure S27**). For this clustering procedure, we first aligned PWMs, using a combination of local alignment and position weighting through the *DNAmotifAlignment* function from the R package *motifStack*[45]. To align motifs, *DNAmotifAlignment* implements an Ungapped Smith–Waterman (local)[10] alignment algorithm based on column comparison scores calculated by Kullback-Leibler distance[46]. *DNAmotifAlignment* requires that its input be sorted by the distance of motifs. To obtain the sorted motifs for alignment, we calculated these distances using *motifDistance* in *motIV*[45]. We plotted the alignment with the function *plotMotifLogoStack* from the *motifStack* package as a dendrogram that visually represents how distant pairs of PWMs are with respect to the DNA sequences bound by a given TF. (**Supplementary Figures S27-S28**)

#### 8. Simulated adaptive walks

To model evolutionary dynamics on our landscapes, we simulated adaptive walks under the strong-selection weak-mutation (SSWM) regime[23,47–49]. This regime is appropriate for *Escherichia coli*, which has a low mutation rate[50] ($\sim 2 \times 10^{-10}$ per base pair per generation), a large effective population size N[50] ($N \approx 1.8 \times 10^{8}$ individuals) and a short mutational target of only 7–8 variable positions within TFBS libraries. Under these conditions, $N\mu \ll 1$, meaning that populations are monomorphic most of the time: a new mutation arises only after the previous mutation has either become fixed or been lost. In this regime adaptive evolution can be modeled as a Markov chain whose states are genotypes. The transition probabilities of this Markov chain are determined by mutation and fixation probabilities. We describe these transition probabilities in more detail below.

#### 8.1. Fitness definition and conversion from expression measurements

We defined fitness directly as the normalized regulation strength, $b_{\mathrm{norm}}$, which ranges continuously from 0 (no measurable regulation) to 1 (regulation strength equal to that of the cognate wild-type TFBS). We had obtained the regulation strength values from sort-seq fluorescence measurements as a weighted average of bin frequencies, inverted, and normalized to the wild-type regulation strength for each TF (**Supplementary Methods 7.2**). We considered two adaptive scenarios. In the first, selection favors regulation by a single transcription factor, such that we defined fitness as the normalized regulation strength for that TF alone. In the second, selection favors dual regulation, such that we defined fitness as the sum of normalized regulation strengths for both TFs in a pair. In both cases, we calculated the selection coefficient $s$ of a mutation directly from the empirical landscape as the fitness difference between mutant and resident genotype.

#### 8.2. Kimura walks

Our main simulations used fixation probabilities derived by Kimura[51] for a Wright–Fisher population subject to selection and drift. For a mutation from genotype $i$ to genotype $j$, the fixation probability $f_{ij}$is given by

$$f_{ij} = \frac{1 - e^{-2s}}{1 - e^{-2Ns}}$$

where $N$ is population size and $s$ is the selection coefficient of the mutant relative to the resident genotype. This formulation allows both selection and drift to influence fixation. Beneficial mutations have higher fixation probabilities than neutral or deleterious mutations, but when $2Ns$ is small, drift can still permit fixation of mutations with weak effects.

To define transition probabilities between neighboring genotypes, we combined these fixation probabilities with the probabilities of individual nucleotide substitutions. Mutation probabilities were weighted according to experimentally measured *E. coli* mutation biases [52]. Thus, for each focal genotype, the probability of moving to a specific single-mutant neighbor was proportional to the product of its mutation probability and its Kimura fixation probability, normalized across all single-mutant neighbors.

Each walk terminated when it reached a local fitness peak, defined as a genotype whose single-nucleotide neighbors all had lower fitness, or after at most 25 mutational steps.

#### 8.3. Population sizes and selection strength

We performed Kimura walks for two population sizes, $N = 10^8$ and $N = 10^2$. We chose these values to contrast a regime in which drift is effectively negligible with one in which drift can substantially affect the fate of weak-effect mutations. Across our empirical landscapes, selection coefficients for individual mutational steps ranged from approximately 0.006 to 0.355 normalized regulation-strength units, with a mean of ≈0.17. For $N = 10^8$, the corresponding values of $2Ns$ are therefore extremely large, placing the system in a regime where selection dominates. For $N = 10^2$, $2Ns$ ranges from approximately 1 to 70, such that mutations with small effects are influenced substantially by drift, whereas larger-effect mutations remain primarily shaped by selection.

#### 8.4. Starting genotypes and number of walks

For the scenario in which selection favored regulation by one transcription factor, we simulated walks in both directions on the landscape of each TF pair. Specifically, when selection favored TF2, we used as starting points the ten genotypes with the highest regulation strength for TF1, and vice versa. From each starting genotype, we performed $10^4$ Kimura walks, for a total of $10^5$ walks per landscape direction.

For the scenario in which selection favored dual regulation, we used all genotypes in each library as starting points. From each starting genotype, we performed $10^4$ Kimura walks. This yielded $256 \times 10^4$, $128 \times 10^4$, and $128 \times 10^4$ walks for the CRP–Fis, CRP–IHF, and Fis–IHF landscapes, respectively.

#### 8.5. Greedy walks

To examine whether our conclusions depend on the SSWM approximation, we also simulated greedy adaptive walks [48,53–55]. In these walks, the current genotype's 1-mutant neighbor conferring the largest fitness increase is fixed at each step. Because the strongest available beneficial mutation is always fixed in a greedy walk, such walks are useful as idealized approximations to conditions in which multiple beneficial mutations compete, as can occur under clonal interference [56–58].

Because greedy walks do not use Kimura fixation probabilities, they are not a limiting case of the Kimura model. As in the Kimura walks, we only considered single-nucleotide substitutions, and terminated each walk when it had reached a local fitness peak, or after at most 25 mutational steps.

#### 8.6. Adaptive walks when selection favors dual regulation

Most analyses in the main text focused on a regime in which selection favors maximal regulation by a single transcription factor, such that the fitness optimum corresponds to the wild-type TFBS of the target TF ($S_R = 1$). Here we consider a different evolutionary scenario, in which selection favors simultaneous regulation by two transcription factors, that is, crosstalk. To examine how readily such states can evolve, we simulated adaptive walks on the experimentally mapped TFBS landscapes under the SSWM regime [23,47–49] (which we described above).

Specifically, for each TF pair, we initiated $10^4$ adaptive walks from every genotype in the corresponding TFBS library. This yielded a total of $256 \times 10^4$, $128 \times 10^4$, and $128 \times 10^4$walks for the CRP–Fis, CRP–IHF, and Fis–IHF landscapes, respectively, where 256, 128, and 128 are the numbers of TFBS variants in each library. In these simulations, we defined fitness as the sum of normalized regulation strengths for both TFs (CRP–Fis: Figure 5e–g; CRP–IHF: Supplementary Figures S30a,c,e; Fis–IHF: Supplementary Figures S30b,d,f).

Under this fitness definition, the CRP–Fis and Fis–IHF landscapes each remained single-peaked. Accordingly, all adaptive walks reached the single global optimum in each landscape.

At these optima, both TFs exhibited intermediate regulation strength $S_R$: for the CRP–Fis landscape, the peak had $S_R = 0.63$ for CRP and $S_R = 0.75$ for Fis; for the Fis–IHF landscape, the peak had $S_R = 0.68$ for Fis and $S_R = 0.76$ for IHF. By contrast, the CRP–IHF landscape contained two peaks. Of all adaptive walks on this landscape, 87% reached the first peak, where $S_R = 0.68$ for CRP and $S_R = 0.68$ for IHF, whereas 13% reached the second peak, where $S_R = 0.87$ for CRP and $S_R = 0.32$ for IHF.

Consistent with the overall smoothness of these landscapes, adaptive walks toward these crosstalk-favoring optima were nearly as short as the minimal genetic distance to the corresponding peak. For example, in the CRP–Fis landscape, the number of fixation steps along adaptive walks did not differ significantly from the minimal mutational distance to the peak (Welch two-sample t-test, $P = 0.85$; Figure 5f–g). Similar patterns occur for the CRP–IHF and Fis–IHF landscapes (Supplementary Figures S30a,c,e and S30b,d,f, respectively). These observations indicate that, even when selection favors dual regulation rather than maximal regulation by one TF, the underlying landscapes remain highly navigable.

To determine how robust this result is to population size, we repeated the simulations for small populations ($N = 10^2$), in which genetic drift is strong. In these simulations, drift had variable effects across landscapes. In the CRP–Fis landscape, it increased CRP regulation strength by 10% and decreased Fis regulation strength by 14% (Supplementary Figure S31a). In the CRP–IHF landscape, the average regulation strength associated with the more frequently reached peak was reduced by 13% for CRP and by 8% for IHF. In the Fis–IHF landscape, drift increased Fis regulation strength by 8% and reduced IHF regulation strength by 13% (Supplementary Figure S31d). These changes reflect the fact that, in small populations, drift can broaden the distribution of genotypes around a peak rather than keeping the population fixed exactly at the optimal genotype.

Genetic drift had little effect on the number of fixation steps required to reach high-fitness regions of the landscape, except in the CRP–IHF landscape. There, the average number of mutations required to reach the more frequently accessed peak increased from 3.4 to 5.0, corresponding to a 45% increase (Welch two-sample t-test, $P < 2.2 \times 10^{-16}$, $N = 256 \times 10^4$; Supplementary Figures S31e–h).

Finally, we examined deviations from the SSWM regime using greedy adaptive walks [48,53–55]. In these walks, the mutation conferring the largest fitness increase is fixed at each step, providing an approximation to evolutionary dynamics in which multiple beneficial mutations compete, as in clonal interference [56–58]. These simulations confirmed that departures from the SSWM regime do not qualitatively alter the ability of populations to evolve toward high-fitness crosstalk states, because peaks were reached with numbers of steps similar to those observed under the SSWM regime (Supplementary Figure S32).

Together, these analyses show that when selection favors dual regulation, high-crosstalk states are readily accessible on our experimentally mapped landscapes. In the single-peaked CRP–Fis and Fis–IHF landscapes, adaptive walks consistently converged on optima with substantial regulation by both TFs, whereas in the CRP–IHF landscape, walks reached one of two such optima, with a strong bias toward one of them. These outcomes are robust to increases in the strength of genetic drift and to deviations from the SSWM regime.

### Supplementary tables

**Table S1. Primers for engineering the pCAW-Sort-Seq-V2 plasmid and cloning libraries.**

| Name | Sequence | Usage |
|---|---|---|
| pCAW_frag1_F | 5’-CGTCCGACTTACGGAAGGTAGATTTTACGGC-3’ | Linearizing the pCAW-Sort-Seq fragment1 for Gibson Assembly |
| pCAW_frag1_R | 5’-CTCGTGCCTAACGGAAGGTAGATTTTACGGC-3’ | Linearizing the pCAW-Sort-Seq fragment1 for Gibson Assembly |
| pCAW_frag2_F | 5’-TAAGATTGCCACGGAAGGTAGATTTTACGGC-3’ | Linearizing the pCAW-Sort-Seq fragment2 for Gibson Assembly |
| pCAW_frag2_R | 5’-AGGCCTGACTACGGAAGGTAGATTTTACGGC-3’ | Linearizing the pCAW-Sort-Seq fragment2 for Gibson Assembly |
| Ultramer_ds_F | 5’-TTCTCAAAAGCTTCCTGCAGTATTC-3’ | Amplifying Ultramer® libraries |
| Ultramer_ds_R | 5’-CGGAAAGCACATCCGGTGAC-3’ | Amplifying Ultramer® libraries |
| TFBS_R | 5’-CCGTTTGTAGCATCACCTTC-3’ | Sequencing the TFBS region |
| pCAW_Gibs_Lib_F | 5’-GTCTGATGAGTCCGTGAGGACG-3’ | Linearizing the pCAW-Sort-Seq plasmid |
| pCAW_Gibs_Lib_R | 5’-GAGAAAGAAAACCGCCGATCCTG-3’ | Linearizing the pCAW-Sort-Seq plasmid |
| Ultramer_Gibs_F | 5’-GGTGGACAGGATCGGCGGTTTTCTTTTCTCTTCTCAAAAGCTTCCTGCAGTATTC-3’ | Amplifying Ultramer® libraries for Gibson Assembly |

| | | |
|---|---|---|
| Ultramer_Gibs_R | 5’-GGCTGTTTCGTCCTCACGGACTCATCAGACCGGAAAGCACATCCGGTG-3’ | Amplifying Ultramer® libraries for Gibson Assembly |

**Supplementary Table S2. TFBS reference sequences used in this study.**

| **Sequence Name** | **Sequence** | **TF** | **Reference** |
|---|---|---|---|
| Scrambled | TCGCCTGCTTGTAGTA | None | [59] |
| $WT_{CRP}$ | AAATGTGATCTAGATCACATTT | CRP | [60,61] |
| $WT_{Fis}$ | GCTCAAATTTTGAGC | Fis | [62–64] |
| $WT_{IHF}$ | TATCAATTTGTTG | IHF | [65,66] |

**Supplementary Table S3. Libraries used in this study.**

| **Sequence Name** | **Sequence** | **TF** | **Library size** |
|---|---|---|---|
| CRP-Fis | A**RM**T**SWR**AT**Y**T**W**GA**K**CACATTT[*] | CRP-Fis | $2^8 = 256$ |
| CRP-IHF | AAAT**R**T**S**A**WY**T**W**G**W**T**S**ACATTT[*] | CRP-IHF | $2^7 = 128$ |
| Fis-IHF | GC**YM**A**M**A**WW**TTGA**KM**[*] | Fis-IHF | $2^7 = 128$ |

[*]IUPAC nucleotide nomenclature for degenerate nucleotides: **R**: A or G; **Y**: C or T; **S**: G or C; **W**: A or T; **K**: G or T; **M**: A or C

**Table S4. Table of strains**

| **Strain** | **Genotype** | **Antibiotic resistance** | **Reference** |
|---|---|---|---|
| SIG10-MAX from Sigma Aldrich *Cloning strain* | F- mcrA Δ(mrr-hsdRMS-mcrBC) endA1 recA1 Φ80dlacZΔM15 ΔlacX74 araD139 Δ(ara,leu)7697galU galK rpsL nupG λ- tonA (StrR) | Streptomycin | Sigma Aldrich |
| *E. coli* DH5*α* *Cloning strain* | F- endA1 gln V44 thi-1 recA1 relA1 gyrA96 deoR nupG Φ80dlacZΔM15 Δ(lacZYA-argF)U169, hsdR17(rK − mK + ), λ− | None | [67] |
| *E. coli* BW25113 *Wild type strain* | Δ(araD-araB)567, ΔlacZ4787(::rrnB-3), λ−, rph-1, Δ(rhaD-rhaB)568, hsdR514 | None | [68] |
| *E. coli* JW5702-4 *Δcrp mutant strain.* | Δ(araD-araB)567, ΔlacZ4787(::rrnB-3), λ−, Δ*crp*-765::kan, rph-1, Δ(rhaD-rhaB)568, hsdR514. | Kanamycin | [68] |
| *E. coli* JW1702-1 Δ*ihfA* mutant strain. | Δ(araD-araB)567, ΔlacZ4787(::rrnB-3), λ−, ΔihfA786::kan, rph-1, Δ(rhaD-rhaB)568, hsdR514. | Kanamycin | [68] |
| *E. coli* JW3229-1 Δ*fis* mutant strain | Δ(araD-araB)567, ΔlacZ4787(::rrnB-3), λ−, Δfis-779::kan, rph-1, Δ(rhaD-rhaB)568, hsdR514. | Kanamycin | [68] |

**Table S5. Table of plasmids used in this study**

| Name | Antibiotic for plasmid selection (concentration, µg/mL) | Relevant features | Source |
|---|---|---|---|
| **pCAW-Sort-Seq-V2** | Chloramphenicol (50) | pBBR1, TetR, *sfgfp, mscarlet-I*, no TFBS, no TF | This study, derived from [1] |
| **pCAW-Sort-Seq-V2-CRPO-CRP** | Chloramphenicol (50) | pBBR1, TetR, *sfgfp, mscarlet-I,* CRP operator, *crp* | This study |
| **pCAW-Sort-Seq-V2-FisO-CRP** | Chloramphenicol (50) | pBBR1, TetR, *sfgfp, mscarlet-I,* Fis operator, *crp* | This study |
| **pCAW-Sort-Seq-V2-IHFO-CRP** | Chloramphenicol (50) | pBBR1, TetR, *sfgfp, mscarlet-I,* IHF operator, *crp* | This study |
| **pCAW-Sort-Seq-V2-N-CRP** | Chloramphenicol (50) | pBBR1, TetR, *sfgfp, mscarlet-I,* Scrambled sequence, *crp* | This study |
| **pCAW-Sort-Seq-V2-FisO-Fis** | Chloramphenicol (50) | pBBR1, TetR, *sfgfp, mscarlet-I,* Fis operator, *fis* | This study |
| **pCAW-Sort-Seq-V2-CRPO-Fis** | Chloramphenicol (50) | pBBR1, TetR, *sfgfp, mscarlet-I,* CRP operator, *fis* | This study |
| **pCAW-Sort-Seq-V2-IHFO-Fis** | Chloramphenicol (50) | pBBR1, TetR, *sfgfp, mscarlet-I,* IHF operator, *fis* | This study |
| **pCAW-Sort-Seq-V2-N-Fis** | Chloramphenicol (50) | pBBR1, TetR, *sfgfp, mscarlet-I,* Scrambled sequence, *fis* | This study |
| **pCAW-Sort-Seq-V2-IHFO-IHF** | Chloramphenicol (50) | pBBR1, TetR, *sfgfp, mscarlet-I,* IHF operator, *ihf* | This study |
| **pCAW-Sort-Seq-V2-CRPO-IHF** | Chloramphenicol (50) | pBBR1, TetR, *sfgfp, mscarlet-I,* CRP operator, *ihf* | This study |
| **pCAW-Sort-Seq-V2-FisO-IHF** | Chloramphenicol (50) | pBBR1, TetR, *sfgfp, mscarlet-I,* Fis operator, *ihf* | This study |
| **pCAW-Sort-Seq-V2-N-IHF** | Chloramphenicol (50) | pBBR1, TetR, *sfgfp, mscarlet-I,* Scrambled sequence, *ihf* | This study |
| **pCAW-Sort-Seq-V2-CF-CRP** | Chloramphenicol (50) | pBBR1, TetR, *sfgfp, mscarlet-I,* CRP-Fis library, *crp* | This study |
| **pCAW-Sort-Seq-V2-CF-Fis** | Chloramphenicol (50) | pBBR1, TetR, *sfgfp, mscarlet-I,* CRP-Fis library, *fis* | This study |
| **pCAW-Sort-Seq-V2-CI-CRP** | Chloramphenicol (50) | pBBR1, TetR, *sfgfp, mscarlet-I,* CRP-IHF library, *crp* | This study |

| | | | |
|---|---|---|---|
| **pCAW-Sort-Seq-V2-CI-IHF** | Chloramphenicol (50) | pBBR1, TetR, *sfgfp, mscarlet-I,* CRP-IHF library, *ihf* | This study |
| **pCAW-Sort-Seq-V2-FI-Fis** | Chloramphenicol (50) | pBBR1, TetR, *sfgfp, mscarlet-I,* Fis-IHF library, *fis* | This study |
| **pCAW-Sort-Seq-V2-FI-IHF** | Chloramphenicol (50) | pBBR1, TetR, *sfgfp, mscarlet-I,* Fis-IHF library, *ihf* | This study |

**Table S6. Table of sequences for CRP-Fis library regulation strength validation**

| Genotype | CRP regulation strength* | Fis regulation strength * |
|---|---|---|
| AGCTCAAATCTTGAGCACATTT | 0.0432 | 0.8483 |
| AGATCAAATTTTGAGCACATTT | 0.0932 | 0.9879 |
| AGCTCAAATCTAGAGCACATTT | 0.1161 | 0.6106 |
| AGATCAAATTTAGAGCACATTT | 0.1662 | 0.7502 |
| AACTCAAATTTTGATCACATTT | 0.2407 | 0.6512 |
| AGATCAAATCTTGATCACATTT | 0.2813 | 0.8115 |
| AGCTCAGATCTTGATCACATTT | 0.3486 | 0.6378 |
| AGCTCTAATTTTGATCACATTT | 0.3553 | 0.8235 |
| AGATGAAATTTTGATCACATTT | 0.4171 | 0.8993 |
| AGCTGTAATTTAGAGCACATTT | 0.4624 | 0.5466 |
| AAATGAGATTTTGAGCACATTT | 0.5285 | 0.4141 |
| AAATGAAATCTTGATCACATTT | 0.5561 | 0.4235 |
| AGCTGAGATCTAGATCACATTT | 0.6005 | 0.3363 |
| AGATCTGATCTTGATCACATTT | 0.6523 | 0.4739 |
| AACTCTGATCTAGATCACATTT | 0.7278 | 0.0193 |
| AAATGAGATCTAGATCACATTT | 0.7895 | 0.0009 |
| AAATGTAATCTAGATCACATTT | 0.8395 | 0.0340 |
| AGATGTGATTTAGATCACATTT | 0.8610 | 0.3240 |
| AACTGTGATCTAGATCACATTT | 0.9068 | 0.0231 |
| AAATGTGATTTAGATCACATTT | 0.9568 | 0.1654 |

* Regulation strength values measured from Sort-Seq data

**Table S7. Table of sequences for CRP-IHF library regulation strength validation**

| Genotype | CRP regulation strength* | IHF regulation strength * |
|---|---|---|
| AAATATCAATTTGTTGACATTT | 0.0024 | 0.9050 |
| AAATATCAACTTGATGACATTT | 0.0842 | 0.6339 |
| AAATATCATCTTGTTGACATTT | 0.1224 | 0.7371 |
| AAATATCATTTAGATGACATTT | 0.1915 | 0.6615 |
| AAATGTCAATTAGTTGACATTT | 0.2438 | 0.6452 |
| AAATGTCATCTTGTTGACATTT | 0.2838 | 0.3823 |
| AAATGTCATTTAGTTGACATTT | 0.3174 | 0.5276 |
| AAATATGAATTAGTTGACATTT | 0.3820 | 0.7056 |
| AAATATGATCTTGTTGACATTT | 0.4220 | 0.4426 |
| AAATATCATTTAGATCACATTT | 0.4903 | 0.6417 |
| AAATGTCAATTAGTTCACATTT | 0.5426 | 0.6255 |
| AAATGTCATCTTGTTCACATTT | 0.5826 | 0.3625 |
| AAATGTGATTTAGTTGACATTT | 0.6170 | 0.2331 |
| AAATGTGATCTAGTTGACATTT | 0.6658 | 0.1828 |
| AAATATGATCTTGTTCACATTT | 0.7207 | 0.4229 |
| AAATGTGAATTTGTTCACATTT | 0.7597 | 0.2361 |
| AAATGTGAATTAGTTCACATTT | 0.8421 | 0.3310 |
| AAATGTGAATTAGATCACATTT | 0.8776 | 0.2210 |
| AAATGTGATCTTGATCACATTT | 0.9175 | 0.0386 |
| AAATGTGATCTAGTTCACATTT | 0.9646 | 0.1630 |

* Regulation strength values measured from Sort-Seq data

**Table S8. Table of sequences for Fis-IHF library regulation strength validation**

| Genotype | Fis regulation strength* | IHF regulation strength* |
|---|---|---|
| GCCAACAAATTGAGA | 0.0306 | 0.4663 |
| GCTAACAAATTGAGA | 0.0699 | 0.6212 |
| GCCAAAAAATTGATA | 0.1187 | 0.7496 |
| GCTAACATATTGATA | 0.1587 | 1.0000 |
| GCTCACAAATTGAGA | 0.2336 | 0.3836 |
| GCTAACATTTTGATA | 0.2983 | 0.9611 |
| GCTCAAAAATTGATA | 0.3217 | 0.6669 |
| GCTCACAATTTGAGA | 0.3732 | 0.3448 |
| GCTAAAATTTTGATA | 0.4170 | 0.8951 |
| GCTCACATTTTGATA | 0.4621 | 0.7235 |
| GCTAACATATTGATC | 0.5474 | 0.8762 |
| GCTCAAATTTTGATA | 0.5808 | 0.6575 |
| GCTCAAATTTTGAGA | 0.6113 | 0.3082 |
| GCTAAAATATTGATC | 0.6661 | 0.8102 |
| GCTCACATATTGAGC | 0.7417 | 0.2892 |
| GCCAAAATTTTGAGC | 0.7970 | 0.2671 |
| GCCCACATTTTGAGC | 0.8420 | 0.0955 |
| GCTCAAATATTGAGC | 0.8604 | 0.2232 |
| GCCCAAATTTTGATC | 0.9301 | 0.3788 |
| GCTCAAATTTTGATC | 0.9694 | 0.5337 |

* Regulation strength values measured from Sort-Seq data

***Table S9. CRP-Fis landscape metrics***

| | CRP | Fis |
|---|---|---|
| Number of nodes (genotypes) | 256 | 256 |
| Number of peaks | 1 | 1 |
| Number of squares[1] | 1792 | 1792 |
| Magnitude epistasis or additivity[2,3] | 76% | 73% |
| Simple sign epistasis[3] | 0% | 4% |
| Reciprocal sign epistasis[3] | 0% | 0.3% |

[1] A square represents the connection between a focal sequence (ab) and a double mutant (AB) via two single mutants (Ab and aB).

[2] This category includes both magnitude epistasis and additivity (no epistasis) without distinguishing them, because neither of the two subcategories affects peak accessibility[33].

[3] Percentages refer to the proportion of all squares in the respective category.

***Table S10. CRP-IHF landscape metrics***

| | CRP | IHF |
|---|---|---|
| Number of nodes (genotypes) | 128 | 128 |
| Number of peaks | 1 | 1 |
| Number of squares[1] | 672 | 672 |
| Magnitude epistasis or additivity[2,3] | 80% | 60% |
| Simple sign epistasis[3] | 0% | 11% |
| Reciprocal sign epistasis[3] | 0% | 1.3% |

[1] A square represents the connection between a focal sequence (ab) and a double mutant (AB) via two single mutants (Ab and aB).

[2] This category includes both magnitude epistasis and additivity (no epistasis) without distinguishing them, because neither of the two subcategories affects peak accessibility[33].

[3] Percentages refer to the proportion of all squares in the respective category.

***Table S11. Fis-IHF landscape metrics***

| | Fis | IHF |
|---|---|---|
| Number of nodes (genotypes) | 128 | 128 |
| Number of peaks | 1 | 1 |
| Number of squares[1] | 672 | 672 |
| Magnitude epistasis or additivity[2,3] | 54% | 64% |
| Simple sign epistasis[3] | 0% | 0% |
| Reciprocal sign epistasis[3] | 0% | 0% |

[1] A square represents the connection between a focal sequence (ab) and a double mutant (AB) via two single mutants (Ab and aB).

[2] This category includes both magnitude epistasis and additivity (no epistasis) without distinguishing them, because neither of the two subcategories affects peak accessibility[33].

[3] Percentages refer to the proportion of all squares in the respective category.

### Supplementary figures

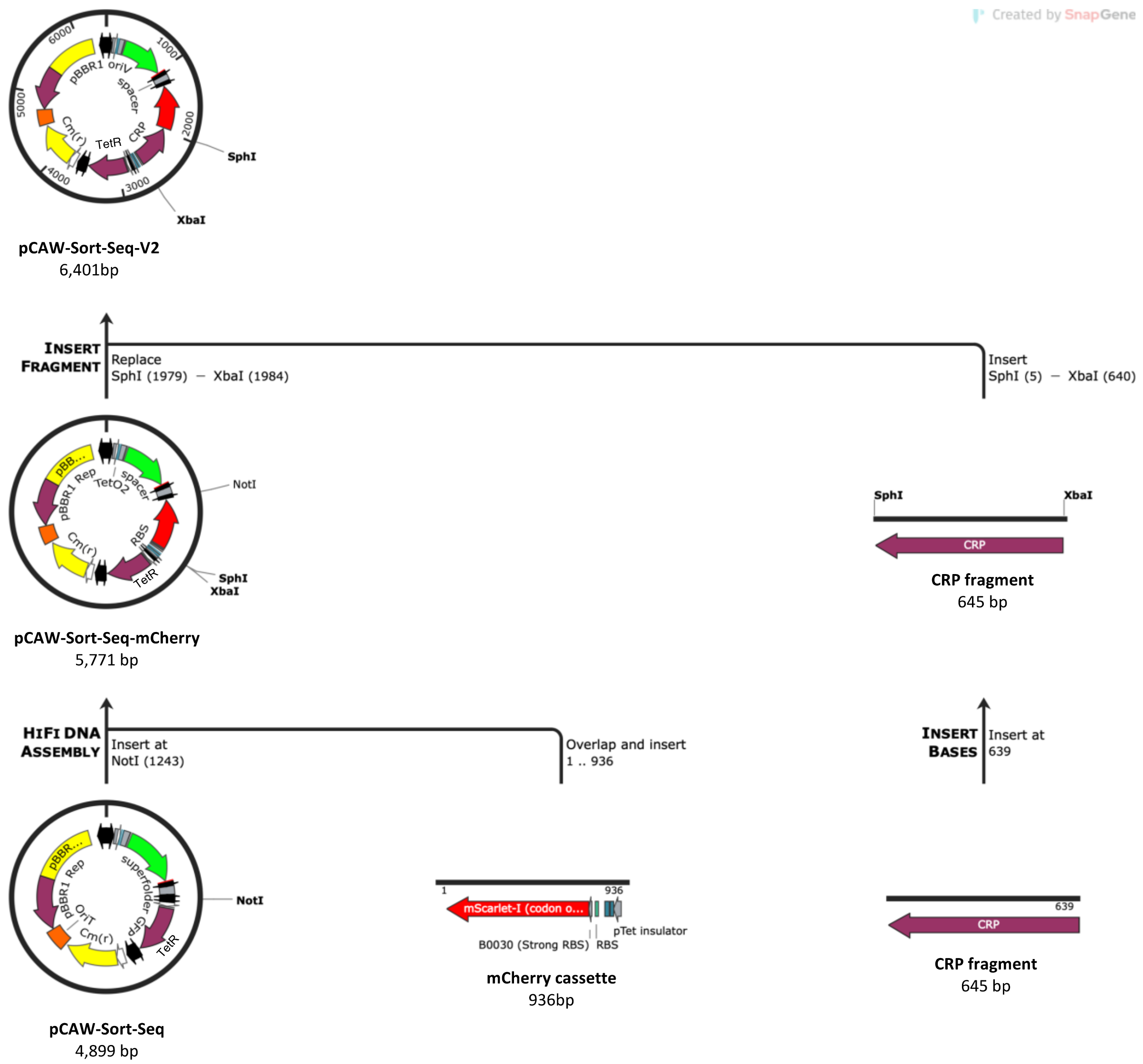


**Supplementary Figure S1. Construction of pCAW-Sort-Seq-V2.** The plasmid pCAW-Sort-Seq-V2 is a derivative of the plasmid pCAW-Sort-Seq[1]. The figure illustrates the sequential cloning steps involved in the plasmid's construction, detailed from the bottom upwards. For an in-depth explanation, see **Supplementary Methods 2-4**. Briefly, we began with the synthesis and cloning of a codon-optimized *mscarlet* gene expression cassette into the pCAW-Sort-Seq plasmid, oriented antiparallel to the tetracycline resistance gene (*tetr*). This generated the intermediate pCAW-Sort-Seq-mscarlet plasmid. Subsequently, we inserted each TF gene, exemplified by the *crp* gene for the purpose of this representation, upstream of the *mscarlet* gene in a bicistronic operon configuration. This step created the TF-specific expression plasmid variants (for CRP, Fis and IHF). The final stage, not depicted here, involved the incorporation of the appropriate wild-type TFBSs and libraries into each plasmid variant upstream of the *gfp* gene, which yielded the plasmids we utilized in our experiments.

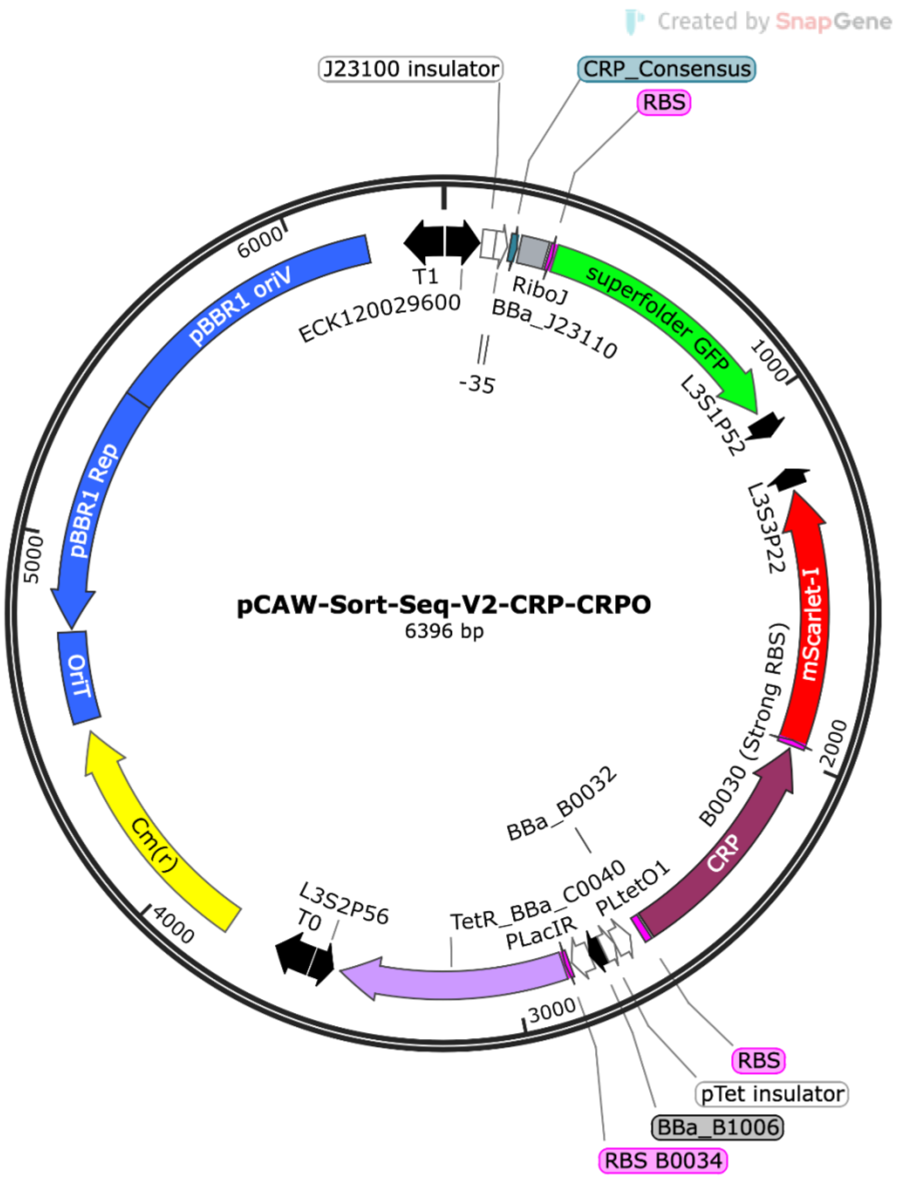


**Supplementary Figure S2. Components of plasmid pCAW-Sort-Seq-V2.** The plasmid system **pCAW-Sort-Seq-V2** is a derivative of the plasmid pCAW-Sort-Seq[1]. It encodes the broad-host, low-copy number replication origin pBBR1 [4] (5 to 10 copies per cell), together with an origin of transference (OriT) for conjugation (blue), and a chloramphenicol resistance gene (yellow). The interchangeable regulatory region where the TFBS is located (cyan) is placed between a constitutive promoter (BBa_J23110487) and a superfolder GFP (*sfgfp*, green) fluorescent reporter gene[6]. The transcriptional insulator RiboJ[7] is located upstream of the *sfgfp* gene. The *tetr* gene (in purple) is derived from the original Tn10 transposon[70–72] under the control of a low-strength constitutive promoter (pLac promoter variant developed in ref. [73]). TetR regulates the expression of a bicistronic operon encoding the TF of interest (represented by CRP, dark purple) and an mScarlet-I red fluorescent reporter[9] (red). Promoters and their respective insulators[74] are represented as white arrows. Transcriptional terminators are represented as black arrows[75]. Ribosome binding sites are represented by pink arrows.

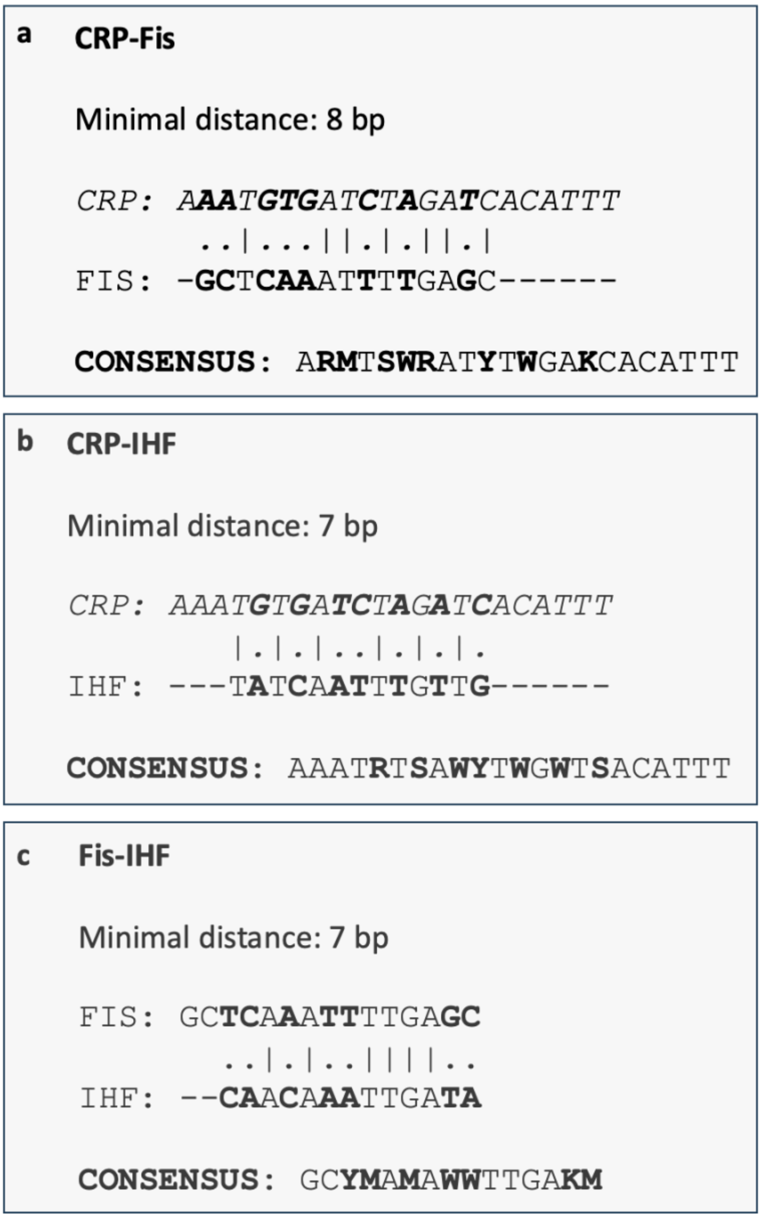


**Supplementary Figure S3. Library design.** Each panel represents the sequence composition of one of our three TFBS libraries, as indicated in the panel title. To obtain the data in each panel, we aligned strong wild-type TFBSs for each TF in a pairwise manner using a Smith-Waterman local alignment algorithm[10] without indels. Dots and vertical bars represent nucleotide mismatches and matches between TFBSs. Consensus sequences used for library construction are represented using the following IUPAC nucleotide nomenclature for degenerate nucleotides: **R**: A or G; **Y**:C or T; **S**: G or C; **W**: A or T; **K**:G or T; **M**:A or C. **a. CRP-Fis Library. b. CRP-IHF Library. c. Fis-IHF Library.**

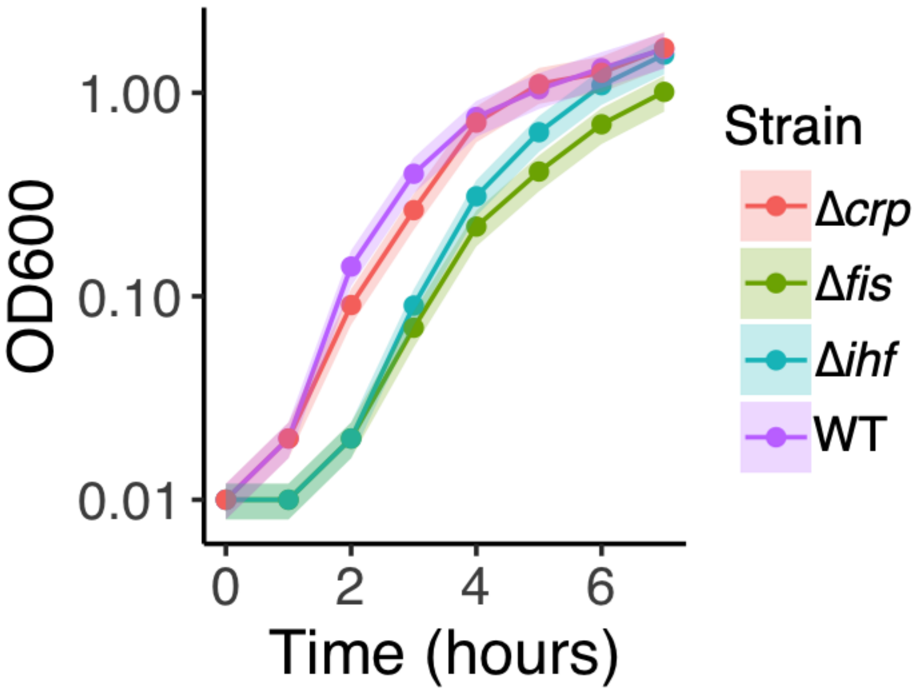


**Supplementary Figure S4. Growth profile for strains used in this study.** Growth profiles for all strains used in this study are represented as optical densities ($OD_{600}$, y-axis) measured over 7 hours of growth at 1-hour intervals (x-axis). We transformed all strains with the plasmid pCAW-Sort-Seq-V2 and grew them in LB medium without inducers. Colors (legend) represent the different strains. Wild-type (purple) refers to the BW25113 parent-strain from which the mutants (Δ*crp*, Δ*fis*, Δ*ihf*) were derived. Each circle represents the mean of three biological replicates, based on independently (plate-reader) measured changes in $OD_{600}$ for each replicate. Shaded areas correspond to one standard deviation over the three replicates. Note that the Δfis and Δihf deletion strains show a modest increase in lag time relative to wild type, consistent with the known pleiotropic roles of Fis and IHF as global regulators. However, all strains reach comparable final densities, and growth differences were not statistically significant (Welch t-test, $p > 0.05$ for all pairwise comparisons at the 7-hour endpoint), confirming that these single deletions do not cause strong fitness defects under the LB growth conditions used in this study.

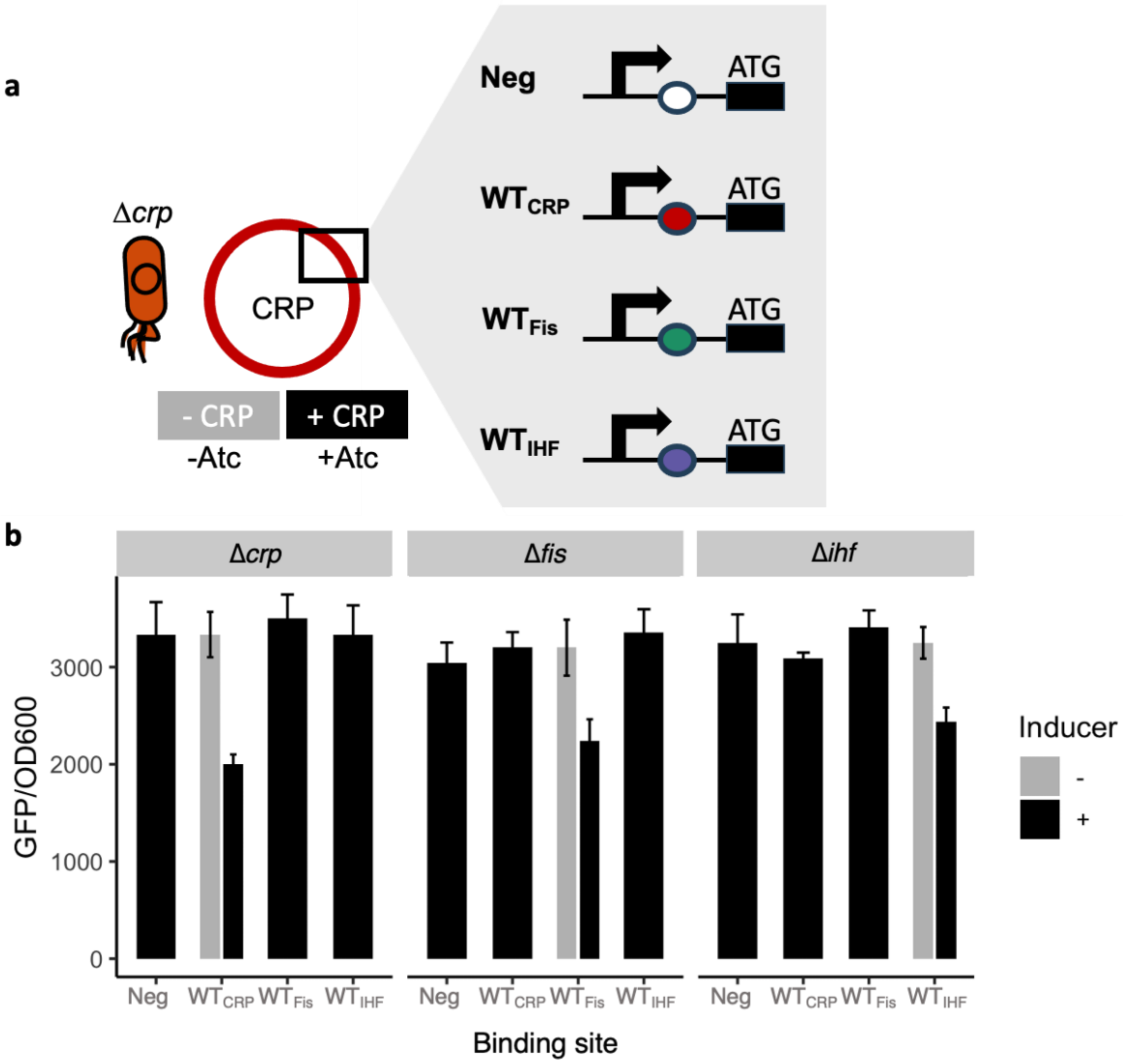


**Supplementary Figure S5. Combinatorial characterization of the wild-type TFBSs in all mutant *E. coli* strains of this study.** The figure shows how we characterized the regulatory interactions between our wild-type TFBSs and the three TFs used in this study (CRP, Fis and IHF). **a. Expression plasmids and bacterial strains.** Our approach involved the creation of three fundamental variants of our reporter plasmids, each tailored to express one of the transcription factors. These plasmids are called pCAW-CRP, pCAW-Fis, and pCAW-IHF. For each of these plasmids, we further developed four derivatives by introducing different TFBSs upstream of the GFP-coding gene (**see Figure 1, Supplementary Figure S1, and grey inset in this figure**). For clarity, this panel focuses on pCAW-CRP and its four derivatives. The plasmid pCAW-CRP is visually represented as a red circle. The TF's state of expression is indicated by two different colored boxes: a grey box symbolizing the uninduced state of transcription factor expression ("-CRP"), and a black box symbolizing the induced state ("+CRP"). These expression states are controlled by the inducer Atc whose absence ("-Atc") or presence ("+Atc") translates to the absence or presence of the TF. Additionally, the regulatory region of the plasmid, crucial for understanding gene expression dynamics, is highlighted in the black square and further enlarged in a grey inset for detailed examination. In this inset, each of the four derivatives of the pCAW-CRP plasmid is showcased with three basic elements (from left to right): the constitutive promoter[69] (black arrow), the TFBS (colored circles), and the start codon ("ATG") of the GFP gene. These derivatives are distinguished by their unique TFBSs. The first derivative, termed Negative ("Neg"), contains a non-functional, scrambled TFBS [59], indicated as a white circle, which does not bind to any of the studied transcription factors[59]. The subsequent three derivatives harbor TFBSs that bind strongly to CRP ("$WT_{CRP}$", red), Fis ("$WT_{Fis}$", green), and IHF ("$WT_{IHF}$", purple). We introduced each of these plasmid constructs into a mutant *E. coli* strain that lacks the gene encoding the corresponding TF (CRP, Fis, or IHF). The figure highlights only one of these strains, i.e., the *crp* mutant strain denoted as Δ*crp* and represented as the red *E. coli* cell on the far left of the figure. Overall, we thus constructed 4 (TFBSs) x 3 (TF-plasmids) = 12 reporter plasmids for our experiments (**Supplementary methods 2-3, Supplementary Table S5**). **b. The wild-type TFBS sequences are TF-specific and do not crosstalk.** The bar plots represent the fluorescence levels (x-axis) associated with each aforementioned reporter plasmid (y-axis), and measured in the three mutant *E. coli* strains (Δ*crp*, Δ*fis* and Δ*ihf*, left, middle and right panels). We recorded these fluorescence levels under two distinct conditions: in the presence of the inducer Atc (indicating the expression of the plasmid-encoded TF, black bars) and in its absence (no expression of the TF, grey bars). The panel shows both mean values (bars) and one standard deviation (whiskers), as derived from three independent endpoint plate reader measurements (GFP fluorescence normalized by $OD_{600}$, arbitrary units) after 12 hours of growth.

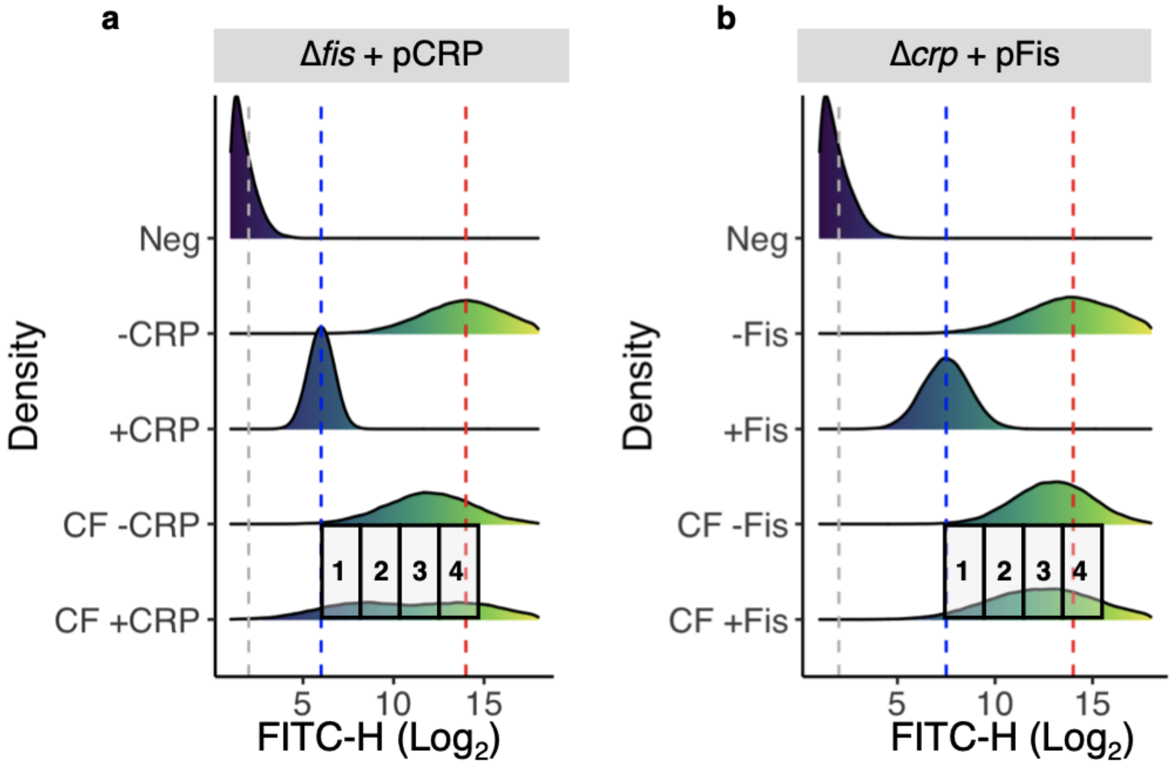


**Supplementary Figure S6. Distribution of fluorescence levels for controls and CRP-Fis library under CRP or Fis expression.** Each density plot represents the distribution of GFP fluorescence for 100,000 cells. Specifically, the horizontal axis shows GFP fluorescence (FITC-H values, arbitrary units, $\log_2$ scale), while each histogram indicates the relative frequency of cells with a given fluorescence value. Heatmap colors indicate fluorescence, from low (dark purple) to high (yellow). The vertical axis indicates the five experimental conditions under which we measured fluorescence: 1) negative control ("Neg"), without GFP expression (promoterless pCAW-v2); 2) positive control with the wild-type TFBS cloned in the pCAW-v2 without induction of the TF ("-TF", i.e., absence of the inducer Atc; note the TF gene is present on the plasmid but not expressed); 3) the same positive control with the expression of the cognate TF ("+TF", inducer Atc present); 4) CRP-Fis library in pCAW-v2 without TF expression ("CF -TF") and 5) CRP-Fis library in pCAW-v2 with TF expression ("CF +TF"). We performed density smoothing using a Gaussian kernel function to create a smooth density plot for each histogram with the aid of the *ggplot2* package[76]. Vertical dashed lines mark thresholds for binning cells harboring TFBS variant library: a grey line for the upper limit of cell autofluorescence (negative control), a blue line for the geometric mean of the positive control with TF (indicating strong binders), and a red line for the geometric mean of the positive control without TF (indicating absence of binders). Rectangular boxes (1 to 4) represent bins/gates for each sample, based on the aforementioned thresholds and spaced equally on a binary logarithmic ($\log_2$) scale. **a. CRP-Fis library analysed for CRP binding.** See **Figure 2a** in the main text for more specific details about the experimental design to dissect TF-TFBS interactions. **b. CRP-Fis library analysed for Fis binding.** The grey box on top of each panel shows the combination of the strain (left) and the TF (right) expressed in the plasmid pCAW-v2.

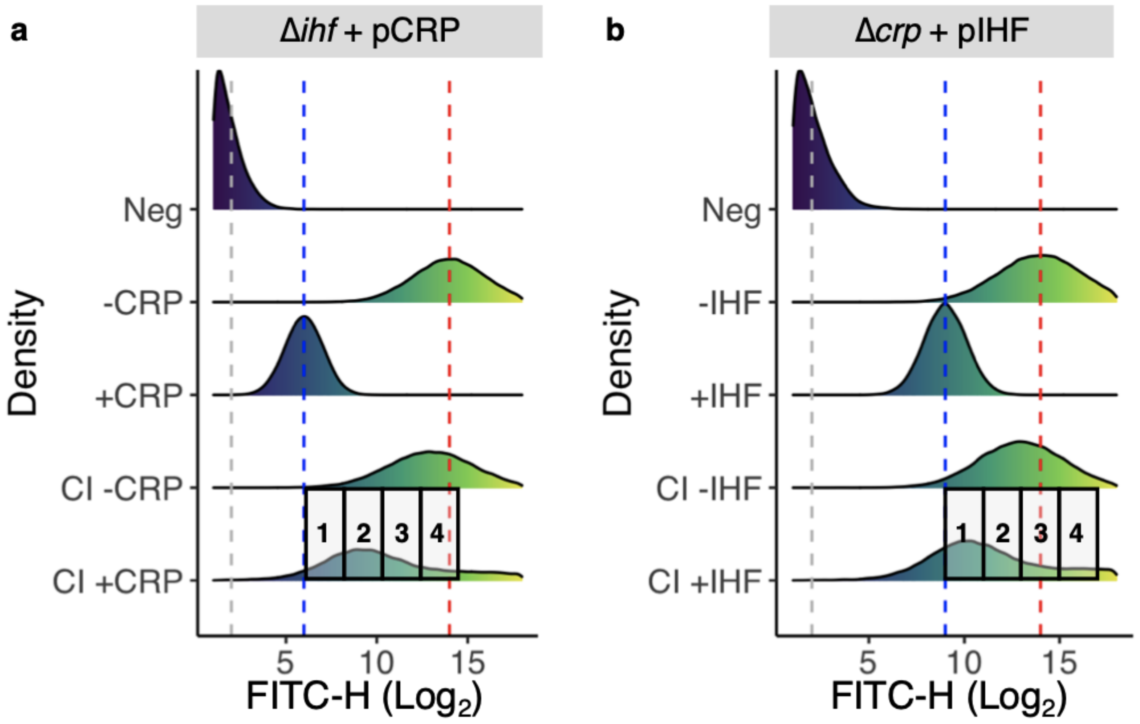


**Supplementary Figure S7. Distribution of fluorescence levels for controls and CRP-IHF library under CRP or IHF expression.** Each density plot represents the distribution of fluorescence levels for 100,000 cells. The horizontal axis shows GFP fluorescence values in FITC-H (arbitrary units, log2 scale), while the vertical axis indicates the relative frequency of each fluorescence value. Heatmap color gradients also indicate FITC-H values, from low (dark purple) to high (yellow). Samples, from top to bottom: 1) negative sample ("Neg"), without GFP expression (promoterless pCAW-v2); 2) positive control with the wild-type TFBS cloned in the pCAW-v2 without induction of the TF ("-TF", i.e., absence of the inducer Atc; note the TF gene is present but not induced); 3) the same positive control with the expression of the cognate TF ("+TF", presence of the inducer Atc); 4) CRP-IHF library in pCAW-v2 without TF expression ("CI -TF") and 5) CRP-IHF library in pCAW-v2 with TF expression ("CI +TF"). We performed density smoothing using a Gaussian kernel function to create a smooth density plot – *ggplot2* package[76]. Vertical dashed lines mark thresholds for binning the TFBS variant library: a grey line for the upper limit of cell autofluorescence (negative control), a blue line for the geometric mean of the positive control with TF (indicating strong binders), and a red line for the geometric mean of the positive control without TF (indicating absence of binders). Rectangular boxes (1 to 4) represent bins/gates for each sample, based on the aforementioned thresholds and spaced equally on a $\log_2$ scale. **a.CRP-IHF library analysed for CRP binding.** Note that the grey box on top represents the combination of strain (Δ*ihf*) and the TF expressed in the pCAW-v2 plasmid (CRP). Refer to **Figure 2a** on the main text for more specific details about the experimental design for dissecting TF-TFBS interactions. **b.CRP-IHF library analysed for IHF binding.** Note that the grey box on top represents the combination of strain (Δ*crp*) and the TF expressed in the pCAW-v2 plasmid (IHF).

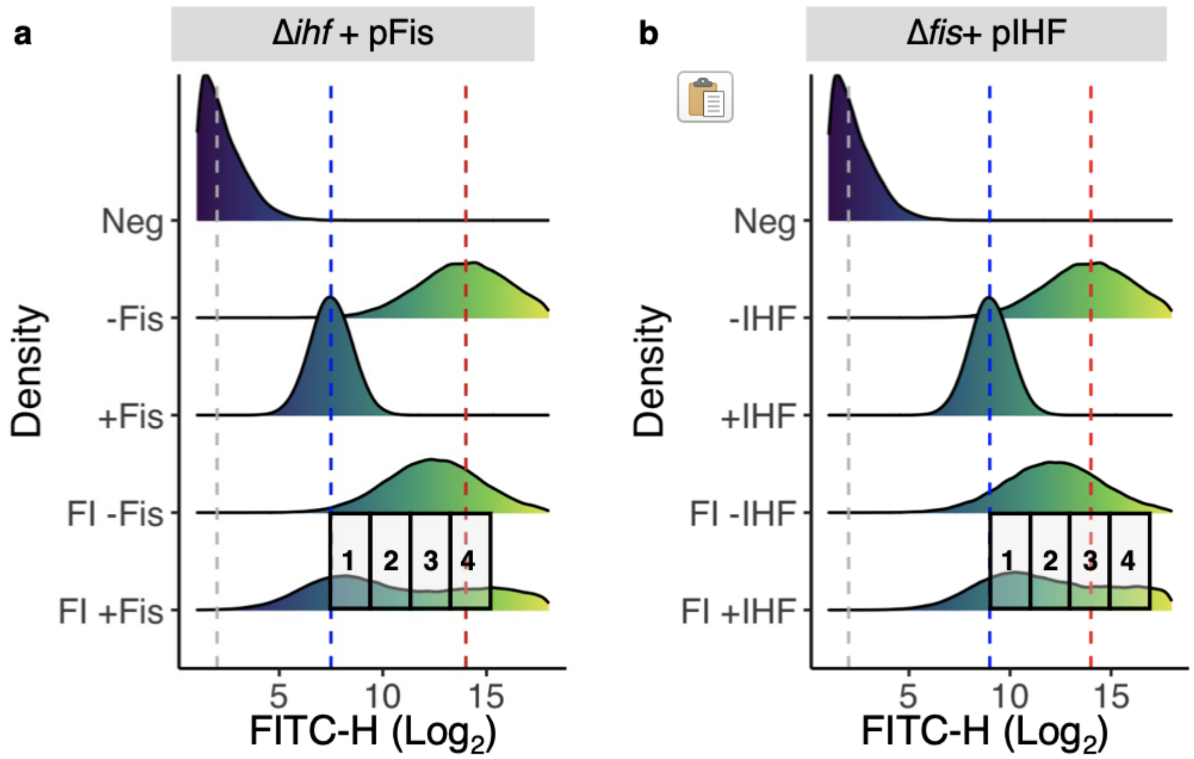


**Supplementary Figure S8. Distribution of fluorescence levels for controls and Fis-IHF library under Fis or IHF expression.** Each density plot represents the distribution of fluorescence levels for 100,000 cells. The horizontal axis shows GFP fluorescence values in FITC-H (arbitrary units, $log_2$ scale), while the vertical axis indicates the relative frequency of each fluorescence value. Heatmap color gradients also indicate FITC-H values, from low (dark purple) to high (yellow). Samples, from top to bottom: 1) negative sample ("Neg"), without GFP expression (promoterless pCAW-v2); 2) positive control with the wild-type TFBS cloned in the pCAW-v2 without the expression of its cognate TF ("-TF", absence of the inducer Atc); 3) the same positive control with the expression of the cognate TF ("+TF", presence of the inducer Atc); 4) Fis-IHF library in pCAW-v2 without TF expression ("FI -TF") and 5) Fis-IHF library in pCAW-v2 with TF expression ("FI +TF"). We performed density smoothing using a Gaussian kernel function to create a smooth density plot – *ggplot2* package[76]. Vertical dashed lines mark thresholds for binning the TFBS variant library: a grey line for the upper limit of cell autofluorescence (negative control), a blue line for the geometric mean of the positive control with TF (indicating strong binders), and a red line for the geometric mean of the positive control without TF (indicating absence of binders). Rectangular boxes (1 to 4) represent bins/gates for each sample, based on the aforementioned thresholds and spaced equally on a $log_2$ scale. **a. Fis-IHF library analysed for Fis binding.** Note that the grey box on top represents the combination of strain (Δ*ihf*) and the TF expressed in the pCAW-v2 plasmid (Fis). Refer to **Figure 2a** on the main text for more specific details about the experimental design for dissecting TF-TFBS interactions. **b. Fis-IHF library analysed for IHF binding.** Note that the grey box on top represents the combination of strain (Δ*fis*) and the TF expressed in the pCAW-v2 plasmid (IHF).

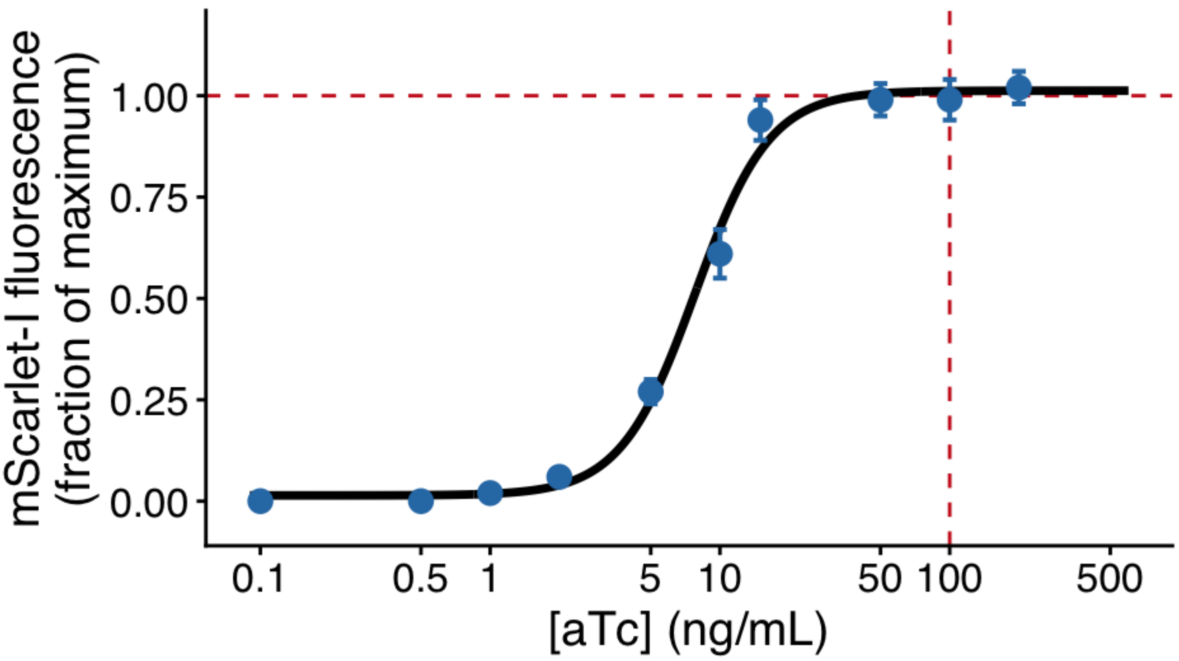


**Supplementary Figure S9. Calibration of Atc-dependent mScarlet-*I* expression.** Circles show mean mScarlet-I fluorescence ± SD from three independent endpoint plate reader measurements for ten Atc concentrations (0–200 ng/mL), normalized to the maximum fluorescence observed at saturating Atc. The black curve is a Hill function fit ($EC_{50}$ = 8 ng/mL, Hill coefficient n = 2.1; parameters from Lutz & Bujard, 1997[5] and Baumschlager et al., 2020[77]). Red dashed lines indicate the Atc concentration used throughout this study (100 ng/mL), which corresponds to ~98% of maximum induction, confirming that transcription factor expression is effectively saturated under our experimental conditions.

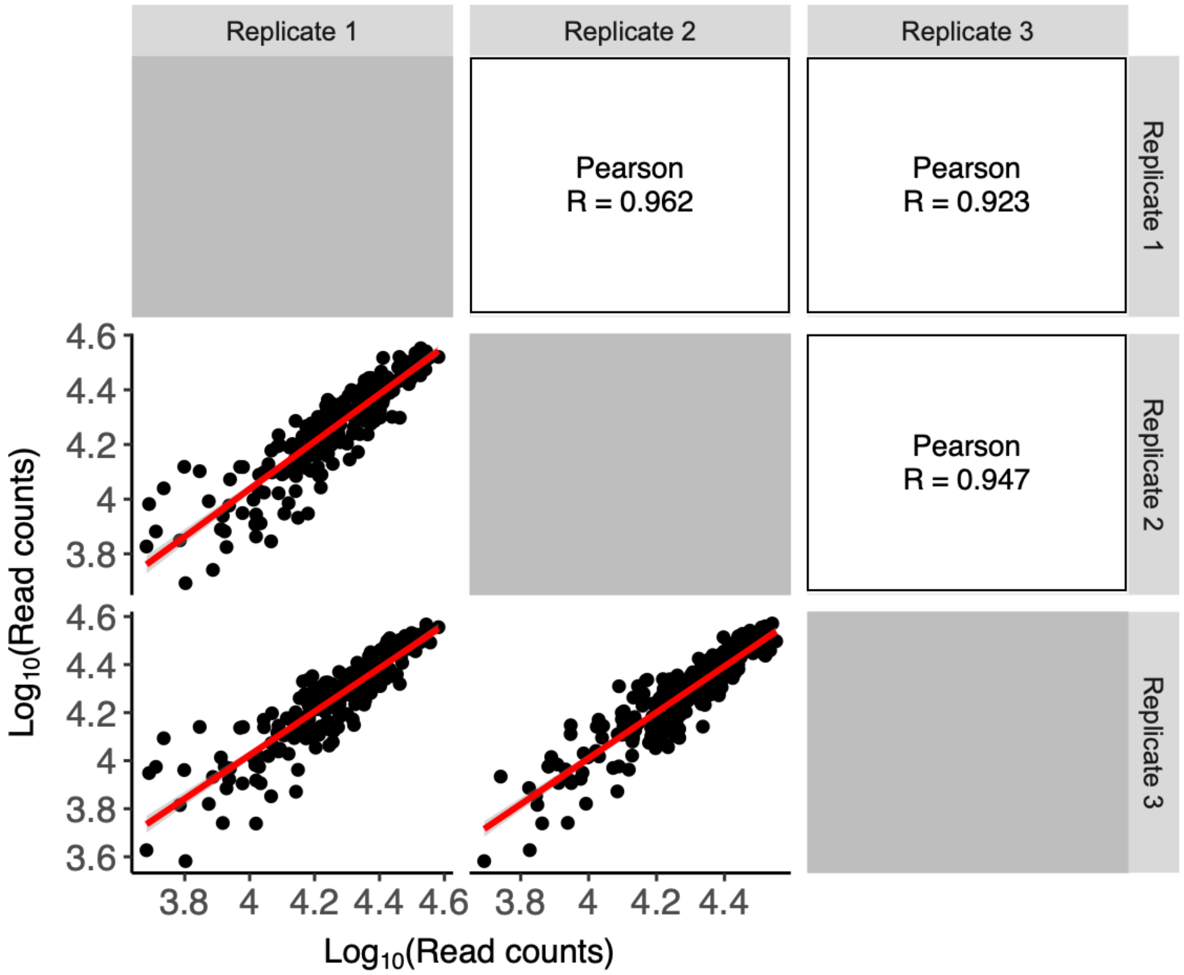


**Supplementary Figure S10. Pairwise associations of sequencing read counts for replicates of the CRP-Fis library.** Sequence read counts are shown as scatter plots in the lower panels, and their Pearson correlation coefficients R are shown in each corresponding upper panel. Red lines are linear regression lines, and red shading around them indicates 95% confidence intervals. Note the logarithmic scale in all data panels.

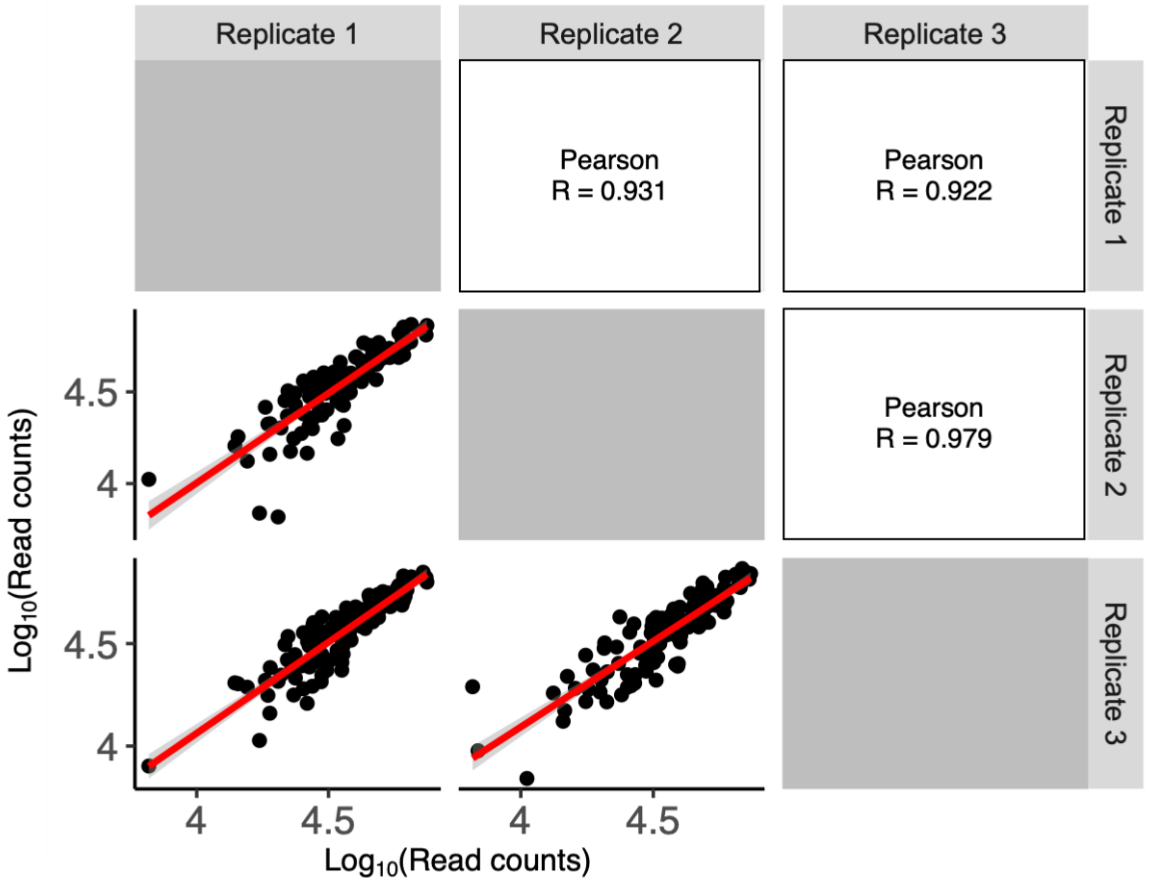


**Supplementary Figure S11. Pairwise associations of sequencing read counts for replicates of the CRP-IHF library.** Sequence read counts are shown as scatter plots in the lower panels, and their Pearson correlation coefficients R are shown in each corresponding upper panel. Red lines are linear regression lines, and red shading around them indicates 95% confidence intervals. Note the logarithmic scale in all data panels.

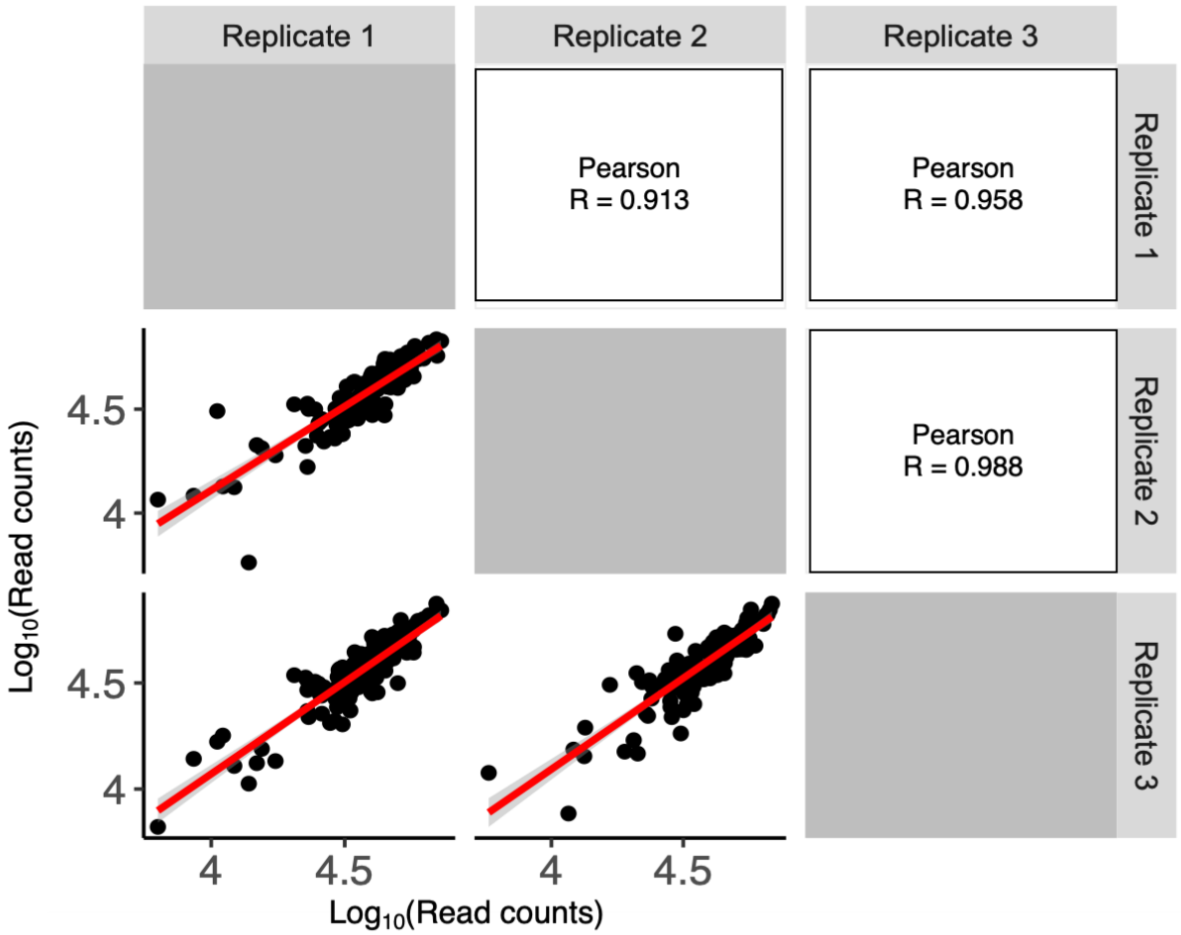


**Supplementary Figure S12. Pairwise associations of sequencing read counts for replicates of the Fis-IHF library.** Sequence read counts are shown as scatter plots in the lower panels, and their Pearson correlation coefficients R are shown in each corresponding upper panel. Red lines are linear regression lines, and red shading around them indicates 95% confidence intervals. Note the logarithmic scale in all data panels.

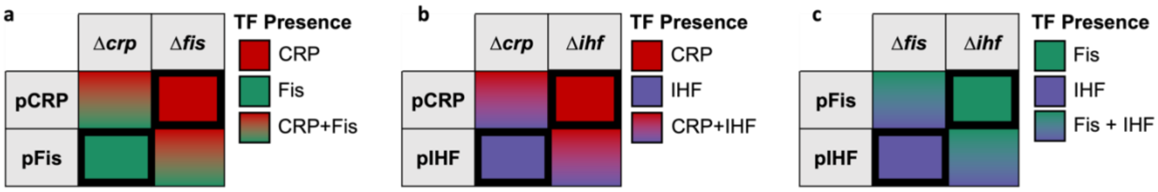


**Supplementary Figure S13. a-c. Specific plasmid-strain combinations ensure the presence of only one of two TFs during our measurements.** Each table schematically represents the intracellular presence of TFs for combinations of a reporter plasmid encoding one of the TFs (rows) transformed into a mutant *E. coli* strain lacking the gene encoding one of the TFs (columns). Pure colors represent the presence of a single TF in the cell (CRP: red; Fis: green, IHF: purple). Color gradients represent the presence of both TFs. We avoided plasmid-strain combinations in which both TFs were present in a cell, because such combinations did not allow us to unambiguously measure the regulation conveyed by one TF's binding to each of the TFBSs in an exaptation landscape. The plasmid-strain combinations we used are highlighted with thick black borders.

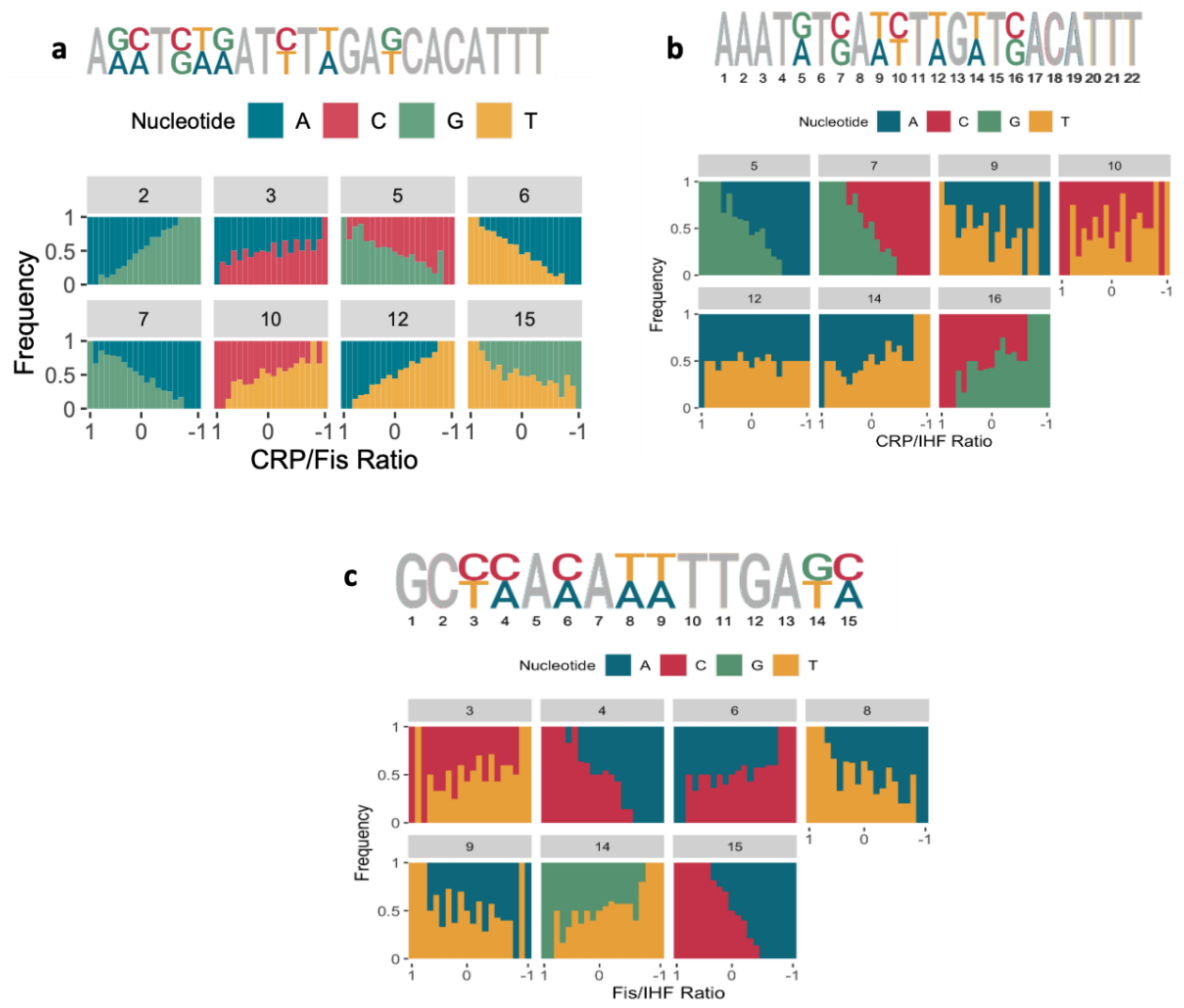


**Supplementary Figure S14. Effects of TF1 and TF2 regulation strength ratios on TFBS nucleotide composition.** The nucleotide composition at specific binding site positions would undergo pronounced changes during the evolution of a binding site for a transcription factor TF2 from that for a transcription factor TF1. To demonstrate this, we binned transcription factor binding sites with different ratios of regulation strength to TF1 and TF2 into 20 equally-spaced intervals of this ratio, and aligned the binding sites in each such bin. This alignment allowed us to compute the frequency of each nucleotide at a given location in a binding site. **a. Shifts in nucleotide composition at mutated binding site positions during shifts in the ratio between regulation strengths for CRP and Fis.** The top of the panel shows a sequence logo with conserved (grey) and variable (colored) sites in our CRP-Fis library. A sequence logo consists of a stack of letters at each position of a DNA sequence, where the relative sizes of the letters indicate the frequency of the corresponding nucleotide in the sequence. Each of the seven panels below this logo represents data from one nucleotide position (numbered in the grey box above the panel), with a stacked bar plot indicating the frequency of specific color-coded nucleotides (vertical axes) for all binding sites with a given ratio of regulation strength to CRP and Fis, i.e., for all binding sites in one of the 20 regulation strength bins (horizontal axes). **b. Like a, but for TFs CRP and IHF. c. Like a, but for TFs Fis and IHF.**

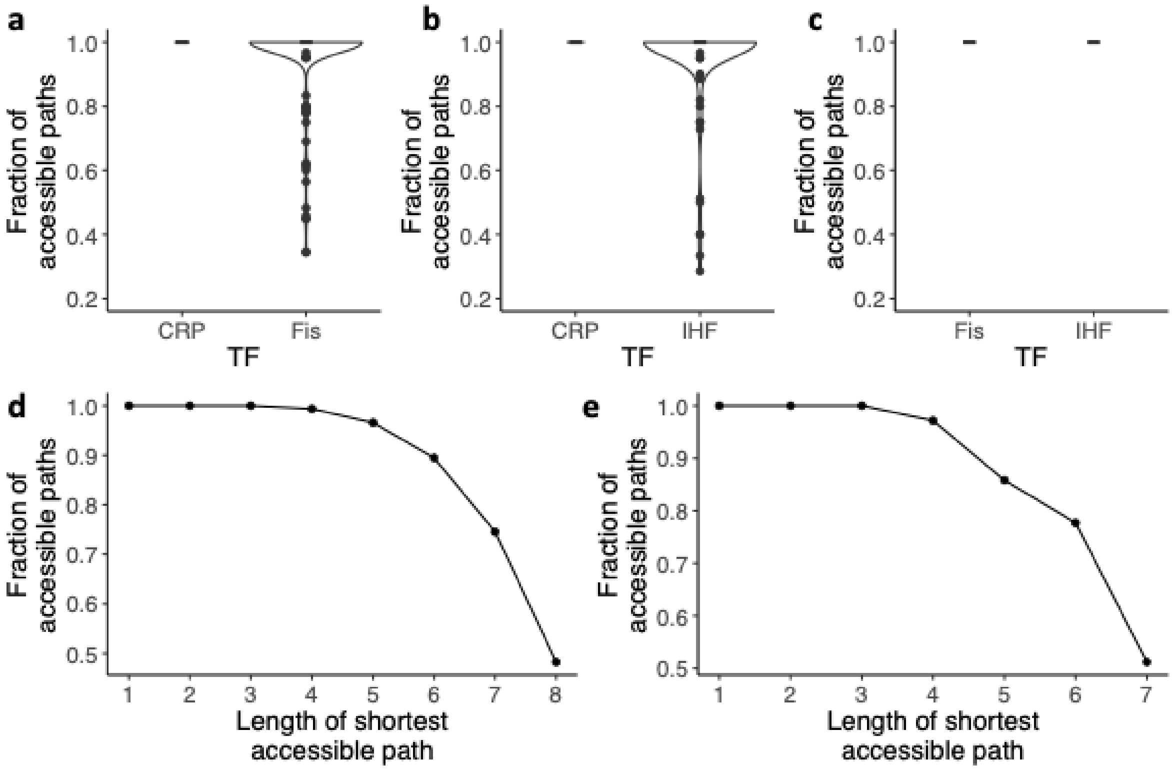


**Supplementary Figure S15. Most paths to peaks are accessible across landscapes. a-c.** Distribution of the fraction of accessible paths **(i.e., the number of accessible paths divided by the total number of all paths accessible and inaccessible) from each non-peak genotype sequence to the peak**. The violin plots display the distribution of the fraction of accessible paths (the fraction of accessible paths relative to all existing paths) for each genotype in each landscape, with the shape of each violin representing a kernel density estimation. Black dots outside the main body of the violin plot represent outliers, defined as data points that fall outside the span of 1.5 times the interquartile range (IQR), which is the range between the first quartile (25th percentile) and the third quartile (75th percentile), capturing the middle 50% of the data. **a. CRP-Fis exaptation landscape.** N = 256 genotypes. Only the Fis landscape harbors genotypes with inaccessible paths to its peak. The total fraction of accessible paths in the landscape, represented as the total number of paths that can reach a peak and divided by the total number of paths (accessible and inaccessible) to the peak, is 0.65, meaning that only 65% of all paths (91,619/140,495) are accessible. **b. CRP-IHF landscape.** N = 128 genotypes. Only the IHF landscape contains genotypes with inaccessible paths to its peak. The total fraction of accessible paths in the landscape is 0.69, meaning that only 69% of all paths (10,202/14,780) are accessible. **c. Fis-IHF landscape.** N = 128 genotypes. All paths are accessible for both Fis and IHF. **d-e. The fraction of accessible paths decreases as path length increases.** The vertical axis displays the fraction of accessible paths (among all possible shortest paths) reaching the high peak. As path length increases, the proportion of accessible paths diminishes. Circles represent the average fraction of accessible paths for each path length. **d. Paths towards the Fis peak in the CRP-Fis landscape. e. Paths towards IHF in the CRP-IHF landscape**

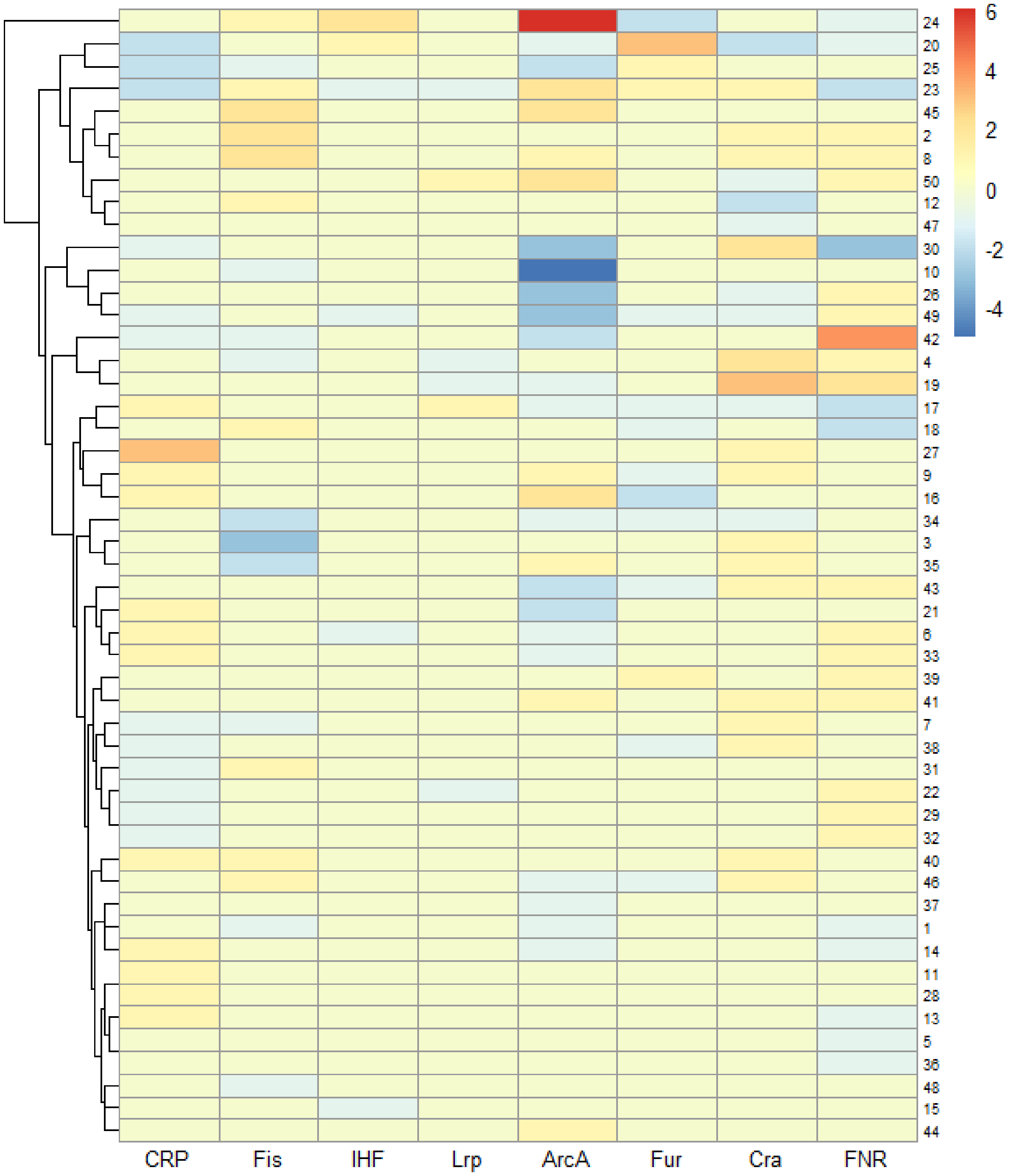


**Supplementary Figure S16. TFBS shifts for global regulators in 50 conserved intergenic regions between *Salmonella typhimurium* and *E. coli.*** The analysis of 50 conserved intergenic regions, selected from 991 orthologous intergenic regions, is visualized in a clustered heatmap. In this heatmap, each row corresponds to an intergenic region, and each column corresponds to a transcription factor. The color scale indicates the change in the number of TFBSs after subtracting the binding site matrix (**Supplementary Methods S7.7**) of *Salmonella* from that of *E. coli*. Warmer colors denote a higher number of TFBSs in *Salmonella* compared to *E. coli*, while cooler colors indicate a lower number of TFBSs in *Salmonella* relative to *E. coli*. See **Supplementary Methods 7.7** for further details on this bioinformatic analysis. We classified a TFBS change as a candidate exaptation event when three conditions were met: (i) a TFBS for TF1 was present in the *E. coli* intergenic region but absent from the aligned Salmonella region; (ii) a TFBS for a different TF2 was present in the *Salmonella* region but absent from the aligned E. coli region; and (iii) the TF1 site in E. coli and the TF2 site in Salmonella overlapped by at least 1 bp in the global alignment. Rows showing changes for only one TF are consistent with lineage-specific gain or loss of TFBSs. Rows showing the loss of a site for one TF together with the gain of a site for another TF at the same aligned position are consistent with exaptive turnover. A worked example of a candidate exaptation event is described in **Supplementary Methods 7.7**.

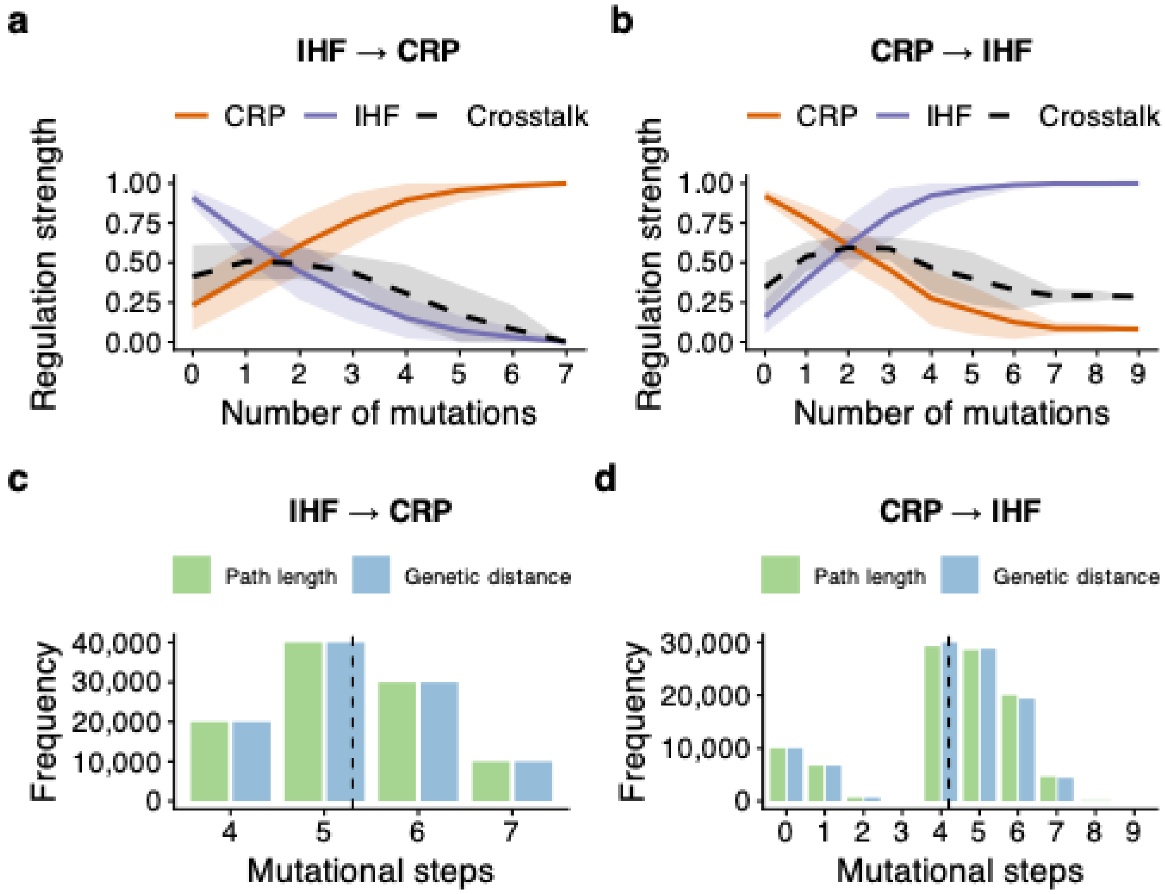


**Supplementary Figure S17. Rapid exaptation on the CRP-IHF landscape under the SSWM regime.** All data presented are based on Kimura adaptive walks with populations of $10^8$ individuals. Data for each plot is based on $10^5$ simulated adaptive walks, i.e., $10^4$ adaptive walks for each of the 10 starting genotypes with the highest regulation strength for the opposite TF. **a-b. All adaptive walks readily attain adaptive peaks.** Each plot shows how regulation strength (vertical axis) for CRP (orange) and IHF (purple) changes as a function of path length traversed (horizontal axis) from the initial sequence to the reached peak during adaptive walks. Crosstalk along the walks is shown by the black dashed line and is defined as the geometric mean of CRP and IHF regulation strengths. Curves represent averages over all $10^5$ adaptive walks, and shading indicates one standard deviation of regulation strengths. **a. All adaptive walks attain the CRP peak.** The data is based on $10^4$ walks starting from each of the 10 strongest binding sites for IHF, favoring increasing CRP binding. **b. All adaptive walks attain the IHF peak.** Like panel (a), but for adaptive walks starting from the 10 strongest binding sites for CRP, favoring increasing IHF binding. **c-d. Evolutionary paths to a peak are not much longer than minimal genetic distances.** Each grouped bar chart shows the distribution of path lengths (green) and genetic distances (blue) for all pairs of starting (non-peak) and ending (peak) genotypes during $10^5$ adaptive walks. The black dashed vertical line represents the mean of both distributions, which is not significantly different between distributions. We used a t-test of the null hypothesis that the means of these distance distributions are statistically indistinguishable. **c. Distribution of accessible path lengths and genetic distances to the CRP peak.** We observed no significant difference between the means of genetic distance and path length (mean ± s.d.: 5.3 ± 0.9; Welch Two Sample t-test: t = 35.113, df = 196,700, P-value= 0.82, $N_1 = 10^5$ genetic distances, $N_2 = 10^5$ path lengths). **d. Distribution of accessible**

**path lengths and genetic distances to the IHF peak.** We observed no significant difference between mean genetic distance and path length (4.1±0.9, mean ± s.d., Welch Two Sample t-test, t = 31.973, df = 196,700, P-value= 0.62, $N_1 = 10^5$ genetic distances, $N_2 = 10^5$ path lengths).

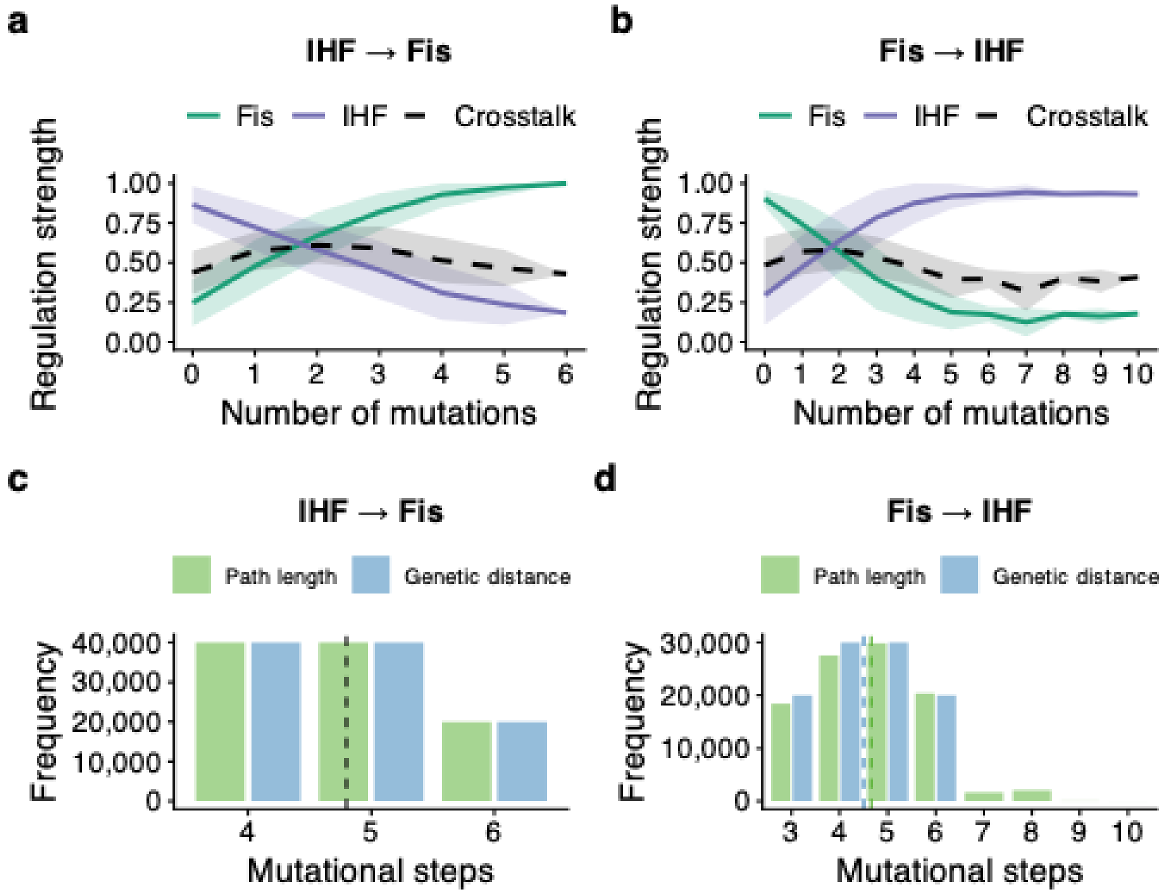


**Supplementary Figure S18. Rapid exaptation on the Fis-IHF landscape under the SSWM regime.** All data presented are based on Kimura adaptive walks with populations of $10^8$ individuals. Data for each plot is based on $10^5$ simulated adaptive walks, i.e., $10^4$ adaptive walks for each of the 10 starting genotypes with the highest regulation strength for the opposite TF. **a-b. All adaptive walks readily attain peaks.** Each plot shows how regulation strength (vertical axis) for Fis (green) and IHF (purple) changes as a function of path length traversed (horizontal axis) from the initial sequence to the reached peak during adaptive walks. Crosstalk along the walks is shown by the black dashed line and is defined as the geometric mean of Fis and IHF regulation strengths. Curves represent averages over all $10^5$ adaptive walks, and shading indicates one standard deviation of regulation strengths. **a. All adaptive walks attain the Fis peak.** The data is based on $10^4$ walks starting from each of the 10 strongest binding sites for IHF, favoring increasing Fis binding. **b. All adaptive walks attain the IHF peak.** Like panel (a), but for adaptive walks starting from the 10 strongest binding sites for Fis, favoring increasing IHF binding. **c-d. Evolutionary paths to a peak are not much longer than minimal genetic distances.** Each grouped bar chart shows the distribution of path lengths (green) and genetic distances (blue) for all pairs of starting (non-peak) and ending (peak) genotypes during $10^5$ adaptive walks. When distribution means are indistinguishable, the black dashed vertical line represents the mean of both distributions. Otherwise, the green and blue horizontal lines represent average path lengths and mutational distance, respectively. We used a t-test of the null hypothesis that the means of these distance distributions are statistically indistinguishable. **c. Distribution of accessible path lengths and genetic distances to the Fis peak.** We observed no significant difference between the distributions of genetic distance and path length (4.8±0.7, mean ± s.d., Welch Two Sample t-test, t = 30.113, df = 196,700, P-value= 0.54, $N_1 = 10^5$ genetic distances, $N_2 = 10^5$ path lengths). **d. Distribution of accessible path**

**lengths and genetic distances to the IHF peak.** We observed a significant yet small difference between the distributions of genetic distance and path length (4.5±1 and 4.7±1.2, respectively, mean ± s.d., Welch Two Sample t-test, $t = 35.973$, $df = 196{,}700$, P-value< $2.2\times10^{-16}$, $N_1 = 10^5$ genetic distances, $N_2 = 10^5$ path lengths).

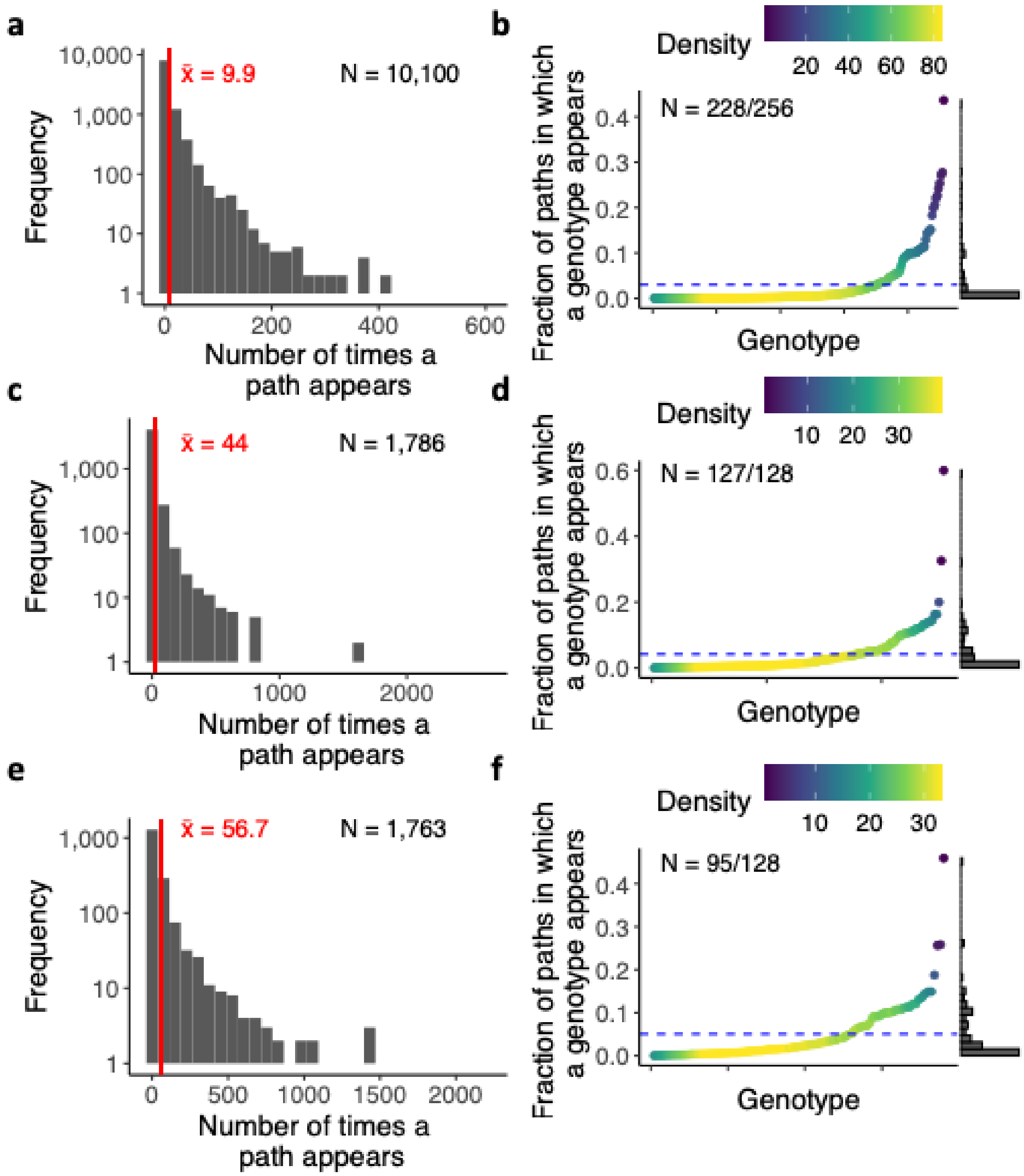


**Supplementary Figure S19. Frequencies of paths and sequences among adaptive walks.** The data is based on simulations with population sizes of $10^8$ individuals. **a-b Adaptive walks from CRP to Fis. a. Distribution of the number of unique paths.** The histogram is based on $10^5$ adaptive walks from CRP to Fis and shows the total number of times any one unique path of mutational steps occurs among these adaptive walks. The total number N of unique paths among the $10^5$ walks is represented at the top right of the histogram. The vertical red line represents the mean of the distribution, which is also shown at the top-left of the histogram ($\bar{x}$, in red). Note the logarithmic ($\log_{10}$) scale on the y-axis. **b. The distribution of genotypic visitation across multiple evolutionary paths.** This analysis considers a total of $10^5$ paths, each starting from one of 10 strongest genotypes for CRP, and explores how often each of the 256 genotypes comprising the landscape is visited within these paths. The total number N of unique genotypes that are observed across all paths is shown at the top left of the plot. The blue horizontal dashed line at y = 0.01, represents the average frequency at which genotypes appear across all paths, meaning that each genotype, on average, appears in 1% of the total number of paths. The color gradient (color legend) shows the number of genotypes along the horizontal axis. The histogram on the right side of the figure shows the overall distribution of the fraction of paths from the total number of paths in which a genotype appears. It is expected that some genotypes are overrepresented among adaptive walks, especially the ones closer to the peak. The reason is that the number of accessible paths to a peak decreases with the distance of a genotype to a peak, leaving fewer genotypes to be traversed by adaptive walks, the closer a genotype is to the peak104,105. **c-d. Like a-b, but for CRP-IHF (IHF to CRP direction). e-f. Like a-b, but for IHF-Fis adaptive walks (Fis to IHF direction).**

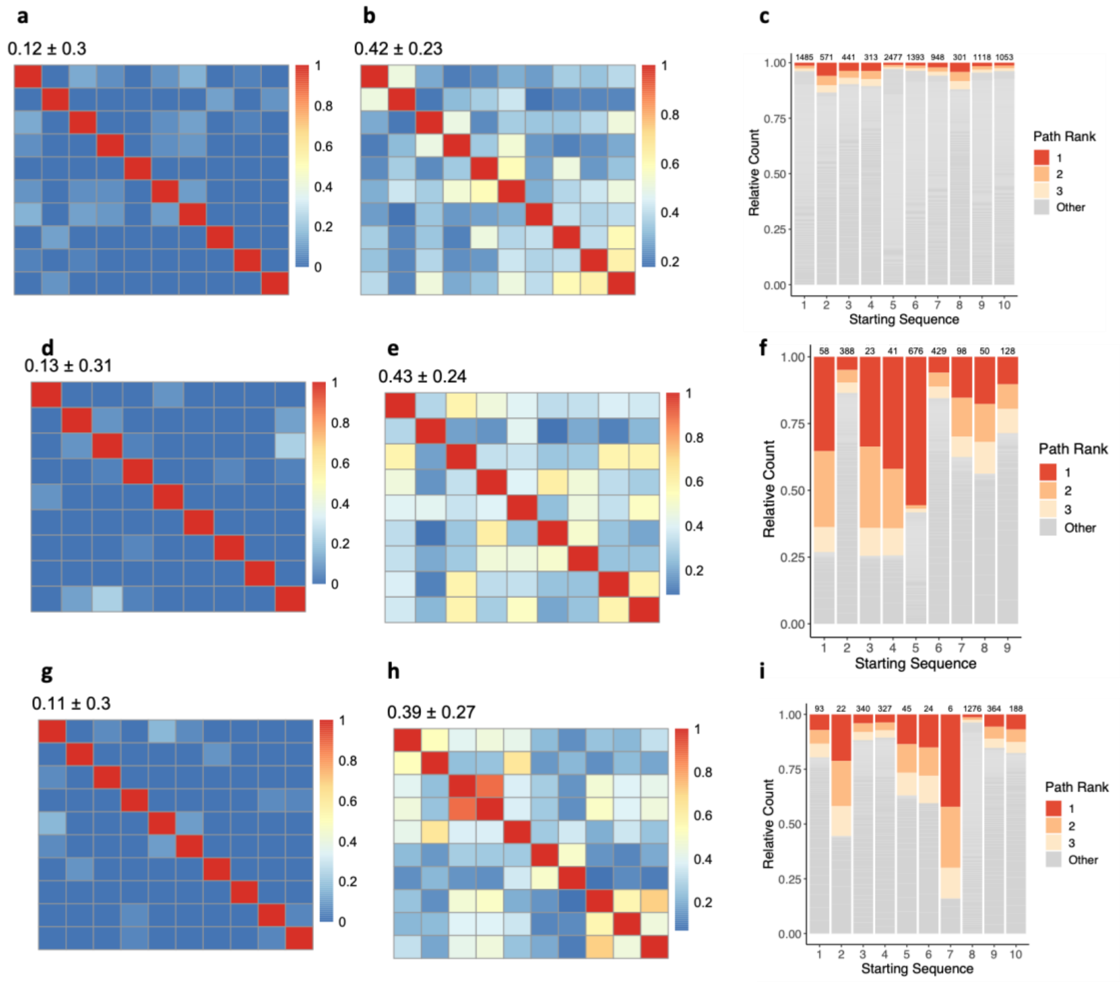


**Supplementary Figure S20. Little path overlap, moderate sequence overlap and strong path preferences during adaptive walks towards stronger binders for a specific TF.** Each plot represents data from $10^5$ Kimura adaptive walk simulations (10 starting genotypes × $10^4$ adaptive walks each) with populations of $10^8$ individuals. **a-c. CRP-Fis landscape (CRP to Fis direction).** We simulated $10^4$ adaptive walks starting from 10 genotypes with the highest regulation strength for CRP selected for stronger regulation strength towards Fis **a. Adaptive walks from distinct starting points share few paths.** For each starting genotype, we placed all unique paths among all paths starting from that genotype ($10^4$ paths) into a set. We then computed the Jaccard index[35] among all possible pairs of these sets (**Supplementary Methods 7.5**). A value of one (red) signifies complete identity between the paths from two different starting genotypes, implying the sets are identical (every path occurs in both sets). Conversely, a value of zero (blue) indicates that the sets of paths starting from different genotypes share no paths. Numbers on top of the heatmap indicate the mean ± one standard deviation of the Jaccard index for all pairs of path sets traversed during walks from different starting genotypes. **b. Paths to peaks share many genotypes.** As in (a), but the heatmap matrix quantifies the fraction of shared genotypes between paths starting from each pair of starting variants. To obtain the data for this matrix, we placed all unique genotypes that occur in all $10^4$ evolutionary paths starting from that genotype (excluding the starting genotype itself) into a set. We then computed the Jaccard index among all possible pairs of these sets (**Supplementary Methods 7.5**). A value of one (red) signifies complete identity between the set of genotypes traversed during walks starting from a given pair of genotypes. Conversely, a value of zero (blue) indicates that the paths share no genotypes. Numbers on top of the heatmap indicate the mean

± one standard deviation of the Jaccard index for all pairs of genotype sets for non-identical starting genotypes. **c. For different starting genotypes, some paths are more frequent than others.** For each of our 10 starting genotypes, we counted how often identical paths, i.e., paths sharing all genotypes along the path, occurred. We then ranked the paths according to their frequency (as a fraction of all $10^4$ paths) and plotted this frequency for each starting genotype as a stacked bar plot (y-axis), in which the frequencies of the three top-ranked paths are displayed in color (red, orange, yellow), and the cumulative frequency of all other paths is shown in grey. Atop each bar, numerical values indicate the total number of unique paths (among $10^4$ paths starting from the initial genotype). **d-f.** As in a-c, but for the CRP-IHF landscape, and for paths towards strong IHF regulation strength**. g-i.** As in a-c, but for the Fis-IHF landscape, and for paths towards strong IHF regulation strength**.**

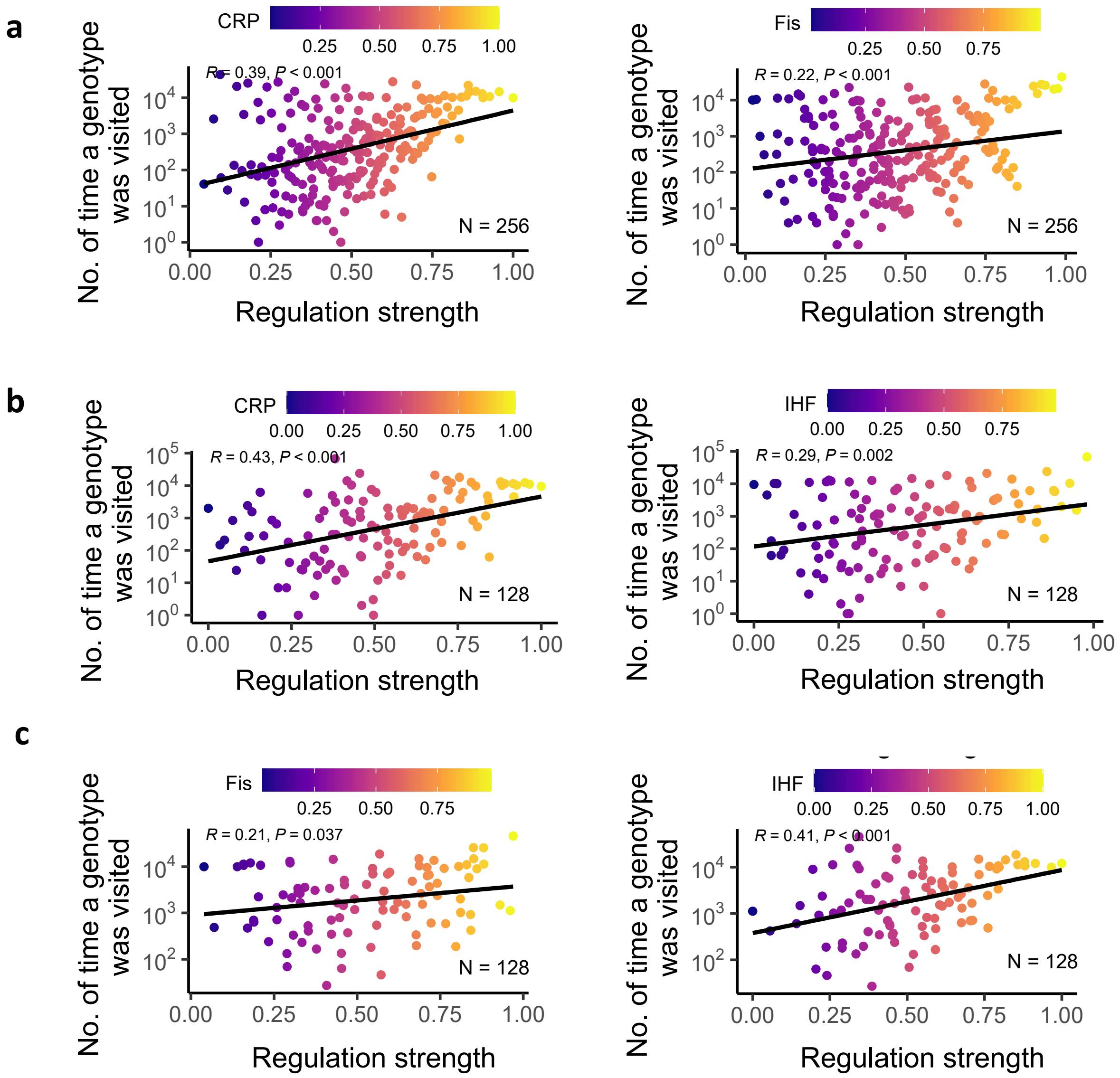


**Supplementary Figure S21. Adaptive walks tend to pass through strong TFBSs more often.** Data in each plot is based on $10^5$ Kimura adaptive walk simulations (10 starting genotypes with the highest regulatory strengths for the opposite TF x $10^4$ adaptive walks each) with population sizes of $10^8$. For each such TFBS (circles) the plot shows both the regulation strength (x-axis) of a TFBS and the number of times the TFBSs occurred in the $10^5$ adaptive walks (y-axis, note logarithmic scale). Circle colors indicate regulation strength (see color legend). Black line: linear regression. R: Pearson correlation coefficient. P-values are based on a t-statistic test of the null hypothesis that there is no association between the number of times genotypes were attained and their regulation strengths. Panels on the left refer to $10^5$ Kimura adaptive walks towards TF1 and panels on the right refer to $10^5$ Kimura adaptive walks towards TF2. **a. CRP-Fis landscape.** Left panel: $10^5$ adaptive walks ending at the Fis peak. Right panel: $10^5$ walks ending at the CRP peak. **b. CRP-IHF landscape**. Left panel: like (a) but with adaptive walks ending at the IHF peak. Right panel: like (a) but with adaptive walks ending at the CRP peak. **c. Fis-IHF landscape.** Left panel: like (a) but with adaptive walks ending at the IHF peak. Right panel: like (a) but with adaptive walks ending at the Fis peak.

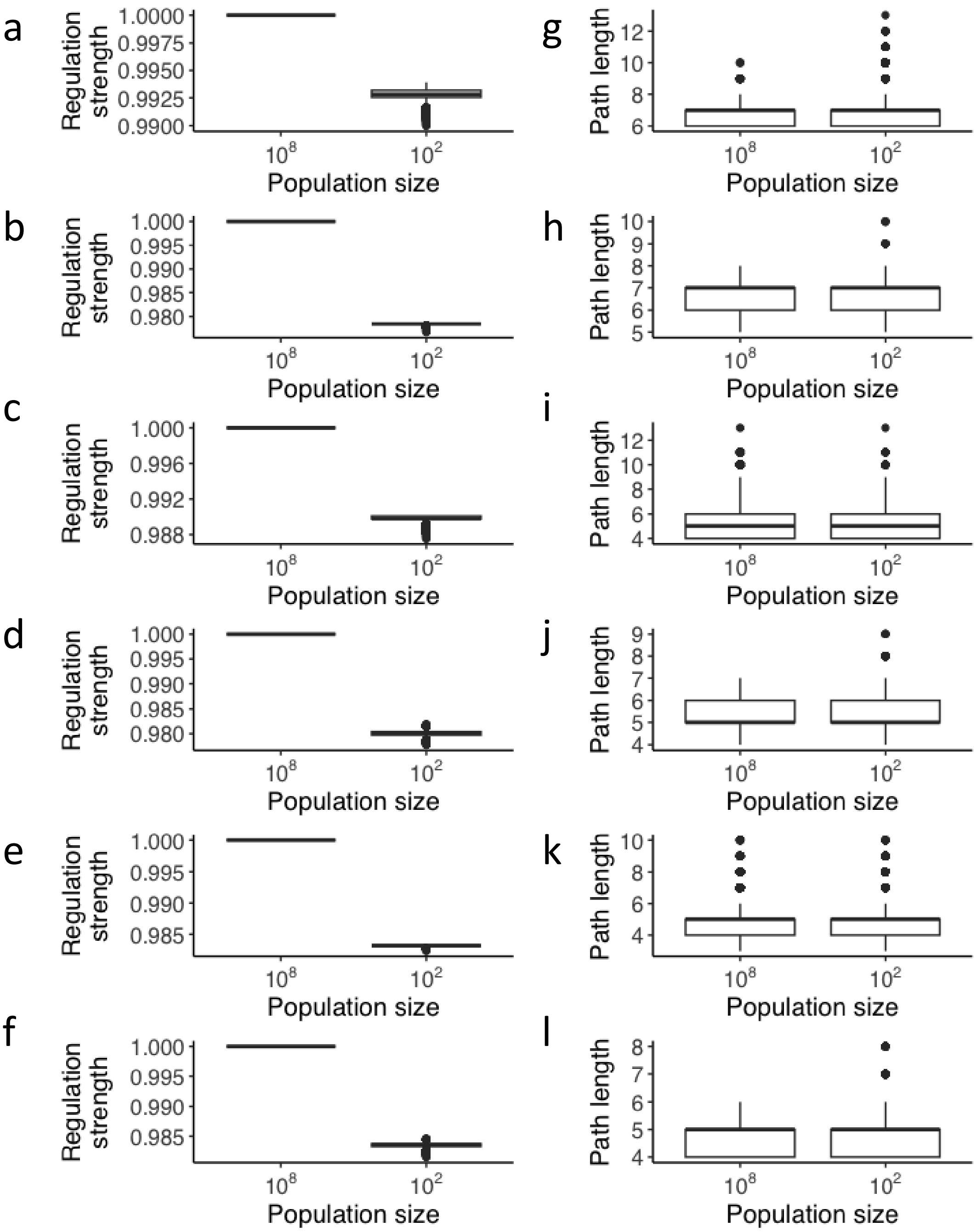


**Supplementary Figure S22. Drift has negligible effects on adaptive walks. a-f. Differences in the fitness (regulation strengths) of the end points of Kimura walks between large ($10^8$) and small ($10^2$) population sizes.** We computed average regulation strengths for small population sizes as the average regulation strengths of the 100 steps after a peak was reached for the first time in each walk (each boxplot contains data from $10^5$ adaptive walks, with 10 starting genotypes and $10^4$ walks for each starting genotype). Although statistically significant differences between the means in small and large populations exist, these are very small (less than 2% of regulation strength). **a. CRP-Fis exaptation landscape with selection in the direction of CRP strong binders.** Mean for large pop.:1.00, mean ± s.d. for small pop.:0.99 ± 0.0005; Welch one Sample t-test, t = -4828.2, df = 99999, *P-value*< $2.2 \times 10^{-16}$, $N_1 = 10^5$ regulation strengths, $N_2 = 1$. **b. Same landscape as (a) but in the other direction.** Mean for large pop.:1.00, mean ± s.d. for small pop.:0.98 ± 0.0001; Welch one Sample t-test, t = -95137, df = 99999, *P-value*< 2.2 x $10^{-16}$, $N_1 = 10^5$ regulation strengths, $N_2 = 1$. **c. CRP-IHF exaptation landscape with selection in the direction of CRP.** Mean for large pop.:1.00, mean ± s.d., for small pop.:0.98 ± 0.0003; Welch one Sample t-test, t = -13691, df = 99999, *P-value*< 2.2 x $10^{-16}$, $N_1 = 10^5$ regulation strengths, $N_2 = 1$. **d. Same landscape as (c) but in the opposite direction.** Mean for large pop.:1.00, mean ± s.d. for small pop.:0.99 ± 0.0005; Welch one Sample t-test, t = -10896, df = 99999, P-value< 2.2 x $10^{-16}$, $N_1 = 10^5$ regulation strengths, $N_2 = 1$. **e. Fis-IHF exaptation landscape with selection in the direction of Fis.** Mean for large pop.:1.00, mean ± s.d. for small pop.:0.98 ± 0.0003; Welch one Sample t-test, t = -17228, df = 99999, P-value< 2.2 x $10^{-16}$, $N_1 = 10^5$ regulation strengths, $N_2 = 1$. **f. Same landscape as (e) but in the opposite direction.** Mean for large pop.:1.00, mean ± s.d. for small pop.:0.98 ± 0.00003; Welch one Sample t-test, t = -160875, df = 99999, P-value< 2.2 x $10^{-16}$, $N_1 = 10^5$ regulation strengths, $N_2 = 1$. **g-l. Distribution of path lengths (evolutionary paths) ending in a peak for both large and small population sizes.** Each boxplot contains data from $10^5$ adaptive walks, i.e., 10 starting genotypes and $10^4$ walks from each starting genotype. We computed the length of a path attaining a peak for small populations as the number of steps until the peak was reached for the first time. (paths in small populations are only 10% longer than in large populations). **g. Path lengths for the CRP-Fis exaptation landscape with selection in the direction of CRP strong binders.** Mean ± s.d. for large pop.:6.80 ± 0.60, mean ± s.d. for **small** pop.:6.87 ± 0.71; Welch Two Sample t-test, t = 24.929, df = 194724, P-value< 2.2 x $10^{-16}$, $N_1 = 10^5$ path lengths, $N_2 = 10^5$ path lengths. **h. Same landscape as (g) but in the other direction.** Mean ± s.d. for large pop.:6.60 ± 0.80, mean ± s.d. for **small** pop.:6.60 ± 0.80; Welch Two Sample t-test, t = 0.26254, df = 2e+05, P-value= 0.7929, $N_1 = 10^5$ path lengths, $N_2 = 10^5$ path lengths. **i. Path lengths for the CRP-IHF exaptation landscape with selection in the direction of CRP.** Mean ± s.d. for large pop.:5.30 ± 0.90, mean ± s.d. for **small** pop.:5.30 ± 0.90; Welch Two Sample t-test, t = 0.31278, df = 2e+05, P-value= 0.7544, $N_1 = 10^5$ path lengths, $N_2 = 10^5$ path lengths. **j. Same landscape as (i) but in the other direction.** Mean ± s.d. for large pop.:5.00 ± 0.91, mean ± s.d. for **small** pop.:5.28 ± 1.06; Welch Two Sample t-test, t = 58.633, df = 182278, P-value< 2.2 x $10^{-16}$, $N_1 = 10^5$ path lengths, $N_2 = 10^5$ path lengths. **k. Path lengths for the Fis-IHF exaptation landscape with selection in the direction of Fis.** Mean ± s.d. for large pop.:4.80 ± 0.75, mean ± s.d. for **small** pop.:4.80 ± 0.75; Welch Two Sample t-test, t = 0.89995, df = 199992, P-value= 0.3681, $N_1 = 10^5$ path lengths, $N_2 = 10^5$ path lengths. **l. Same landscape as (k) but in the other direction.** Mean ± s.d. for large pop.:4.66 ± 1.17, mean ± s.d. for **small** pop.:4.66 ± 1.17; Welch Two Sample t-test, t = 0.61863, df = 199989, P-value= 0.5362, $N_1 = 10^5$ path lengths, $N_2 = 10^5$ path lengths.

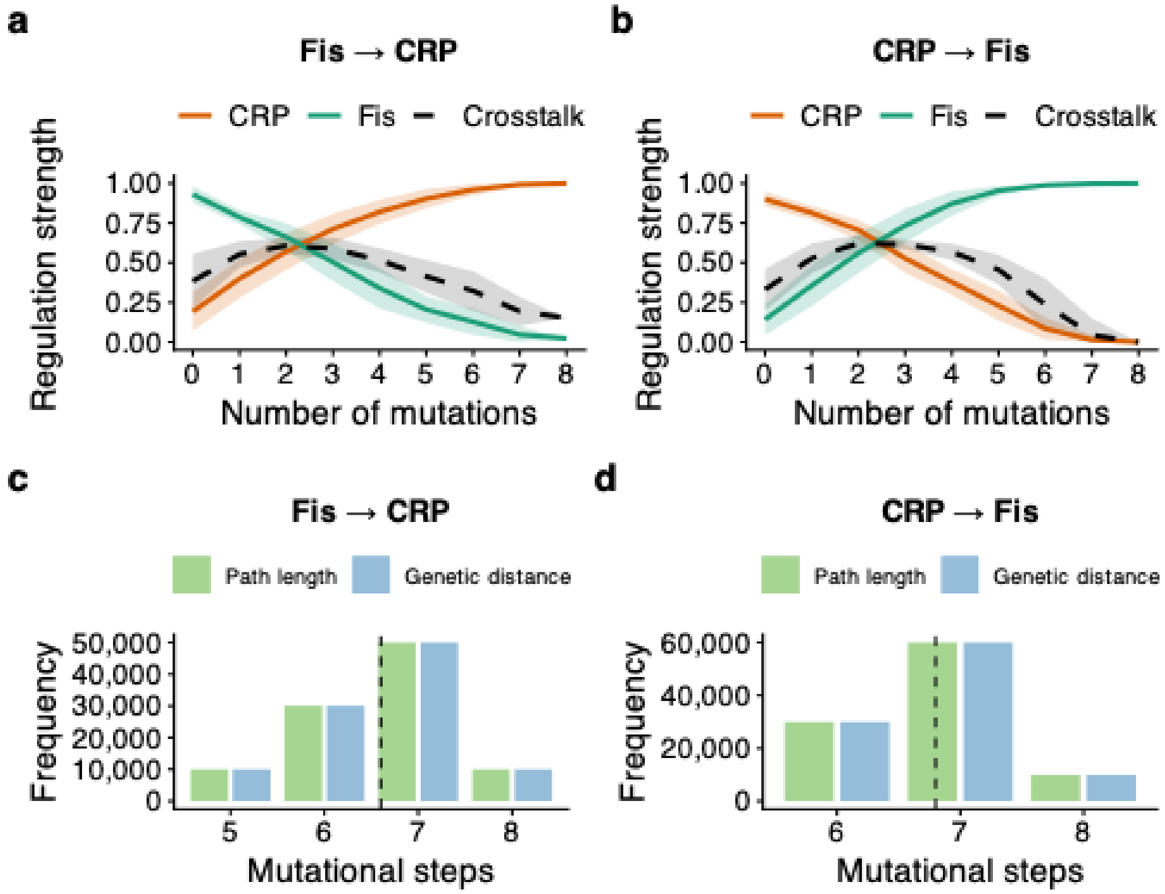


**Supplementary Figure S23. Rapid exaptation on the CRP-Fis landscape outside of the SSWM regime.** Data for each plot is based on $10^5$ simulated adaptive greedy walks, i.e., $10^4$ adaptive walks for each of the 10 starting genotypes with the highest regulation strength for the opposite TF. **a-b. All adaptive walks readily attain peaks.** Each plot shows how regulation strength (vertical axis) for CRP (orange) and Fis (green) changes as a function of path length traversed (horizontal axis) from the initial sequence to the reached peak during adaptive walks. Crosstalk along the walks is shown by the black dashed line and is defined as the geometric mean of TF1 and TF2 regulation strengths. Curves represent averages over all $10^5$ adaptive walks, and shading indicates one standard deviation of regulation strengths. **a. All adaptive walks attain the CRP peak.** The data is based on $10^4$ walks starting from each of the 10 strongest binding sites for Fis, favoring increasing CRP binding. **b. All adaptive walks attain the Fis peak.** Like panel (a), but for adaptive walks starting from the 10 strongest binding sites for CRP, favoring increasing Fis binding. **c-d. Evolutionary paths to a peak are not much longer than minimal genetic distances.** Each grouped bar chart shows the distribution of path lengths (green) and genetic distances (blue) for all pairs of starting (non-peak) and ending (peak) genotypes during $10^5$ adaptive walks. The black dashed vertical line represents the mean of both distributions, which is not significantly different between distributions. We used a t-test of the null hypothesis that the means of these distance distributions are statistically indistinguishable. **c. Distribution of accessible path lengths and genetic distances to the CRP peak.** No significant difference was observed between the distributions of genetic distance and path length (mean ± s.d.: 6.6 ± 0.8 for both genetic distance and path length; Welch Two Sample t-test: t = 31.973, df = 196,700, P-value= 0.7, $N_1 = 10^5$ genetic distances, $N_2 = 10^5$ path lengths). **d. Distribution of accessible path lengths and**

**genetic distances to the Fis peak.** No significant difference was observed between the distributions of genetic distance and path length (mean ± s.d.: 6.8 ± 0.6; Welch Two Sample t-test: $t = 35.113$, $df = 196{,}700$, P-value= 0.82, $N_1 = 10^5$ genetic distances, $N_2 = 10^5$ path lengths).

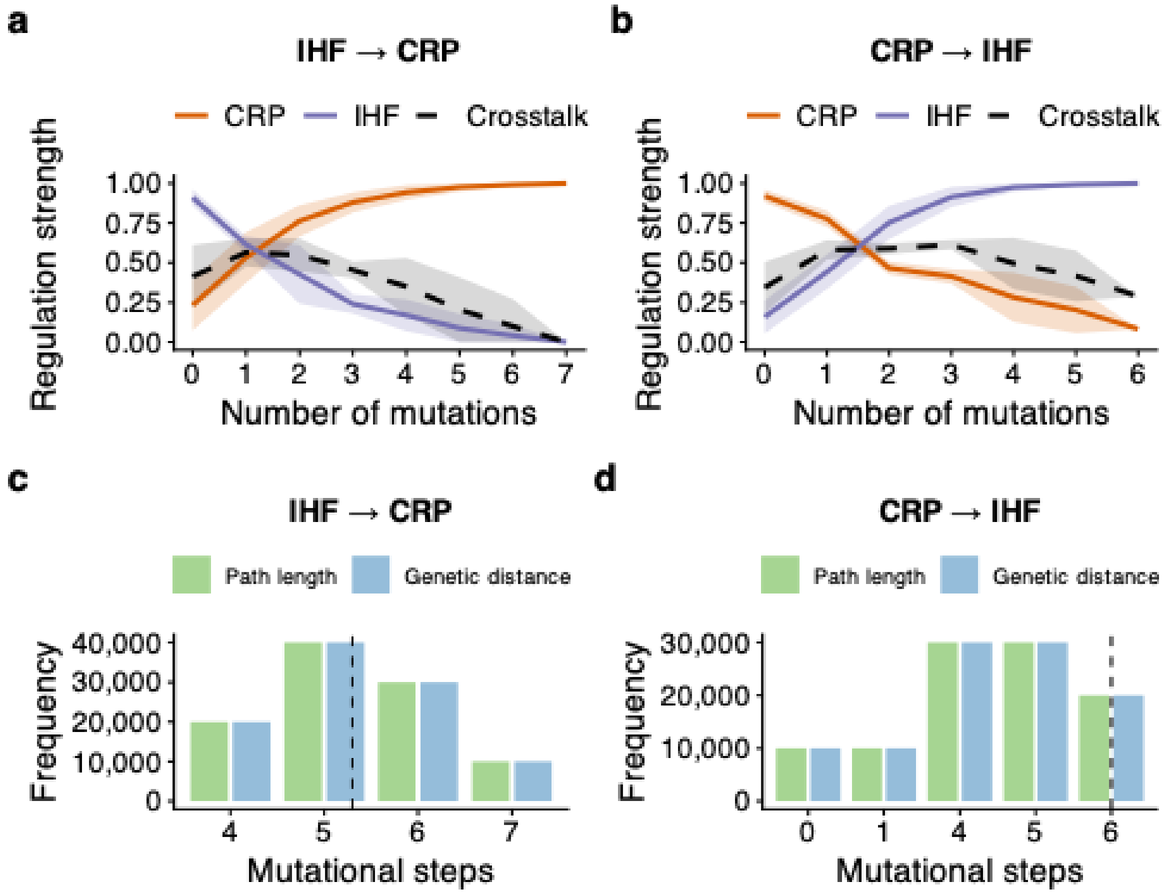


**Supplementary Figure S24. Rapid exaptation on the CRP-IHF landscape outside of the SSWM regime.** Data for each plot is based on $10^5$ simulated adaptive greedy walks, i.e., $10^4$ adaptive walks for each of the 10 starting genotypes with the highest regulation strength for the opposite TF. **a-b. All adaptive walks readily attain peaks.** Each plot shows how regulation strength (vertical axis) for CRP (orange) and IHF (purple) changes as a function of path length traversed (horizontal axis) from the initial sequence to the reached peak during adaptive walks. Crosstalk along the walks is shown by the black dashed line and is defined as the geometric mean of CRP and IHF regulation strengths. Curves represent averages over all $10^5$ adaptive walks, and shading indicates one standard deviation of regulation strengths. **a. All adaptive walks attain the CRP peak.** The data is based on $10^4$ walks starting from each of the 10 strongest binding sites for IHF, favoring increasing CRP binding. **b. All adaptive walks attain the IHF peak.** Like panel (a), but for adaptive walks starting from the 10 strongest binding sites for CRP, favoring increasing IHF binding. **c-d. Evolutionary paths to a peak are not much longer than minimal genetic distances.** Each grouped bar chart shows the distribution of path lengths (green) and genetic distances (blue) for all pairs of starting (non-peak) and ending (peak) genotypes during $10^5$ adaptive walks. The black dashed vertical line represents the mean of both distributions, which is not significantly different between distributions. We used a t-test of the null hypothesis that the means of these distance distributions are statistically indistinguishable. **c. Distribution of accessible path lengths and genetic distances to the CRP peak.** We observed no significant difference between the means of genetic distance and path length (5.3±0.9, mean ± s.d., Welch Two Sample t-test, $t = 31.973$, $df = 196{,}700$, P-value= 0.62, $N_1 = 10^5$ genetic distances, $N_2 = 10^5$ path lengths). **d. Distribution of accessible path lengths and genetic distances to the IHF peak.** We observed no significant difference between mean genetic distance and path length (6±0.8, mean ± s.d., Welch Two

Sample t-test, t = 35.113, df = 196,700, P-value= 0.77, $N_1 = 10^5$ genetic distances, $N_2 = 10^5$ path lengths).

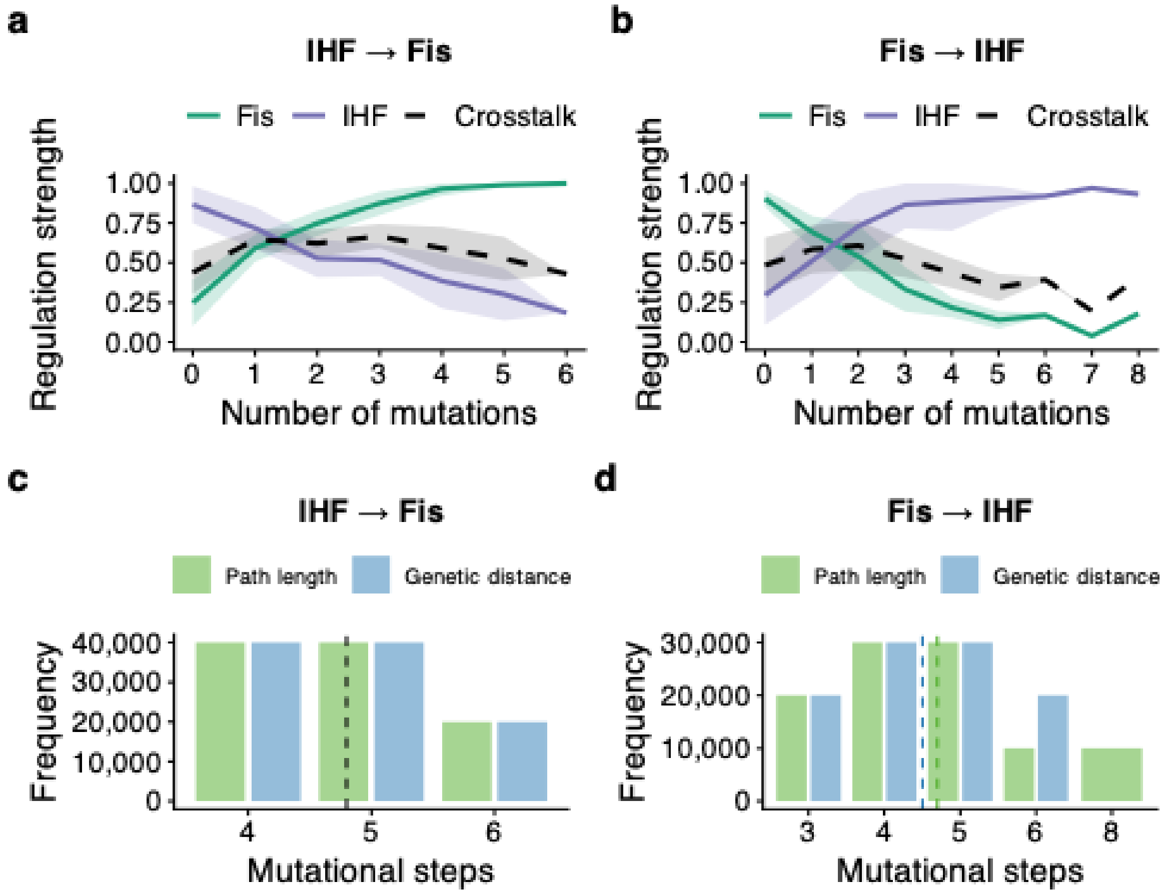


**Supplementary Figure S25. Rapid exaptation on the Fis-IHF landscape outside of the SSWM regime.** Data for each plot is based on $10^5$ simulated adaptive greedy walks, i.e., $10^4$ adaptive walks for each of the 10 starting genotypes with the highest regulation strength for the opposite TF. **a-b. All adaptive walks readily attain peaks.** Each plot shows how regulation strength (vertical axis) for Fis (green) and IHF (purple) changes as a function of path length traversed (horizontal axis) from the initial sequence to the reached peak during adaptive walks. Crosstalk along the walks is shown by the black dashed line and is defined as the geometric mean of Fis and IHF regulation strengths. Curves represent averages over all $10^5$ adaptive walks, and shading indicates one standard deviation of regulation strengths. **a. All adaptive walks attain the Fis peak.** The data is based on $10^4$ walks starting from each of the 10 strongest binding sites for IHF, favoring increasing Fis binding. **b. All adaptive walks attain the IHF peak.** Like panel (a), but for adaptive walks starting from the 10 strongest binding sites for Fis, favoring increasing IHF binding. **c-d. Evolutionary paths to a peak are not much longer than minimal genetic distances.** Each grouped bar chart shows the distribution of path lengths (green) and genetic distances (blue) for all pairs of starting (non-peak) and ending (peak) genotypes during $10^5$ adaptive walks. When distribution means are indistinguishable, the black dashed vertical line represents the mean of both distributions. Otherwise, the green and blue horizontal lines represent average path lengths and mutational distance, respectively. We used a t-test of the null hypothesis that the means of these distance distributions are statistically indistinguishable. **c. Distribution of accessible path lengths and genetic distances to the Fis peak.** We observed no significant difference between the distributions of genetic distance and path length (4.8±0.7, mean ± s.d., Welch Two Sample t-test, $t = 35.973$, $df = 196{,}700$, P-value= 0.39, $N_1 = 10^5$ genetic distances, $N_2 = 10^5$ path lengths).

**d. Distribution of accessible path lengths and genetic distances to the IHF peak.** We observed a significant yet small difference between the distributions of genetic distance and path length (4.5±1 and 4.7±1.4, respectively, mean ± s.d., Welch Two Sample t-test, $t = 30.113$, $df = 196{,}700$, P-value< $2.2\times10^{-16}$, $N_1 = 10^5$ genetic distances, $N_2 = 10^5$ path lengths).

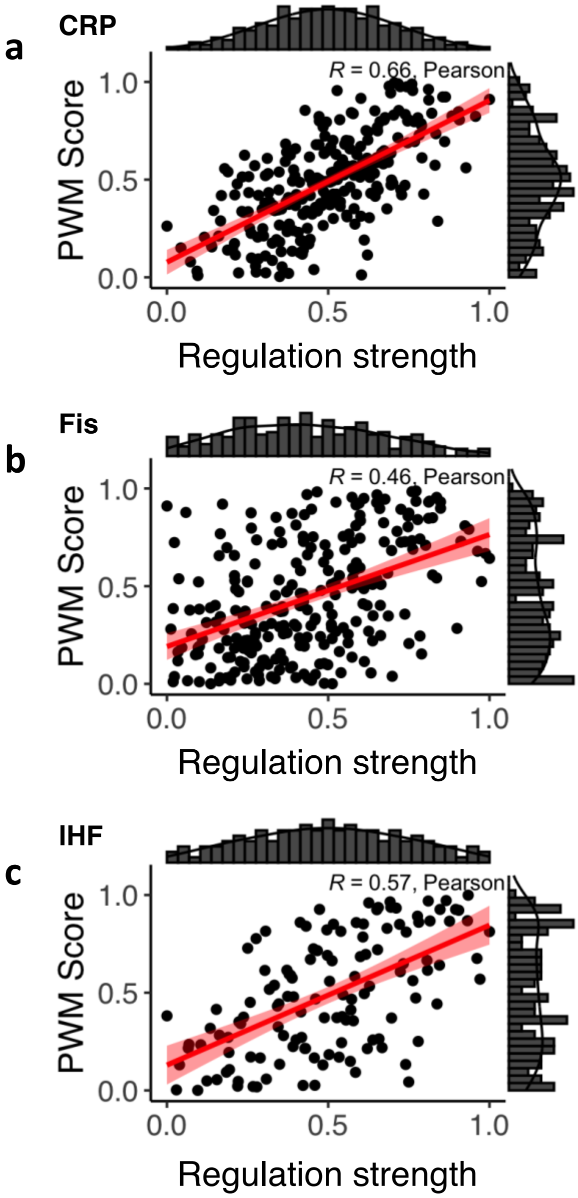


**Supplementary Figure S26. PWM scores are associated with measured regulation strengths.** Data are based on 300 experimentally validated TFBS sequences from our sort-seq experiments (N = 100 per TF: CRP, Fis, and IHF). For each TFBS, we compared measured regulation strength (x-axis) with the predicted PWM score (y-axis). Red lines indicate the linear regression fit; shaded regions represent the 95% confidence interval of the regression. R denotes the Pearson correlation coefficient. We assessed statistical significance using a two-sided Pearson correlation test (df = 98). Histograms on the top and right show the marginal distributions of regulation strengths and PWM scores, respectively. **a. CRP TFBSs**. R = 0.66, $P = 1.1 \times 10^{-13}$ **b. Fis TFBSs**. R = 0.46, $P = 1.9 \times 10^{-6}$ **c. IHF TFBSs:** R = 0.57, $P = 1.3 \times 10^{-9}$.

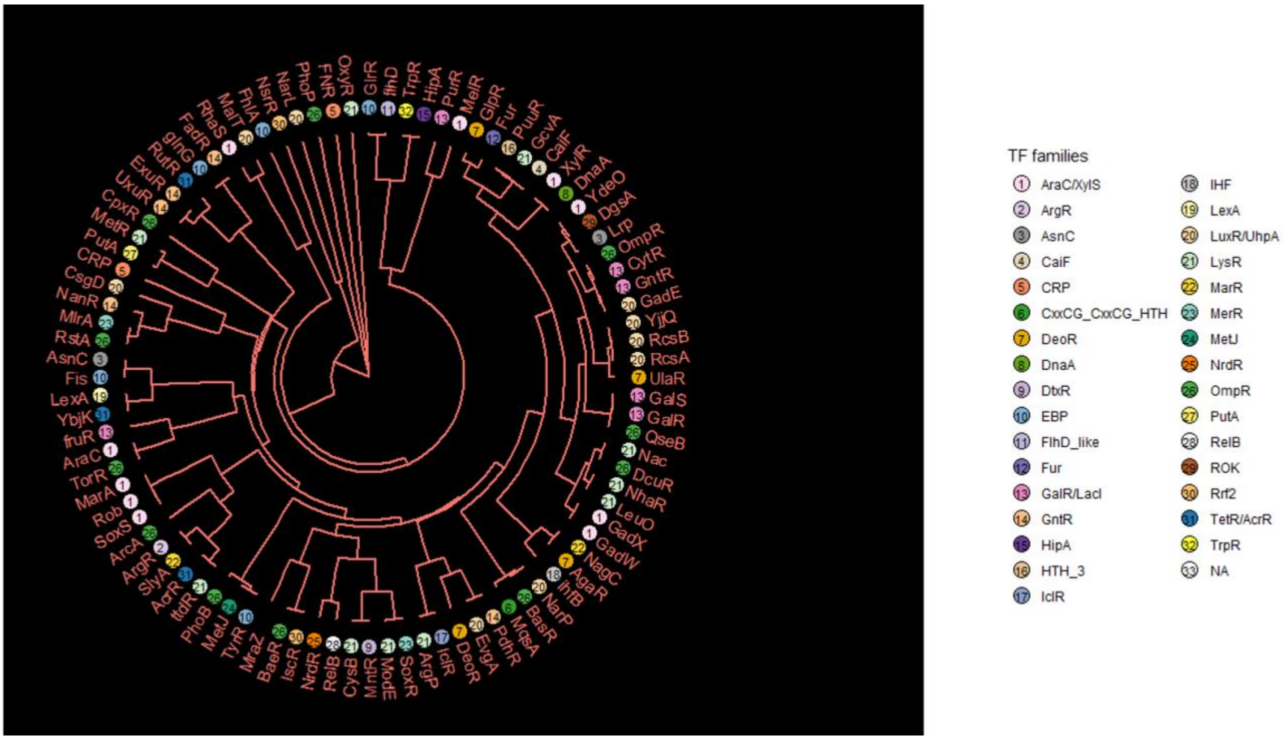


**Supplementary Figure S27. Clustering of *E. coli* PWMs for exploring potential crosstalk candidates.** The circular dendrogram represents the clustering of 109 available PWMs from RegulonDB278. Annotations for 33 TF families are represented as colored/numbered circles. Briefly, we retrieved PWMs from RegulonDB[38] and aligned them through a combination of local Smith-Waterman alignment [10], and position weighting based on positional information. We then used these alignments to compute pairwise distances for all pairs of PWMs, clustered the resulting distance matrix hierarchically, and plotted the dendrogram produced by the clustering algorithm using the *ggtree*[79] package from R (**see Supplementary Methods 7.9**). This *in silico* analysis shows that TFs with similar PWMs have greater potential for crosstalk than transcription factors with less similar PWMs.

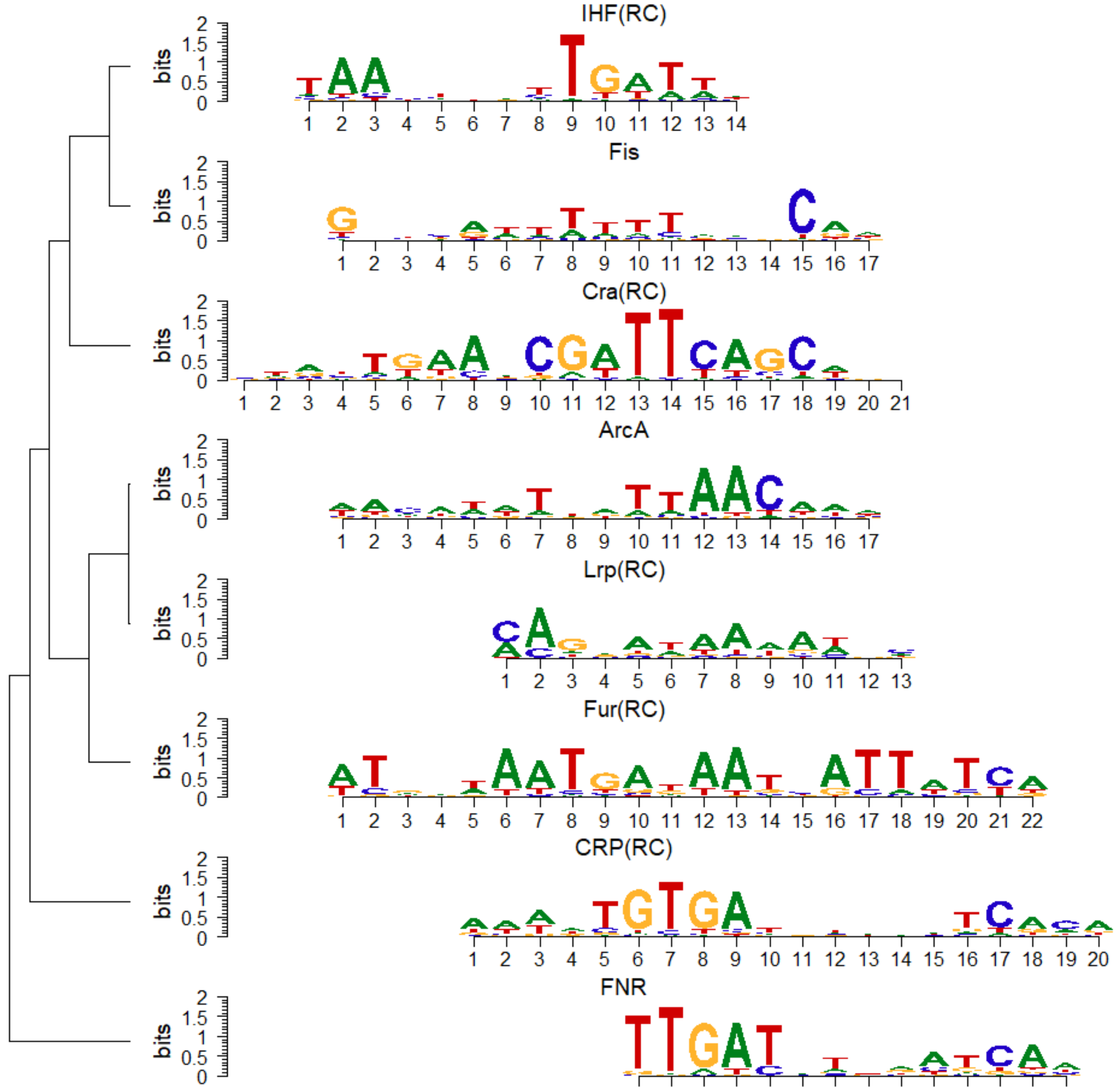


**Supplementary Figure S28. Clustering of PWMs for *E. coli's* global regulators.** The figure displays the hierarchical clustering of PWMs for eight global regulators in *E. coli*. Each regulator's PWM is represented by a sequence logo, showing the nucleotide positions along the x-axis, and the bits score along the y-axis. The nucleotide colors are as follows: Adenine (A) is green, Thymine (T) is red, Cytosine (C) is blue, and Guanine (G) is orange. The height of each letter in the sequence logo indicates the relative frequency of that nucleotide at each position, with higher letters representing higher conservation and significance. The bits score quantifies the information content at each position in the PWM, with higher scores indicating more conserved positions that contribute to the binding specificity of the transcription factor. The dendrogram on the left side of the figure shows the hierarchical clustering of the PWMs which groups transcription factors with similar PWMs together (**see Supplementary Methods 7.9**). This clustering suggests potential crosstalk due to overlapping binding site characteristics. The transcription factors included are IHF, Fis, Cra, ArcA, Lrp, Fur, CRP, and FNR. This analysis utilizes local Smith-Waterman alignment and positional weighting based on positional information to compute pairwise distances between PWMs. We clustered the resulting distance matrix hierarchically and visualized it using the ggtree package in R[79]. The data indicates that transcription factors with similar PWMs have a greater potential for regulatory crosstalk.

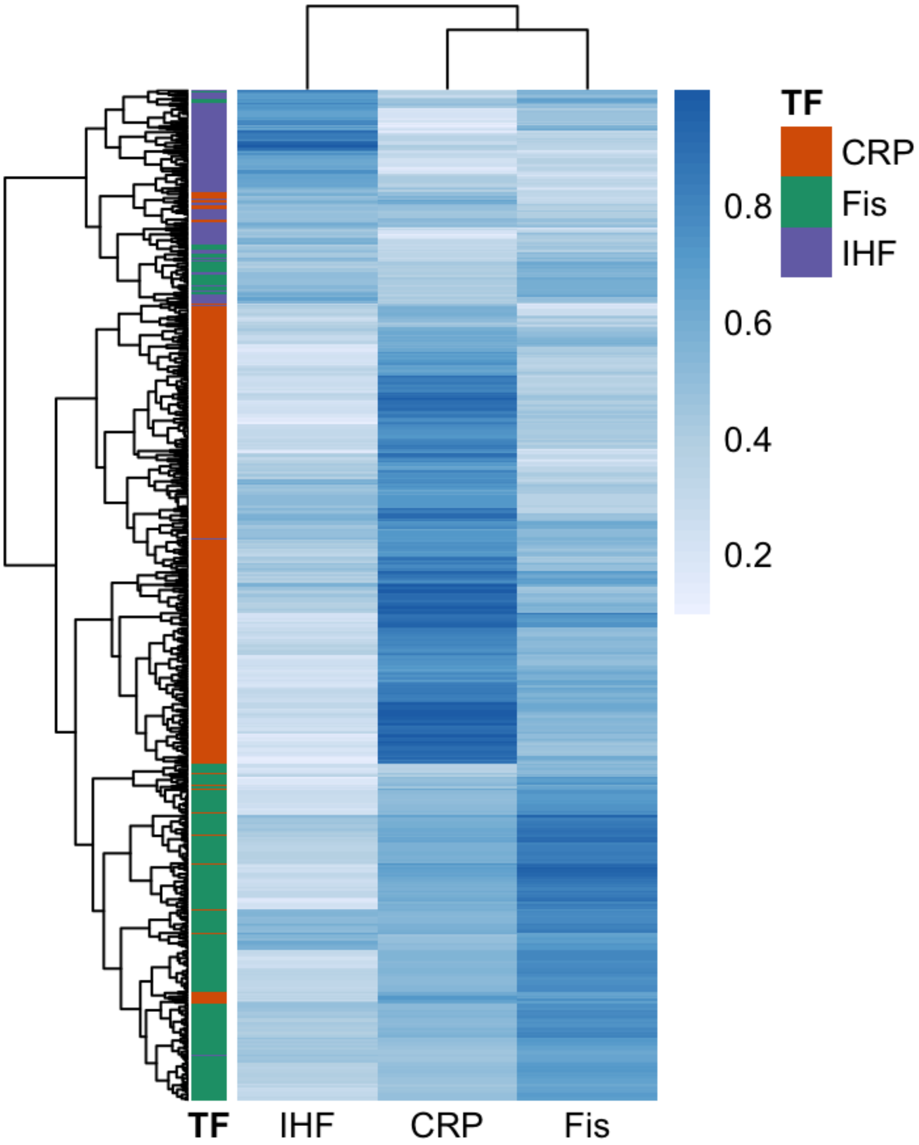


**Supplementary Figure S29. Hierarchical clustering of predicted crosstalk between CRP, Fis, and IHF for experimentally validated TFBSs for the three TFs.** The heatmap shows normalized position weight matrix (PWM) scores (0–1) for 756 experimentally validated TFBSs from RegulonDB. Rows correspond to individual TFBS sequences, ordered by their annotated cognate transcription factor: CRP (orange, N = 370), Fis (green, N = 267), and IHF (purple, N = 119). Columns show the PWM score of each sequence for the CRP, Fis, and IHF PWMs, where white indicates low predicted binding and dark blue indicates high predicted binding. If TFBSs were strictly specific for one TF, sequences would score highly only against their cognate TF. Instead, CRP- and Fis-annotated TFBSs frequently exhibit high PWM scores for the non-cognate TF: 45% of CRP sites and 61% of Fis sites exceed a cross-TF PWM score of 0.5, indicating widespread predicted cross-reactivity between these regulators. In contrast, IHF-annotated TFBSs show substantially lower crosstalk with CRP and Fis.

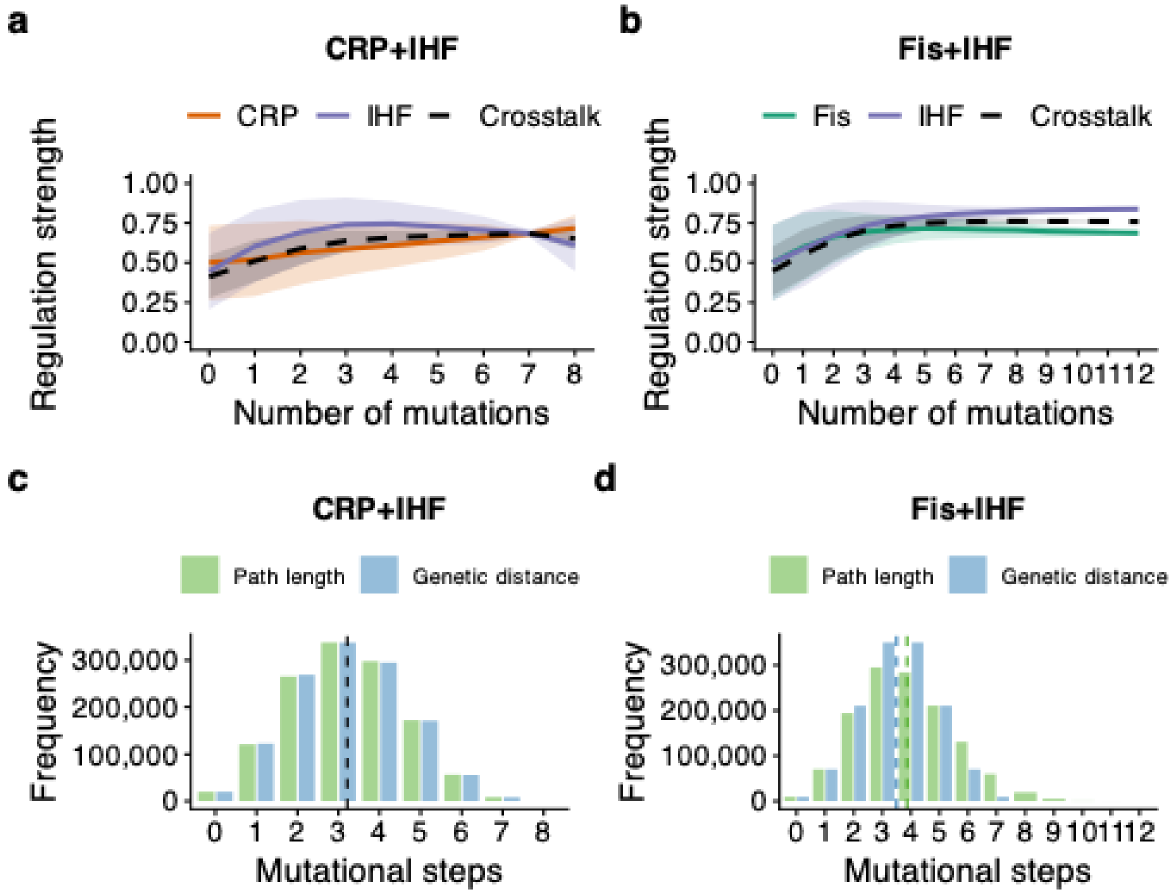


**Supplementary Figure S30. Rapid evolution of crosstalk in exaptation landscapes.** All data presented are based on Kimura adaptive walks with populations of $10^8$ individuals. Data for each plot is based on $10^4$ adaptive walks starting from each of the 128 TFBS variants in the CRP-IHF and Fis-IHF landscapes. In these simulations, selection favored dual regulation, such that we defined fitness as the sum of regulation strengths for both TFs. For each landscape, a crosstalk peak is a peak genotype that maximizes simultaneous regulation by both TFs. **a-b. All adaptive walks rapidly attain a crosstalk peak.** Each plot shows how regulation strength (vertical axis) for CRP (orange), Fis (green) and IHF (purple) changes as a function of path length traversed (horizontal axis) from the initial sequence to the reached peak during adaptive walks. Crosstalk along the walks is shown by the black dashed line and is defined as the geometric mean of TF1 and TF2 regulation strengths. Curves represent averages over all 128 $\times$ $10^4$ adaptive walks, and shading indicates one standard deviation of regulation strengths. **a. All adaptive walks attain the CRP-IHF crosstalk peak.** The panel shows the regulation strength (vertical axis) for CRP (orange) and IHF (purple) plotted against the genetic distance from the initial sequence to the attained peak (horizontal axis). **b. All adaptive walks attain the Fis-IHF crosstalk peak.** Same as panel (a) but for the Fis-IHF peak. **c-d. Evolutionary paths to intermediate peaks are not much longer than minimal genetic distances.** Each grouped bar chart shows the distribution of path lengths (green) and genetic distances (blue) for all pairs of starting (non-peak) and ending (peak) genotypes during $10^5$ adaptive walks. When distribution means are indistinguishable, the black dashed vertical line represents the mean of both distributions. Otherwise, the green and blue horizontal lines represent average path lengths and mutational distance, respectively. We used a t-test of the null hypothesis that the means of these distance distributions are statistically indistinguishable. **c. Distribution of**

**accessible path lengths and genetic distances to the CRP-IHF peak.** The distributions are not statistically distinguishable (mean ± s.d.: 3 ± 1.3, Welch Two Sample t-test: t = -5.5782, df = 2,029,535, P-value= 0.7, N = 128 × $10^4$). **d. Distribution of accessible path lengths and genetic distances to the Fis-IHF peak.** The difference between the means of genetic distance and path length is small, yet statistically distinguishable (mean ± s.d.: 3.5 ± 1.5 and 3.9 ± 1.6, respectively, Welch Two Sample t-test: t = 197.48, df = 2,406,898, P-value< 2.2 × $10^{-16}$, N = 128 × $10^4$).

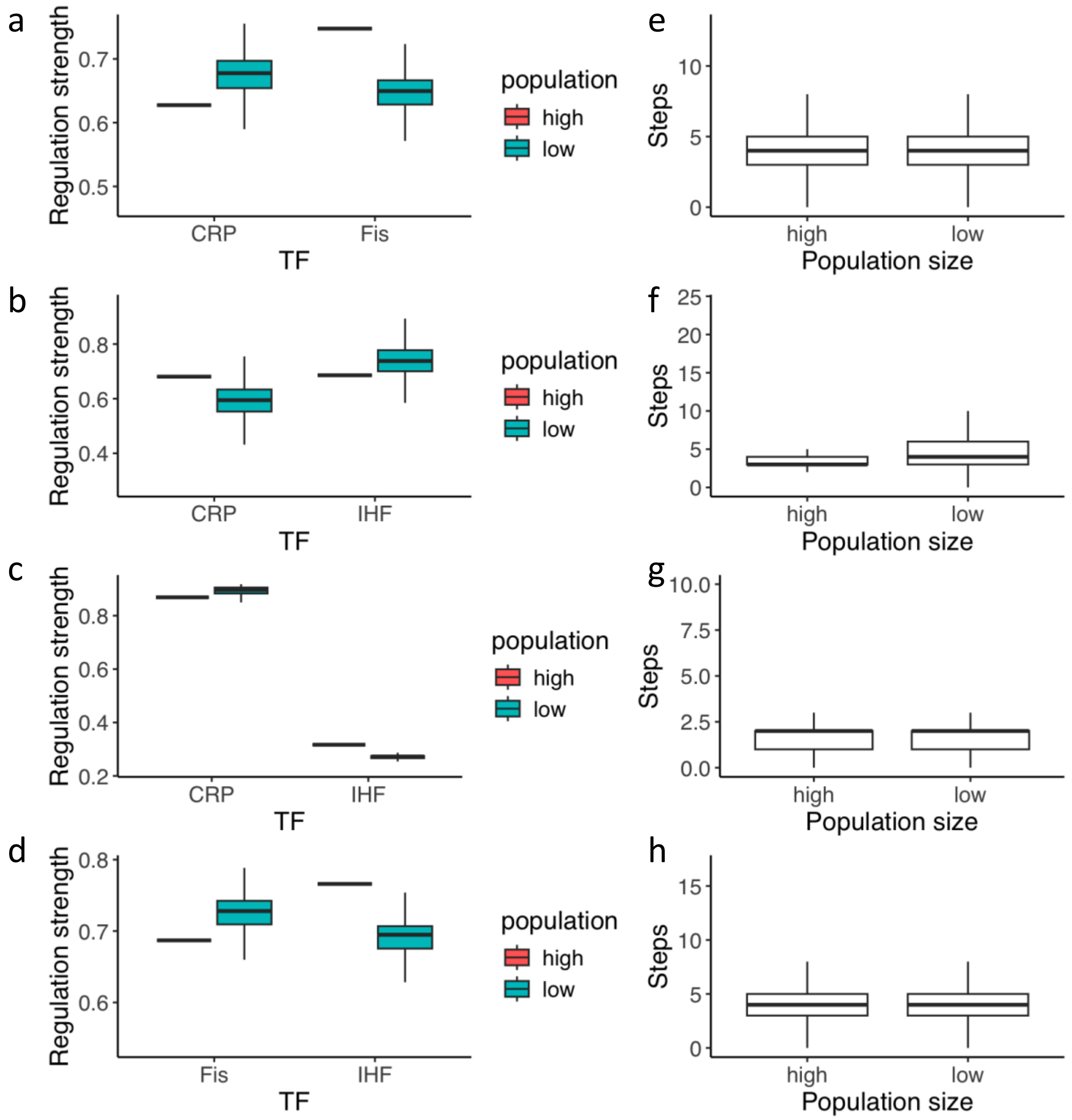


**Supplementary Figure S31. Drift has small effects on adaptive walks favouring crosstalk. a-d. Differences in the regulation strengths of both TFs at the endpoints of Kimura walks for large ($10^8$) and small ($10^2$) populations.** We analyzed the **r**egulation strengths for both TFs for each exaptive landscape (x-axis). We computed average regulation strengths for small populations (blue boxplots) as the average regulation strengths for 100 mutation-fixation steps after a peak had been reached for the first time in each walk ($10^5$ walks per boxplot, with 10 starting genotypes and $10^4$ walks from each). Although statistically significant differences between the means for small and large populations exist, they are very small (<10% in regulation strength). **a. CRP-Fis exaptation landscape.** Mean for large pop. 0.63, mean ± s.d. for small pop. 0.67 ± 0.046; Welch one Sample t-test, t = -2318.5, df = 2559999, P-value< 2.2 x $10^{-16}$, $N_1$ = 256x$10^4$ regulation strengths, $N_2$ = 1. **b CRP-IHF exaptation landscape with adaptive walks ending at the most accessed intermediate peak.** Mean for large pop. 0.68, mean ± s.d. for small pop. 0.59 ± 0.089; Welch one Sample t-test, t = 1600, df = 1126612, P-value< 2.2 x $10^{-16}$, $N_1$ = 128x$10^4$ regulation strengths, $N_2$ = 1. **c CRP-IHF exaptation landscape with adaptive walks ending at the second most accessed intermediate peak.** Mean for large pop. 0.32, mean ± s.d. for small pop. 0.28 ± 0.038; Welch one Sample t-test, t

= 318.81, df = 159099, P-value< 2.2 x $10^{-16}$, $N_1$ = 128x$10^4$ regulation strengths, $N_2$ = 1. **d Fis-IHF exaptation landscape.** Mean for large pop. 0.69, mean ± s.d. for small pop. 0.72 ± 0.037; Welch one Sample t-test, t = -1666.8, df = 1279999, P-value< 2.2 x 10-$^{16}$, $N_1$ = 128x$10^4$ regulation strengths, $N_2$ = 1. **e-h. Distribution of path lengths (evolutionary paths) ending in a peak for both large and small populations.** The length of a path attaining a peak for small population sizes was computed as the number of steps until the peak was reached for the first time. Although the difference in means between the small and large populations was statistically significant, the increase in lengths for the small population was less than 10% compared to the large population. **e. Path lengths for the CRP-Fis exaptation landscape.** Mean ± s.d. for large pop. 4.06 ± 0.012, mean ± s.d. for small pop. 4.07 ± 0.012; Welch Two Sample t-test, t = 11.126, df = 5119663, P-value< 2.2 x $10^{-16}$, $N_1$= $N_2$= 256x$10^4$ path lengths. **f. Path lengths for the CRP-IHF exaptation landscape with adaptive walks ending at the most accessed intermediate peak.** Mean ± s.d. for large pop. 3.42 ± 1.56, mean ± s.d. for small pop. 4.98 ± 1.55; Welch Two Sample t-test, t = 484.09, df = 1520268, P-value< 2.2 x $10^{-16}$, $N_1$= $N_2$= 256x$10^4$ path lengths. **g Path lengths for the CRP-IHF exaptation landscape with adaptive walks ending at the second most accessed intermediate peak.** Mean ± s.d. for large pop. 4.17 ± 0.080, mean ± s.d. for small pop. 4.24 ± 0.080; Welch Two Sample t-test, t = 21.659, df = 2245001, P-value< 2.2 x $10^{-16}$, $N_1$= $N_2$= 256x$10^4$ path lengths. **h Path lengths for the Fis-IHF exaptation landscape.** Mean ± s.d. for large pop. 3.93 ± 0.002, mean ± s.d. for small pop. 3.93 ± 0.002; Welch Two Sample t-test, t = 0, df = 5119998, P-value= 1, $N_1$= $N_2$= 256x$10^4$ path lengths.

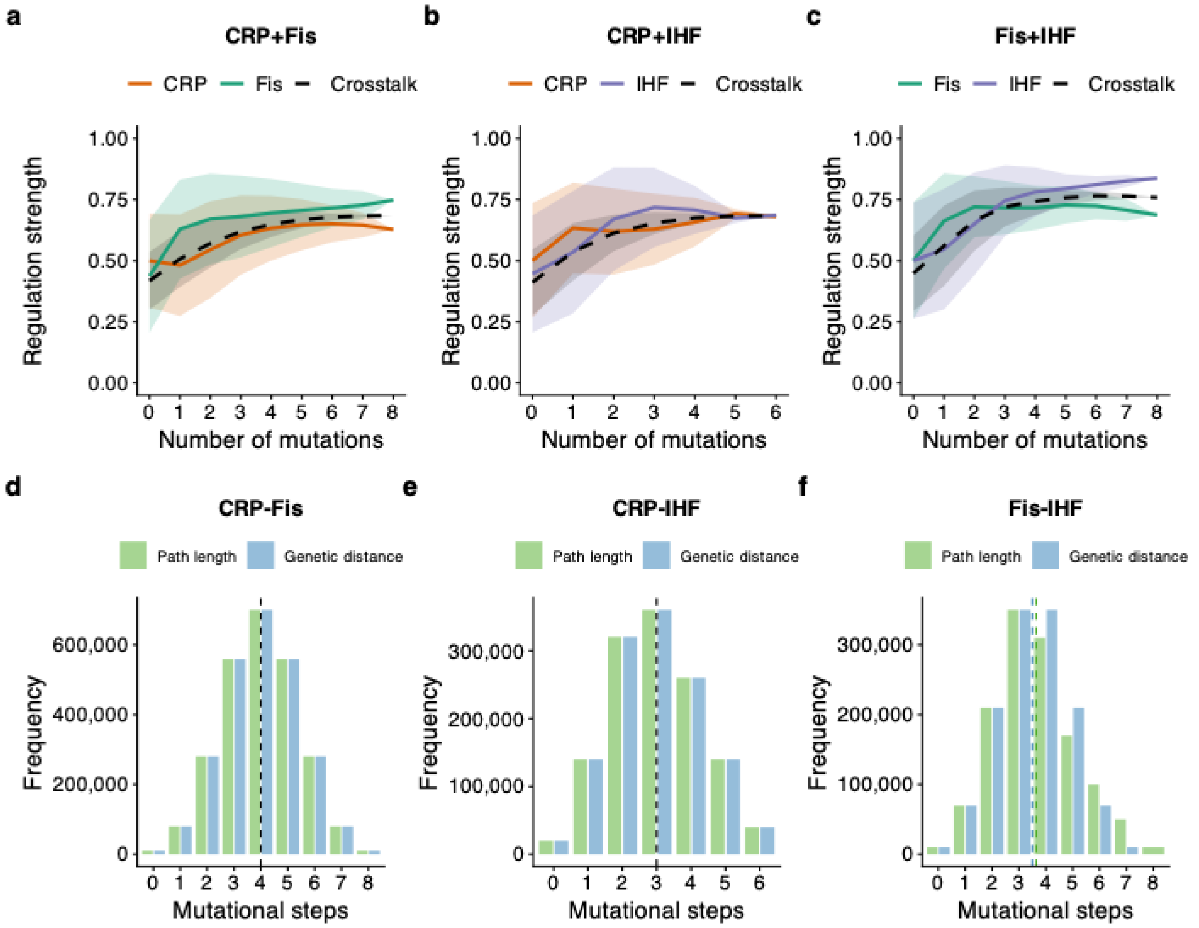


**Supplementary Figure S32. Rapid evolution of crosstalk in exaptation landscapes outside of the SSWM regime.** All data are based on $10^4$simulated greedy adaptive walks initiated from each genotype in each landscape (CRP–Fis: $256 \times 10^4$, CRP–IHF: $128 \times 10^4$, and Fis–IHF: $128 \times 10^4$). In these simulations, selection favored dual regulation, such that we defined fitness as the sum of regulation strengths for both TFs. For each landscape, a crosstalk peak is a peak genotype that maximizes simultaneous regulation by both TFs. **a-c. All adaptive walks rapidly attain a crosstalk peak.** Each plot shows how regulation strength (vertical axis) for CRP (orange), Fis (green) and IHF (purple) changes as a function of path length traversed (horizontal axis) from the initial sequence to the reached peak during adaptive walks. Crosstalk along the walks is shown by the black dashed line and is defined as the geometric mean of CRP and IHF regulation strengths. Curves represent averages over all adaptive walks (CRP-Fis: 256 x $10^4$, CRP-IHF: 128 x $10^4$; Fis-IHF: 128 x $10^4$), and shading indicates one standard deviation of regulation strengths. **a. All adaptive walks readily attain the CRP-Fis crosstalk peak.** The panel shows how the regulation strength for CRP and Fis change along the traversed path length from the initial sequence to the attained CRP-Fis peak. **b. All adaptive walks attain the CRP-IHF crosstalk peak.** Same as panel (a) but for the CRP-IHF peak. **c. All adaptive walks attain the Fis-IHF crosstalk peak.** Same as panel (a) but for the Fis-IHF peak. **d-f. Evolutionary paths to a crosstalk peak are not much longer than minimal genetic distances.** Each grouped bar chart shows the distribution of path lengths (green) and genetic distances (blue) for all pairs of starting (non-peak) and ending (peak) genotypes during $10^5$ adaptive walks. When distribution means are indistinguishable, the black dashed vertical line represents the mean of both distributions. Otherwise, the green and blue horizontal lines represent average path lengths and mutational distance, respectively. We used a t-test of the null hypothesis that the means of these distance distributions are statistically indistinguishable. **d. Distribution of**

**accessible path lengths and genetic distances to the CRP-Fis crosstalk peak.** We observed no significant difference between the means of genetic distance and path length (4±1.4, mean ± s.d., for both distance metrics, Welch Two Sample t-test t = -5.5782, df = 2,029,535, P-value= 0.7, $N_1 = N_2 = 256 \times 10^4$). **e. Distribution of accessible path lengths and genetic distances to the CRP-IHF crosstalk peak.** We observed no significant difference between the means of genetic distance and path length (3 ± 1.3, mean ± s.d., Welch Two Sample t-test t = -5.5782, df = 2,029,535, P-value= 0.67, $N_1 = N_2 = 128 \times 10^4$). **f. Distribution of accessible path lengths and genetic distances to the Fis-IHF crosstalk peak.** The difference between the means of genetic distance and path length is small, yet statistically significant (3.5 ± 1.5 and 3.6 ± 1.3, respectively, mean ± s.d., Welch Two Sample t-test, t = 197.48, df = 2,406,898, P-value< $2.2 \times 10^{-16}$, $N_1 = N_2 = 128 \times 10^4$).

1807
